# Astrocyte-targeted gene delivery of interleukin 2 specifically increases brain-resident regulatory T cell numbers and protects against pathological neuroinflammation

**DOI:** 10.1101/2022.02.28.482297

**Authors:** Lidia Yshii, Emanuela Pasciuto, Pascal Bielefeld, Loriana Mascali, Pierre Lemaitre, Marika Marino, James Dooley, Lubna Kouser, Stijn Verschoren, Vasiliki Lagou, Hannelore Kemps, Pascal Gervois, Antina de Boer, Oliver T. Burton, Jérôme Wahis, Jens Verhaert, Samar Tareen, Carlos P. Roca, Kailash Singh, Carly E. Whyte, Axelle Kerstens, Zsuzsanna Callaerts-Vegh, Suresh Poovathingal, Teresa Prezzemolo, Keimpe Wierda, Amy Dashwood, Junhua Xie, Elien Van Wonterghem, Eline Creemers, Meryem Aloulou, Willy Gsell, Oihane Abiega, Sebastian Munck, Roosmarijn E. Vandenbroucke, Annelies Bronckaers, Robin Lemmens, Bart De Strooper, Ludo Van Den Bosch, Uwe Himmelreich, Carlos P. Fitzsimons, Matthew G. Holt, Adrian Liston

## Abstract

The ability of immune-modulating biologics to prevent and reverse pathology has transformed recent clinical practice. Full utility in the neuroinflammation space, however, requires identification of both effective targets for local immune-modulation and a delivery system capable of crossing the blood-brain-barrier. The recent identification and characterization of a small population of regulatory T cells (Tregs) resident in the brain presents one such potential therapeutic target. Here we identified brain IL2 levels as a limiting factor for brain-resident Tregs. We developed a gene-delivery approach for astrocytes, with a small-molecule on-switch to allow temporal control, and enhanced production in reactive astrocytes to spatially-direct delivery to inflammatory sites. Mice with brain-specific IL2 delivery were protected from traumatic brain injury, stroke and multiple sclerosis models, without impacting the peripheral immune system. These results validate brain-specific IL2 gene-delivery as effective protection against neuroinflammation, and provide a versatile platform for delivery of diverse biologics to neuroinflammatory patients.

## Introduction

Acute central nervous system (CNS) trauma is the leading cause of death and disability for people under the age of 45 years ^1^. Although the causes of trauma are diverse, the common end result is significant neuronal damage, or neuronal loss, in the affected region ^2^. This is thought to underlie the cognitive, sensorimotor function and personality changes typically seen in patients ^1^. To date, drug treatments adopting a ‘neuro-centric’ approach have failed to deliver significant clinical benefits for the treatment of CNS injury ^1,3^, indicating that this approach is too narrow. Acute CNS injury is now recognized as triggering a multi-cellular response, involving CNS-resident immune cells (microglia and astroglia) alongside infiltration of peripheral immune cells to the brain parenchyma ^4^. While there is evidence to support a neuroprotective effect of immune activation during the initial CNS response, prolonged activation invariably becomes neurotoxic ^4-6^. The involvement of the immune system in CNS trauma counters the narrative of inevitable neurological damage, with immune-modulating biologics emerging as a key therapeutic option to control the inflammatory response. However, adoption of immune-modulating biologics in the neuroinflammatory clinical space first requires identification of biologics with effective anti-inflammatory potential in the CNS, distinct from peripheral effects, coupled with parallel development of delivery systems capable of crossing the blood-brain-barrier.

Interleukin 2 (IL2) is a high potential immune-modulating biologic, based on its capacity to support the survival and proliferation of regulatory T cells (Tregs). Tregs possess potent immunoregulatory capacity, and are common in the blood and secondary lymphoid organs, with a small population resident in the healthy CNS only recently characterized in mice and humans ^7^. While the capacity of IL2 supplementation to expand circulating Tregs and inhibit the activation and function of the immune system during neuroinflammation has been well demonstrated, these effects can be attributed to Treg function in secondary lymphoid organs. For example, more severe pathology is observed following systemic Treg-depletion in mouse models of neuroinflammation, such as the Experimental Autoimmune Encephalomyelitis (EAE) model of multiple sclerosis (MS) ^8,9^, or models of stroke ^10,11^ and traumatic brain injury (TBI) ^12^. In neuroinflammatory diseases, such as EAE, where T cells trigger the inflammatory cascade ^13^, Treg-depletion can enhance peripheral priming and infiltration of neuropathogenic T cells, regardless of any putative role for tissue-resident Tregs in the brain. Even in injury-driven neuroinflammation, such as stroke or TBI, the systemic Treg-depletion typically used to assess function also drives massive peripheral inflammation ^14,15^, with potential consequences on blood-brain barrier permeability and the nature of the leukocyte infiltrate ^12^. The involvement of CNS-resident Tregs, as opposed to peripheral-resident Tregs, in the control of neuroinflammatory pathology thus remains obscured. This unknown remains one of the key limitations in the clinical utility of IL2 in the neurology space, with the need to define potential for CNS-based impact, as opposed to systemic effects.

The functional distinction between systemic and CNS-based IL2 delivery is critical for any therapeutic exploitation in neuroinflammatory disease. In many neuroinflammatory conditions, severe systemic immunodeficiency also occurs in parallel, and thus therapeutic approaches which exacerbate peripheral immune suppression would be contraindicated. Treatments which rely on systemic IL2 provision to drive expansion of circulating Tregs as a mechanism to control CNS inflammation are, therefore, unlikely to see wide adoption in the clinic. By contrast, CNS-specific increases in IL2 could allow treatment of neuroinflammation without inducing peripheral immunosuppression. Here we demonstrate the highly efficacious control of neuroinflammation by CNS-based IL2, using a synthetic biological circuit to drive local production of IL2, with a concomitant expansion of CNS-resident Tregs achieved while leaving the peripheral immune system intact. Furthermore, we provide a solution to the biologic delivery problem for the brain, with an AAV-based therapeutic delivery system capable of providing exquisite temporal and spatial control over biologic production. The demonstrated neuroprotection in four independent neuroinflammatory models provides a clear pathway to clinical exploitation of brain-specific IL2 gene-delivery, and a platform for the delivery of diverse biologics, potentially suitable for broad classes of neuroinflammatory disease and injury.

## Results

### Transgenic production of IL2 in the brain drives Treg expansion and neuroprotection during traumatic brain injury

The potent capacity of Tregs to prevent inflammation makes increased IL2 expression (with its proven ability to expand the Treg population ^16^), an attractive therapeutic strategy for neuroinflammatory pathology. In peripheral organs the main source of IL2 is activated CD4 conventional T cells (Tconv). A negative feedback loop between Tregs and activated CD4 T cells normally limits IL2 provision, creating a stable Treg niche ^17^. In the brain, by contrast, IL2 levels are ∼10-fold lower than the serum (**Figure 1A**), with the most common IL2-producing cell type being neurons (**Figure 1B, Figure S1**). As Tregs undergo elevated apoptosis during IL2-starvation ^16^, we sought to bypass IL2 as a limiting factor using a transgenic model of IL2 autocrine expression by Tregs (**Figure 1C**). By effectively bypassing IL2 silencing, *Foxp3*^*Cre*^ *RosaIL2* mice exhibit a profound expansion of peripheral Treg numbers (**Figure 1D**). Notably, however, expansion does not occur in the brain (**Figure 1D**), with the expansionary effect of increased IL2 production in the periphery primarily observed on Tregs of the circulating phenotype, rather than the brain-resident CD69^+^ population (**Figure 1E, Figure S2A**). These results, combined with the undesirable side-effects of peripheral Treg expansion, limit the practical utility of peripheral IL2 dosing to treat neuroinflammatory pathology. Therefore, we sought to exploit the impermeable nature of the blood-brain barrier through ectopic expression of IL2 by brain cells. Using a transgenic expression system to drive cell-type-specific expression of low levels of IL2, we could activate IL2 expression in oligodendrocytes or neurons using Cre drivers (**Figure 1C,F**). While the PLP-Cre driver in oligodendrocytes gave additional systemic effects, αCamKII-Cre-mediated IL2-expression in neurons resulted in brain-specific expansion of Tregs (**Figure 1G, H**). FlowSOM cluster analysis of Tregs was used for a non-supervised clustering of dominant Treg phenotypes in the brain and periphery of the analyzed mice. Compared to Tregs from the wildtype brain, Tregs in the αCamKII^IL2^ brain were substantially depleted for the naïve cluster (CD26L^hi^CD44^low^) and highly significantly enriched in a cluster expressing multiple residential markers (CD69, CD103, ST2, KLRG1) and a CD25^hi^CD69^+^PD1^+^ cluster (**Figure 1I, Figure S2B,C**). Together, this implies enrichment in resident Tregs, with the elevation in CD25 and PD-1 known to be induced by IL2 exposure ^18^. Using imaging-based approaches, elevated numbers of Tregs were observed beyond the vasculature (**Figure 1J, Figure S3, Video S1,2**), distributed across large areas of brain (**Figure S4**). Costaining with the vascular basement membrane marker laminin α4 produced similar results, with elevated numbers of Tregs sparsely distributed across entire coronal sections in αCamKII^IL2^ mice (**Figure S5, Supplementary Resource 1**). By flow cytometry, increases in Treg numbers were observed across the brain regions assessed (**Figure S6**). Only minor changes were observed in the non-Treg brain-resident leukocyte populations of αCamKII^IL2^ mice, with limited shifts in population frequency and marker expression (**Figure S2D-K**). Single-cell sequencing demonstrated that, within the CD4 T cell compartment, only the residential Treg cluster (enriched for CD69 expression) was affected in frequency by brain-IL2 provision (**Figure 1K-N, Figure S7A-C**). Aside from this numerical expansion, no major transcriptional alterations were observed in the Treg population (**Figure S7D**). Brain Treg expansion did not alter the electrophysiology of neurons (**Figure S8**), and resulted in no major adverse behavioral changes (**Figure S9**) or excess mortality (aged to 18 months with 90% survival vs 93% in the control littermates. n=11, 15).

**Figure 1.**
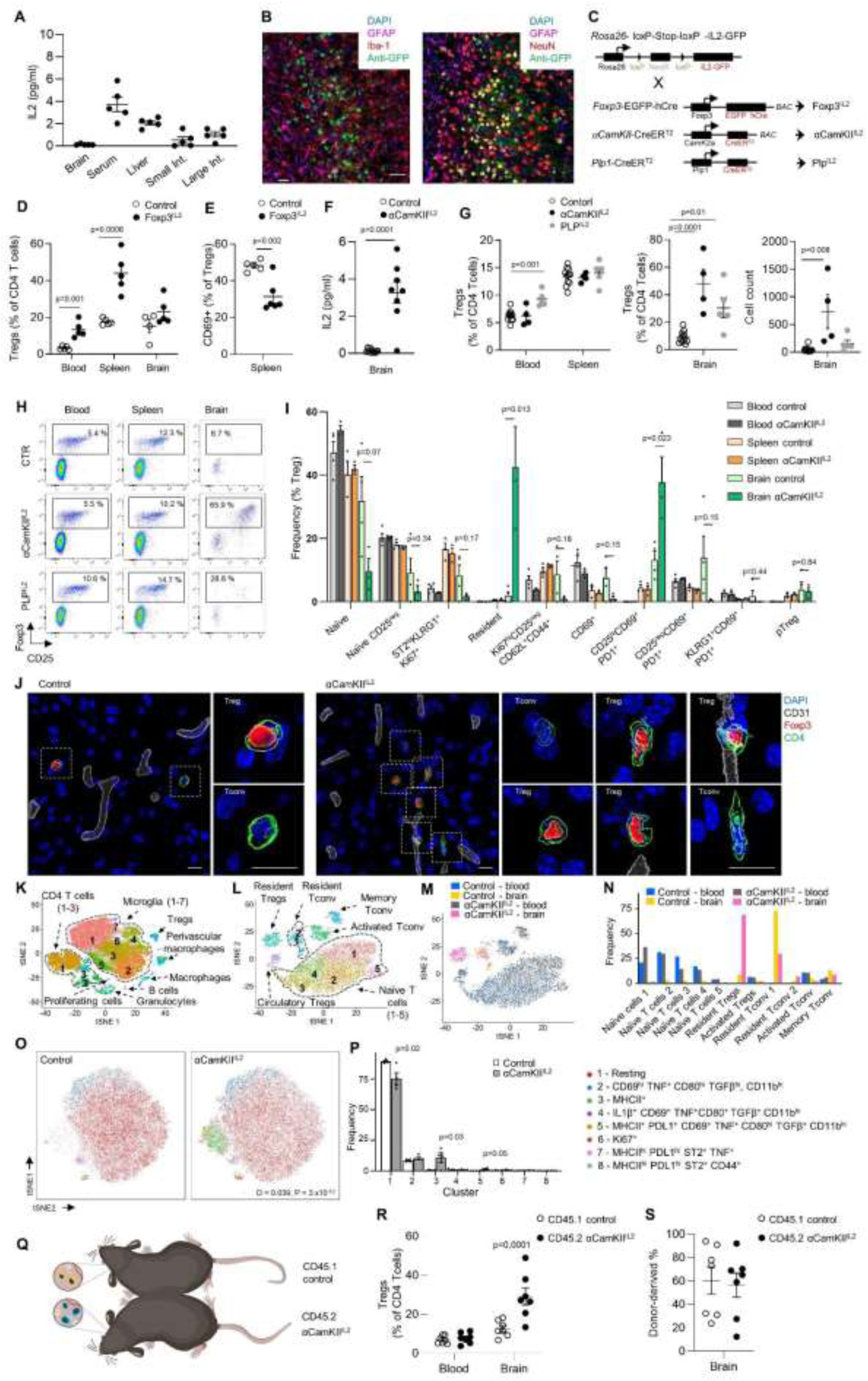
Local expression of IL2 drives a brain-specific expansion of Tregs. **(A)** IL2 levels detected by ELISA in the brain, serum, liver, small intestine and large intestine (n=5/group). **(B)** Representative confocal images from the midbrain of IL2^GFP^ reporter mice, stained using antibodies against the major CNS cell types: IL2-producing cells were mainly NeuN+ neurons. Scale bar, 50 µm. **(C)** Schematic of transgenic systems for IL2 expression initiated by *Foxp3, Plp1* or *αCamKII* promoters. **(D)** Frequency of Foxp3^+^ cells within CD4 T cells, and **(E)** CD69^+^ cells within Tregs, in wildtype and Foxp3^IL2^ mice (n = 5, 5), measured by flow cytometry. **(F)** IL2 levels detected by ELISA in the brain of wildtype and αCamKII^IL2^ mice (n=11,8). **(G)** Frequency of Foxp3^+^ cells within CD4 T cells in the blood, spleen and brain for control, PLP1^IL2^ and αCamKII^IL2^ mice, at least 4 weeks post-tamoxifen treatment (n = 8, 5, 4), measured by flow cytometry. Absolute number of Foxp3^+^ cells in the brain. **(H)** Representative staining for panel (G) indicating Foxp3 and CD25 co-expression. **(I)** Healthy perfused mouse brains, spleen and blood from wildtype and αCamKII^IL2^ mice were compared by high-dimensional flow cytometry (n = 4, 3). Subpopulations were defined by FlowSOM clusters and annotated based on key markers (CD62L, CD44, CD103, CD69, CD25, PD-1, Nrp1, ICOS, KLRG1, ST2, Ki67, Helios, T-bet, CTLA4). **(J)** Surface rendering of confocal images. Representative image of Treg in the mid-brain in wildtype and αCamKII^IL2^ mice. Scale bar, 10 µm. Pre-rendered images in Figure S3. **(K)** tSNE projection of 28,590 sorted cells (20,021 microglia, 6332 T cells and 2237 non-target leukocyte populations) from wildtype or αCamKII^IL2^ mice. Clusters were annotated based on the expression of key lineage markers. Non-target cell types were annotated as labelled, prior to removal. **(L)** tSNE projection of the CD4 T conventional and Tregs clusters. Immune populations were annotated based on key lineage markers, or **(M)** based on origin from wildtype or αCamKII^IL2^ mice, **(N)** with cluster quantification. **(O)** Healthy perfused mouse brains from wildtype and αCamKII^IL2^ mice were compared by high-dimensional flow cytometry (n = 4, 4; 64,927 cells plotted). tSNE of microglia built on key markers (CD64, MHCII, TGFβ, LAMP1, CD44, CD69, PDL1, ST2, Ki67, CD80, IL1β, CX3CR1, CD45, TNF), **(P)** with cluster quantification. **(Q)** CD45.1 mice were parabiosed to tamoxifen-treated CD45.2 αCamKII^IL2^ mice. **(R)** Percentage of Tregs with the CD4 T cell population in the blood and perfused brain of parabiotic pairs (n=7). **(S)** Proportion of Tregs originating from the CD45.1 or CD45.2 donor in the perfused brain of parabiotic pairs.

In order to investigate the transcriptional changes induced in the targeted population, namely CD4 T cells and microglia (the primary immunological cell type in the brain), we turned to single-cell sequencing. Bulk CD4 T cells and CD11b^+^ myeloid cells were sorted, sequencing libraries prepared and sequenced. Reads were then mapped to the mouse genome and transcripts identified, followed by tSNE projection with subsequent identification of Tregs and microglia, based on the expression of canonical markers (**Figure S7**). Single-cell sequencing identified an increase in MHCII-related gene expression in microglia, potentially enabling enhanced interaction between brain Tregs and microglia, but otherwise no major transcriptional changes were detected (**Figure S7**). Increased MHCII expression was validated by flow cytometry, where ∼15% of microglia expressed MHCII in αCamKII^IL2^ mice, with the microglia otherwise normal and not expressing inflammatory or activation markers (**Figure 1O, P** and **Figure S10**). The exception was PD-L1, which was increased on microglia in αCamKII^IL2^ mice (**Figure S10**), corresponding with the increase in PD-1 expression in brain Tregs (**Figure S3**). As PD-1 engagement on Tregs protects the cells from apoptosis ^18^, the upregulation of these interaction partners may contribute to the Treg expansion observed following IL2 upregulation. Together, these results demonstrate a rare example of the blood-brain barrier working in favor of an intervention, with brain-specific expression of IL2 resulting in a local expansion of the resident Treg population without off-target impacts on peripheral Tregs.

We previously characterized the brain-resident Treg population as a semi-transient migratory population ^7^. Using parabiotic experiments, brain Treg seeding was determined to largely occur through activated Tregs in the blood, of which the majority rapidly die or leave within days, while ∼5% gain a residential phenotype and dwell for weeks ^7^. To determine whether brain-delivered IL2 primarily works through enabling more efficient seeding of activated Tregs entering the CNS, or through the selective expansion (or retention) of Tregs already-resident, we performed parabiosis on αCamKII^IL2^ mice. αCamKII^IL2^ mice, or wildtype control mice, were treated with tamoxifen to initiate IL2 expression and then parabiosed to congenically-disparate mice (**Figure 1Q**). This creates a system where a single circulatory system feeds into two brains, one (wildtype) with normal, limiting, levels of IL2, and the other (αCamKII^IL2^) with IL2 levels elevated to that of the serum. Following equilibration of circulating cells, we assessed the Treg compartment of the brain of parabiotic pairs by flow cytometry. First, we found that αCamKII^IL2^ mice, but not controls or parabiotic pairs, exhibited an increase in brain-resident Tregs (**Figure 1R**). This demonstrates that the expansion is restricted to the supplemented brain niche, and is not transmitted via circulatory factors. Second, in both wildtype brains and αCamKII^IL2^ brains, the brain-resident Treg population demonstrated equivalent representation of host and donor Tregs (**Figure 1S**). This result was consistent with our previous characterization of brain Tregs as transient residents, fed by a consistent incoming flow ^7^. The equivalent expansion of host and donor Tregs in the IL2-supplemented brain suggests that brain-specific IL2 levels expand the total niche size available for brain Tregs, without altering the homeostatic migratory kinetics that allows this niche to be filled by incoming blood-derived cells. Together, these results demonstrate that brain-specific IL2 levels are the limiting factor in controlling the population kinetics of incoming Tregs, as a niche functionally distinct from the peripheral system.

Having identified that brain and peripheral Treg populations are reliant on compartmentalized IL2 niches, we next sought to determine whether supplementing the paucity in brain IL2 could alleviate pathological neuroinflammation-driven pathology. While IL2 and Treg therapy have been reported in other contexts ^19-22^, these approaches expanded the systemic Treg population, and thus the effects observed cannot be definitively attributed to local CNS effects. To test whether brain-specific supplementation of IL2 could influence neuroinjury, independent of systemic immunosuppression, we used a controlled cortical impact model of TBI to deliver an acute insult. In wildtype mice, this injury typically leads to wide-spread neurodegeneration 14 days after injury (**Figure S11**), concomitant with astrogliosis and microgliosis, indicative of an inflammation-mediated mechanism. αCamKII^IL2^ mice, by contrast, exhibited a high degree of protection against damage at the inflammatory site (**Figure 2A**), with reduced lesion size and partial preservation of neuronal tissue (**Figure 2B,C**). Microgliosis was not observed in the post-TBI cortex, with increased Iba1 intensity in αCamKII^IL2^ mice in the post-TBI striatum (**Figure 2D**). Astrogliosis was unaffected by the mouse genotype (**Figure 2E**). Despite the partial anatomical preservation in αCamKII^IL2^ mice, the leukocyte influx in the brain remained relatively unchanged, apart from the increased Treg frequency present already prior to TBI (**Figure 2F,G, Figure S12**). These results demonstrate proof-of-principle for local brain-specific IL2 production as a potent suppressor of neuroinflammation-induced pathology.

**Figure 2.**
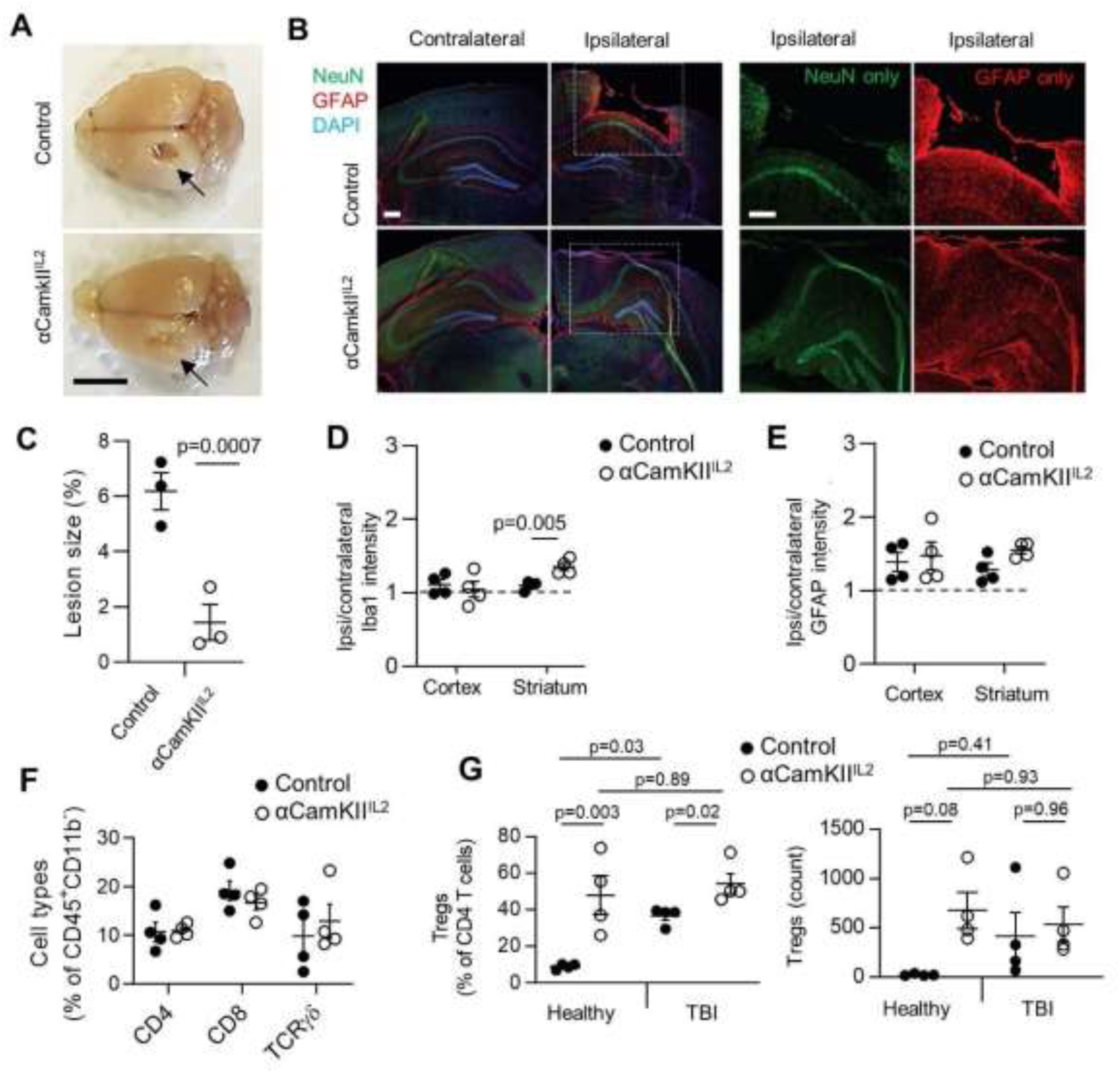
Protection from neuroinflammation following brain-specific expansion of Tregs. **(A)** Control littermates and αCamKII^IL2^ mice were tamoxifen treated at 6 weeks and controlled cortical impacts to induce moderate TBI were given at 12 weeks. Mice were examined 15 days post-TBI (n = 3, 3). Representative photos illustrating damage to the surface of the brain at the injury site. Arrow, site of impact. Scale bar, 0.5 cm. **(B)** Representative immunofluorescence staining of the cortical tissue 14 days after cortical impact. GFAP (astrocytes), NeuN (neurons), DAPI (nuclei). Scale bars, 50 µm. **(C)** Lesioned area, shown as percentage of the entire hemisphere (n = 3, 3). **(D)** Relative Iba1 and **(E)** GFAP expression levels in the cortex and striatum (ratio of expression in ipsilateral vs contralateral hemispheres). **(F)** TBI-induced perfused brains from wildtype and αCamKII^IL2^ mice were compared at 15 days post-TBI by high-dimensional flow cytometry (n=4,4). Frequency of CD4, CD8 and γδ T cells within CD45^+^CD11b^-^ cells. **(G)** TBI-induced perfused brains from wildtype and αCamKII^IL2^ mice were compared prior to TBI, or at 15 days post-TBI by high-dimensional flow cytometry (n=4/group). Frequency of Tregs within CD4^+^ T cells (left) and absolute number of Tregs (right).

### A ‘dual lock’ IL2 gene-delivery system drives expansion of the brain-resident Treg population

We next sought to translate our transgenic system to a gene-delivery approach with clinical potential. To improve the biological properties of the IL2 micro-targeting while avoiding neurons as targets, we shifted IL2 production to astrocytes, using the *GFAP* promoter to drive IL2 expression. *GFAP*-mediated IL2 expression has several theoretical advantages: 1) astrocytes are efficient secretory cells, widely distributed across the brain; 2) astrocytes are well represented in the spinal cord, potentially expanding the utility of this approach to spinal cord-dependent diseases; 3) localized astrogliosis occurs during neuroinflammatory events such as TBI (**Figure 3A**), and 4) the *GFAP* promoter is more active during astrogliosis, concentrating cargo production in the inflamed region of the brain (**Figure 3B**). To take advantage of astrocytes as an expression system, we designed a ‘dual-lock’ delivery system for IL2, using AAV-PHP.B combined with a modified *GFAP* promoter capable of driving robust expression widely in astrocytes (**Figure 3C**). The system combines the enhanced CNS gene delivery (brain and spinal cord) seen with the PHP.B capsid following systemic delivery ^23,24^, with the secondary specificity of using a modified endogenous promoter restricted to astrocytes (*GFAP*), such that (peripheral) off-target transduction would be unable to drive cargo expression. This ‘dual-lock’ system results in astrocyte-driven cargo expression, as assessed by GFP expression using of AAV-PHP.B.*GFAP*-GFP (PHP.*GFAP*-GFP) (**Figure 3C**). Both S100β^+^GFAP^+^ and S100β^+^GFAP^-^ cortical astrocytes were able to express the transgene, with limited coexpression observed with neurons, microglia, oligodendrocytes or NG2^+^ cells (**Figure 3C,D**). Elevated expression of the cargo was observed around the TBI impact site (**Figure 3E**), demonstrating that the upregulation seen in endogenous promoter activity (**Figure 3B**) was faithfully recapitulated by the exogenous *GFAP* promoter in the vector (**Figure 3B**).

**Figure 3.**
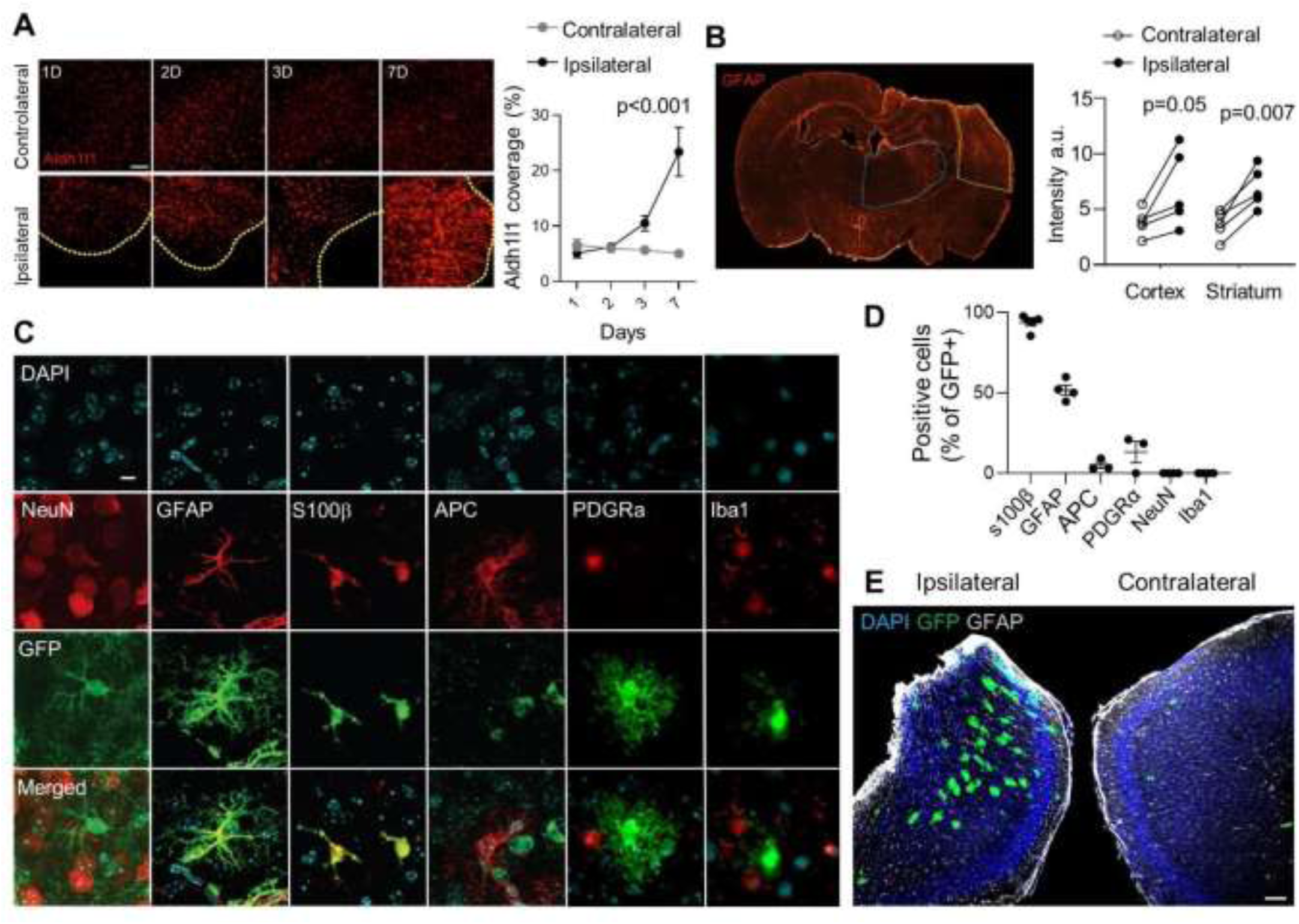
Synthetic delivery to the brain via a ‘dual lock’ gene delivery system. **(A)** Wildtype mice were given controlled cortical impacts to induce moderate TBI and examined at 1, 2, 3 and 7 days post-TBI (n = 5). Representative images (left) and quantification (right) of astrocyte coverage in the cortex adjacent to the lesion (delineated in yellow) or corresponding contralateral cortical area, ascertained via Aldh1l1 immunostaining (n=3). Scale bar 100 µm. **(B)** Representative staining (left) and quantified expression (right) of GFAP in the cortex (yellow) and striatum (blue), 14 days post-TBI (n=5), with quantification. Scale bar, 50 µm. **(C)** The *GFAP* promoter restricts gene expression (as assessed using GFP scoring) to astrocytes in adult mouse brain, based on characteristic cell morphology and by immunostaining for the astrocyte specific markers, GFAP and S100β. Off-target expression was not detected when slices were counter-stained for NeuN (neurons), APC (oligodendrocytes), PDGFRα (NG2^+^ cells) and Iba1 (microglia). Scale bar, 20 µm. Data are representative images seen in 3 slices from each of 3 independent mice receiving a PHP.*GFAP*-GFP (control) vector. **(D)** Quantification of GFP colocalisation with cell lineage markers in PHP.*GFAP*-GFP-treated mice. **(E)** Wildtype mice were given controlled cortical impacts to induce moderate TBI, treated with PHP.*GFAP*-GFP and examined at 14 days post-TBI/treatment. Representative image of GFP production in the ipsilateral region surrounding the impact site or the corresponding contralateral cortical area. Scale bar, 100 µm.

Having validated the AAV-PHP.B.*GFAP* system for brain-specific expression, we sought to determine whether it could be applied to IL2 delivery with similar results to the αCamKII^IL2^ system. Using AAV-PHP.B.*GFAP*-IL2 (PHP.*GFAP*-IL2) delivery, we observed a 3-fold increase in brain IL2 production (**Figure 4A**), observed over the course of 14 days (**Figure 4B**). The increase in brain IL2 concentrations was paralleled by an increase in brain Treg frequency (**Figure 4C**) and absolute number (**Figure 4D**). The increase in brain Tregs was not observed in the superficial or deep cervical lymph nodes (**Figure S13A-D**), but was mirrored in the pia mater (**Figure S13E,F**). The expansion of the Treg population was dose-dependent (**Figure 4E,F**), and restricted to the brain, without expansion of Tregs in the blood, spleen or other peripheral tissues (**Figure 4G, Figure S13G-H**). The expanded Tregs were of the CD69^+^ residential phenotype (**Figure 4H**), and were observed in the brain tissue beyond the vasculature (**Figure 4I, Figure S14** and **Video S3,4**). Critically, no major off-target effects were observed, either in terms of peripheral Treg expansion (**Figure 4H**) or the population size and phenotype of non-Tregs in the brain (**Figure S15**). PHP.*GFAP*-IL2 treatment was well tolerated, with no excess mortality through 300+ days of monitoring (n=14). Both neuronal function (measured electrophysiologically) and astrocyte function (Ca^2+^ imaging) were unaffected by PHP.*GFAP*-IL2 treatment (**Figure S16**) and no behavioral abnormalities were observed in treated mice (**Figure S17**). The blood-brain barrier remained histologically and functionally intact following gene delivery (**Figure S18**). A similar degree of Treg expansion was observed with neuronal-derived IL2, following treatment with PHP.*CamKII*-IL2 treatment (**Figure S19**), demonstrating source-independent effects of IL2 on Treg expansion. Therefore, our ‘dual-lock’ PHP.*GFAP*-IL2 approach combines the key desired attributes for treatment of neuroinflammatory pathology: increased IL2 production in the brain, a rapid and sustained effect, and a restricted locus of action to avoid generalized immune suppression.

**Figure 4.**
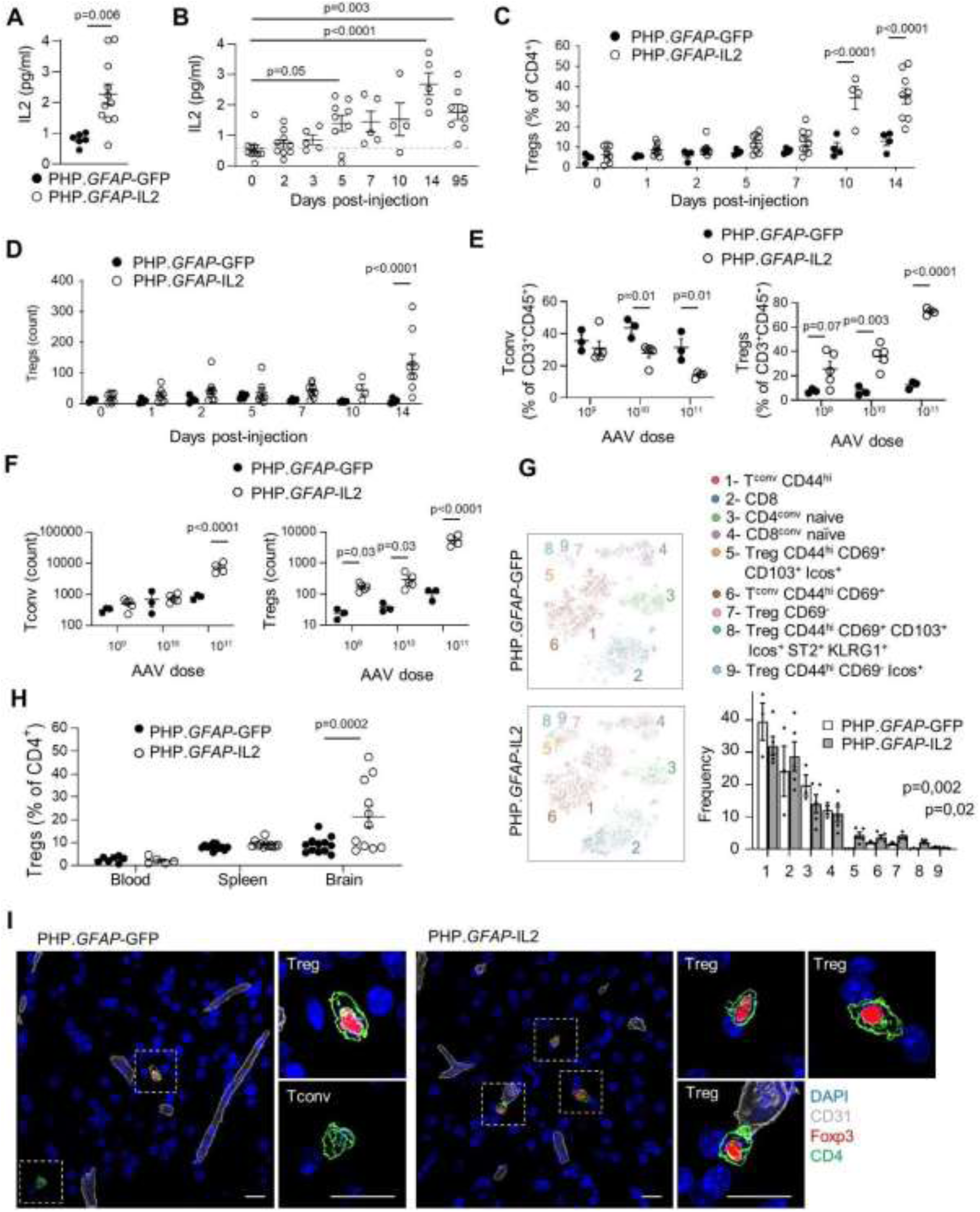
‘Dual lock’ delivery of IL2 to the brain expands local regulatory T cells. **(A)** IL2 levels detected by ELISA in the brain of wildtype mice, 14 days post-treatment with PHP.*GFAP*-GFP or PHP.*GFAP*-IL2 (n=11/group). **(B)** Time-course of IL2 levels in the brain of mice treated with PHP.*GFAP*-IL2 (n= 10, 9, 5, 9, 5, 4, 8). **(C)** Time-course of Treg expansion, as a proportion of CD4 T cells or **(D)** absolute number, in the brain of mice treated with PHP.*GFAP*-GFP or PHP.*GFAP*-IL2 (n=4, 9). **(E)** Wildtype mice were administered 1×10^9^, 1×10^10^ or 1×10^11^ vector genomes (total dose) of PHP.*GFAP*-GFP or PHP.*GFAP*-IL2 by intravenous injection and assessed for the frequency or **(F)** absolute number of conventional T cells (left) and Tregs (right) in the perfused brain 14 days after treatment (n=3-5/group). **(G)** Blood, spleen and perfused mouse brain from PHP.*GFAP*-GFP and PHP.*GFAP*-IL2-treated mice were compared by high-dimensional flow cytometry for Treg numbers (n=11-12/group). **(H)** tSNE of CD45^+^CD11b^-^CD19^-^CD3^+^ T cells built on key markers (CD4, CD8, Foxp3, CD62L, CD44, CD103, CD69, CD25, PD-1, Nrp1, ICOS, KLRG1, ST2, Ki67, Helios, CTLA4) from perfused brain. Colors indicate annotated FlowSOM clusters; results are quantified in the bar graph. Mean ± SEM. **(I)** Representative images (surface rendered confocal sections) of Tregs in the mid-brain of PHP.*GFAP*-GFP and PHP.*GFAP*-IL2 -treated mice. Scale bar, 10 µm.

### Brain-specific IL2 gene-delivery provides preventative and curative neuroprotection during neuroinflammatory injury and disease

To test the therapeutic potential of the dual-lock PHP.*GFAP*-IL2, we treated mice with a control PHP.B (encoding GFP) or PHP.*GFAP*-IL2 and then exposed them to TBI. The strong protective effect was apparent at a gross morphological level (**Figure 5A**), with reduced loss of cortical tissue at 14 days post-injury, as shown by histology (**Figure 5B,C**) and MRI (**Figure 5D**). A trend towards reduced microgliosis was observed in PHP.*GFAP*-IL2-treated mice, and astrogliosis was significantly reduced in the damaged cortex (**Figure 5E**). The neuroprotective effect was also observed at the behavioral level, where the poor performance of post-TBI mice in the Morris Water Maze and Novel Object Recognition behavioral tests was completely reversed in PHP.*GFAP*-IL2-treated mice (**Figure 5F-H**). These results validate the neuroprotective potential of synthetic IL2 delivery that we observed in the transgenic model, with the dual-lock gene delivery system being potentially translatable to the patients.

**Figure 5.**
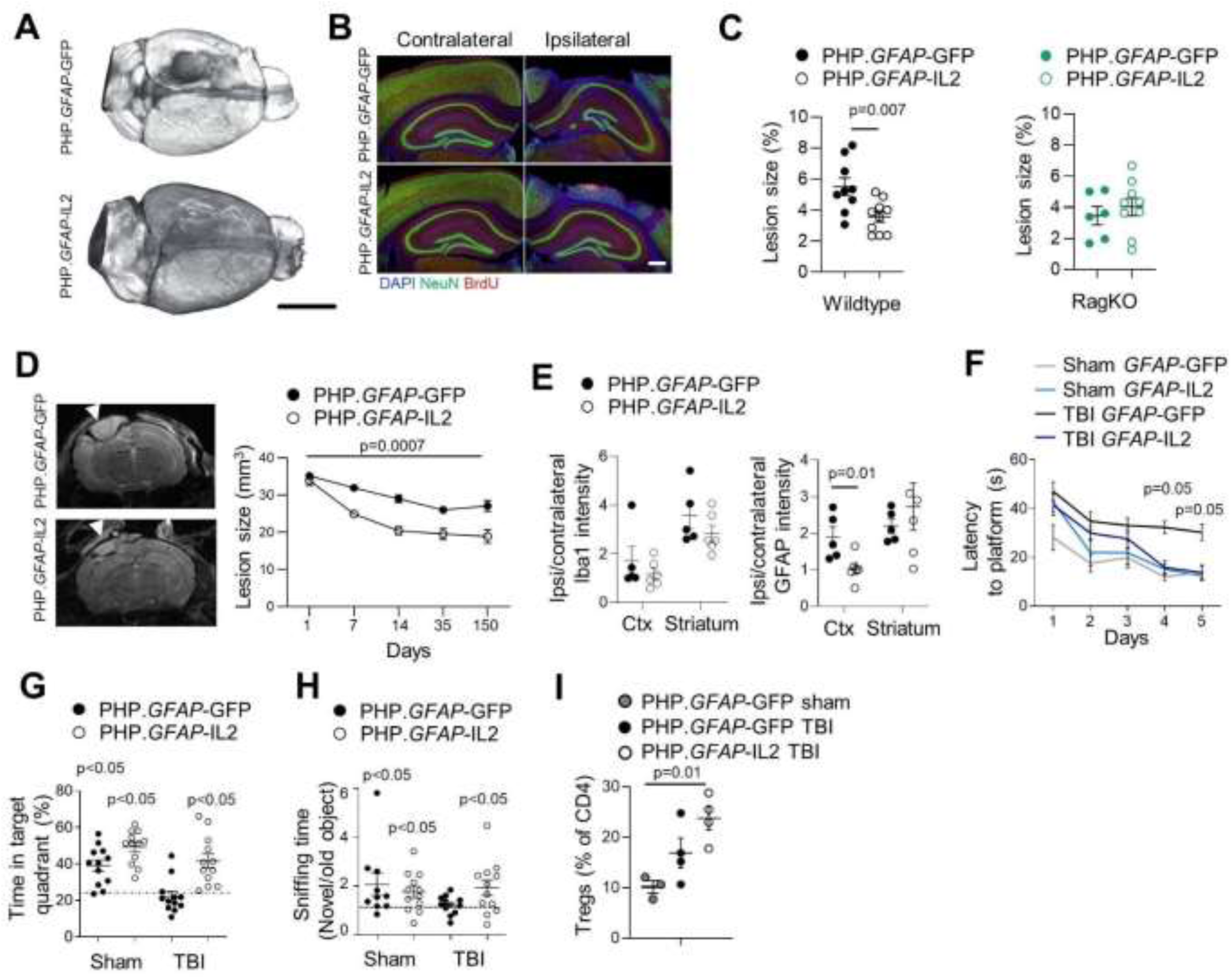
Synthetic expansion of brain regulatory T cells effectively prevents neurological damage during traumatic brain injury. **(A)** Wildtype mice, treated with PHP.*GFAP*-IL2 (or PHP.*GFAP*-GFP control vector) on day -14 were given controlled cortical impacts to induce moderate TBI and examined at 14 days post-TBI (n = 5, 6). Macroscopic damage to the surface of the brain at the injury site, representative tomography. Scale bar, 0.5 cm. **(B)** Representative immunofluorescence staining of the cortical tissue 14 days after controlled cortical impact surgery (n = 5, 6). NeuN (neurons), BrdU (proliferation), DAPI (nuclei). Scale bar, 50 µm. **(C)** Quantification of cortical area lost due to lesion 14 days post-TBI in wildtype mice (left), treated with PHP.*GFAP*-IL2 (or control vector) on day -14 (n=9,10), or RagKO mice (right), treated with PHP.*GFAP*-IL2 (or control vector). **(D**) Representative MRI and MRI-based quantification of lesion size in PHP.*GFAP*-GFP or PHP.*GFAP*-IL2-treated mice on days 1, 7, 14, 35, 150 post-TBI (control n = 16, 16, 12, 11, 10; IL2 n = 16, 16, 16, 12, 9). Impact site indicated by arrow. **(E)** Relative Iba1 and GFAP expression levels in the cortex and striatum (ratio of expression in ipsilateral vs contralateral hemispheres). **(F)** Latency to find a hidden platform in the Morris Water Maze test during acquisition learning, for PHP.*GFAP*-GFP and PHP.*GFAP*-IL2-treated mice, with and without (sham) TBI. P-value for TBI PHP.*GFAP*-GFP against TBI PHP.*GFAP*-IL2. **(G)** Percentage of total time spent in the target quadrant during the probe trial. P-value, one-sample t-test against chance level. **(H)** Ratio of time spent exploring a novel object over an old object during day 2 of the Novel Object Recognition paradigm. P-value, one-sample t-test against chance level. **(I)** Mice treated with PHP.*GFAP*-GFP control or PHP.*GFAP*-IL2 (day -14) were given controlled cortical impacts to induce moderate TBI and examined at 15 days post-TBI (n = 3, 4, 4); a sham TBI was included in the PHP.*GFAP*-GFP group. Perfused brains from sham, TBI and PHP.*GFAP*-IL2-treated TBI mice were compared by high-dimensional flow cytometry for frequency of Tregs as a proportion of CD4 T cells.

In order to test the mechanism of action, we first performed TBI and PHP.*GFAP*-IL2 treatment on RagKO mice, which do not show an adaptive immune response. Compared to wildtype mice subjected to TBI, lesions in RagKO mice following TBI were generally smaller (**Figure 5C**), consistent with a partial role for adaptive immunity in TBI pathology. When given PHP.*GFAP*-IL2, RagKO mice did not exhibit any beneficial effect from the brain-targeted IL2 expression (**Figure 5C**). These results formally exclude mechanisms of IL2 action based on direct effects on the neuronal or glial compartments, and strongly support the observed Treg expansion as the mechanistic driver of neuroprotection. The Treg effect was, however, likely mediated through modification of the local environment, with little change observed to the inflammatory influx (**Figure S20**), other than the increase in brain Tregs (**Figure 5I**) with enhanced amphiregulin production (**Figure S20**). Treatment did, however, prevent microgliosis formation, with the increase in microglia following TBI abrogated in PHP.*GFAP*-IL2-treated mice (**Figure S20A**). We therefore performed single-cell transcriptomics analysis of T cells and microglia from treated and control mice, given TBI or sham surgery. Within the identified T cell clusters (**Figure S21**), the only population shifted in frequency was Tregs, with increases in PHP.*GFAP*-IL2-treated mice, both in sham and TBI animals (**Figure 6A**). The transcriptome of expanded Tregs was largely conserved, with increases in the IL2 receptor components *Il2ra* and *Il2rb* and the anti-apoptotic gene *Bcl2* (**Figure 6B**), suggesting efficacy via numerical increase rather than the upregulation of unique effector molecules. Microglia were clustered into two super-clusters, one representing homeostatic microglia and one representing activated microglia (**Figure 6C,D, Figure S21**). While activated microglia were sharply elevated following TBI, the proportion of microglia in activated states were equivalent in IL2-treated and control mice (**Figure 6E**). Upregulation of MHCII in IL2-treated activated microglia was the prominent transcriptional change observed (**Figure 6F, Figure S21**). Notably, MHCII^high^ microglia from IL2-treated mice formed a distinct subcluster within the activated microglia cluster (**Figure 6C,D**). While the main activated subcluster expressed the classical disease-associated microglia (DAM) transcriptional profile ^25^, the MHCII^high^ subcluster had upregulated transcription of MHCII components, but were otherwise low expressors of inflammatory mediators (**Figure 6G, Figure S21**). IL2-treatment was associated with a skewing of activated microglia away from the classical DAM phenotype and towards the atypical MHCII^high^ phenotype (**Figure 6E**). As MHCII^high^ microglia accumulated at the lesion border (**Figure 6H**), this unique population may serve as a buffer against neurotoxic inflammation.

**Figure 6.**
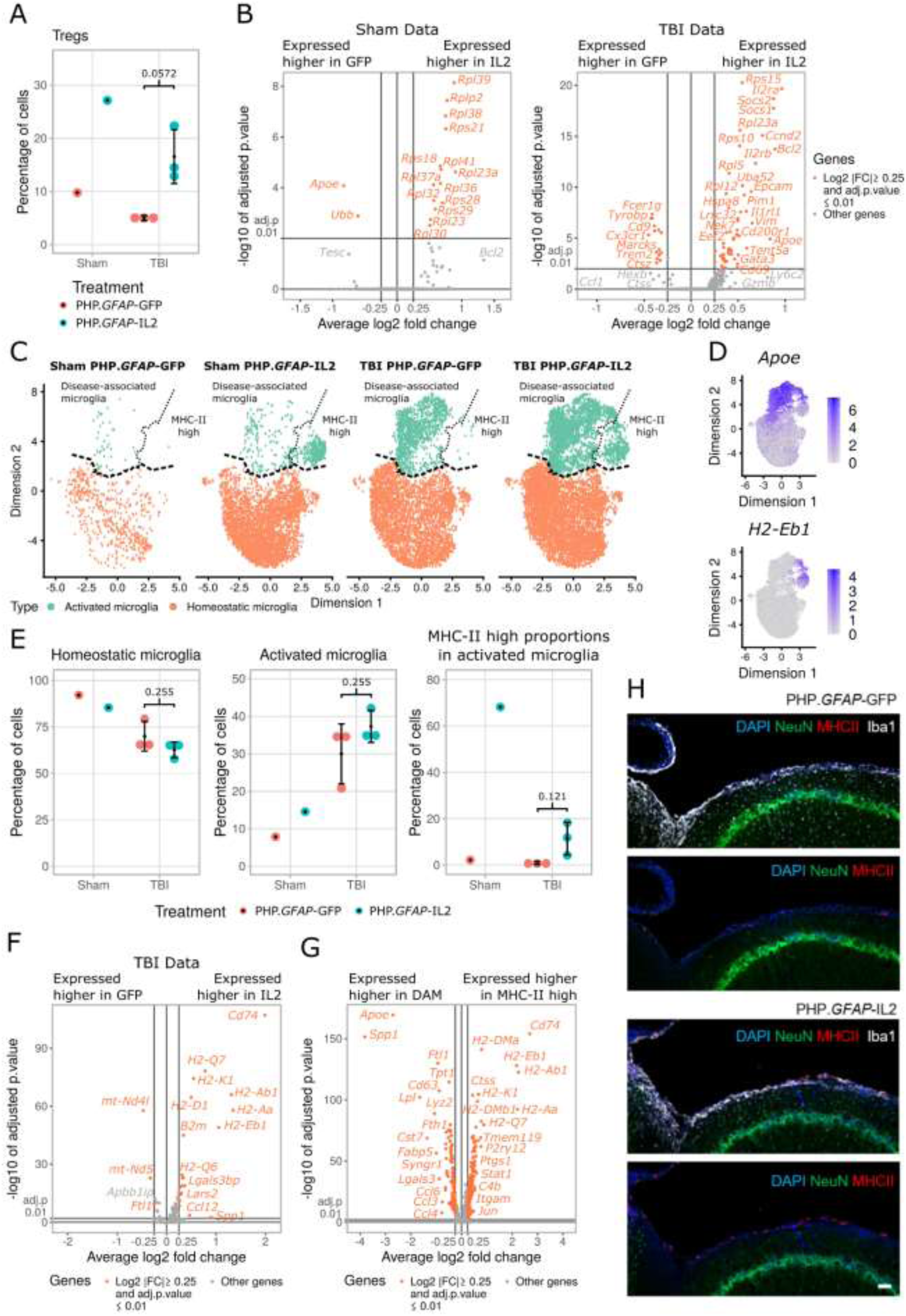
Brain-specific delivery of IL2 drives microglial transcriptional divergence during TBI. Wildtype mice, treated with PHP.*GFAP*-IL2 (or PHP.*GFAP*-GFP control vector) on day -14 were subjected to controlled cortical impacts to induce moderate TBI or sham surgery. 14 days post-TBI, T cells and microglia were sorted from the ipsilateral hemisphere of the perfused brains for 10x single-cell transcriptomics. **(A)** T cells were clustered and annotated, based on markers defined in Supplementary Figure 21A, B. Quantification of the Treg cluster based on group. **(B)** Volcano plot showing differential gene expression in the Treg cluster between PHP.*GFAP*-GFP- and PHP.*GFAP*-IL2-treated mice, for sham (left) and TBI (right). **(C)** Microglia UMAP representation, showing the location of cells per cluster for each treatment group. Cluster annotation based on expression of **(D)** ApoE and H2-Eb1 expression; expression of additional inflammatory markers is shown in Supplementary Figure 21E. **(E)** Quantification of the homeostatic and activated microglia clusters, and, within the activated microglia cluster, the relative contribution of the DAM and MHCII^high^ subclusters. **(F)** Volcano plot showing differential gene expression in the activated microglia cluster between PHP.*GFAP*-GFP- and PHP.*GFAP*-IL2-treated mice, for TBI. **(G)** Volcano plot showing differential gene expression, independent of treatment group, for the DAM vs MHCII^high^ subclusters. **(H)** Representative immunofluorescence staining of the cortical tissue at 14 days post-TBI (n = 5, 6). NeuN, MHCII, DAPI and (top only) Iba1. Scale bar, 50 µm.

To test the versatility of the dual-lock system, we extended these findings to other neuroinflammatory pathologies. We tested mouse models of ischemic stroke, as this is the most common cause of fatal neurological pathology in humans. Mice were pre-treated with PHP.*GFAP*-IL2 prior to the induction of a distal middle cerebral artery occlusion (dMCAO). Compared to control treated mice, PHP.*GFAP*-IL2-treated mice developed a macroscopically smaller lesion (**Figure 7A**), with a ∼50% reduction in the histological lesion size at day 14 (**Figure 7B**) and reduced lesion sizes detected by MRI from day 1 to 14 post-induction (**Figure 7C**). In the photothrombotic stroke model, mice pre-treated with PHP.*GFAP*-IL2 exhibited reduced macroscopic damage (**Figure 7D)**, and a ∼30% reduction in infarct size as quantified by combined scar tissue and ischemic tissue identification (**Figure 7E**). In both the dMCAO (**Figure S22**) and photothrombotic models (**Figure S23**), analysis of the immunological compartment indicated elevated numbers of T cells (CD4, CD8 and Treg) in the brain following injury, with no notable effect of treatment on this elevation. We then turned to the EAE model of MS. Using pre-treatment of mice, PHP.*GFAP*-IL2 resulted in a lower incidence, reduced clinical time-course and a lower cumulative clinical score in the MOG model (**Figure 7F**). As with the stroke models, the immunological composition of the brain was largely unchanged at the time-points assessed, including the number of Tregs present, although elevated production of anti-inflammatory cytokines was observed (**Figure S24**). The equilibration of brain-resident Tregs in control and treated mice, 4+ weeks after initial PHP.*GFAP*-IL2 treatment, across both stroke and EAE, suggests either a maximal duration of efficacy following a single AAV dose, or a confounding effect of pathology-derived Tregs obscuring the treatment-derived Treg increase. Further, the observed efficacy in these models, despite Treg normalization at end-stage, suggests that either the major effects were implemented during earlier phases of disease, or that immune-modulation of a local population, such as microglia, extends beyond the period of elevated Treg numbers. Despite the questions on treatment kinetics raised, the efficacy of treatment in these models demonstrates the extended therapeutic potential of PHP.*GFAP*-IL2 beyond TBI to other neuroinflammatory pathologies.

**Figure 7.**
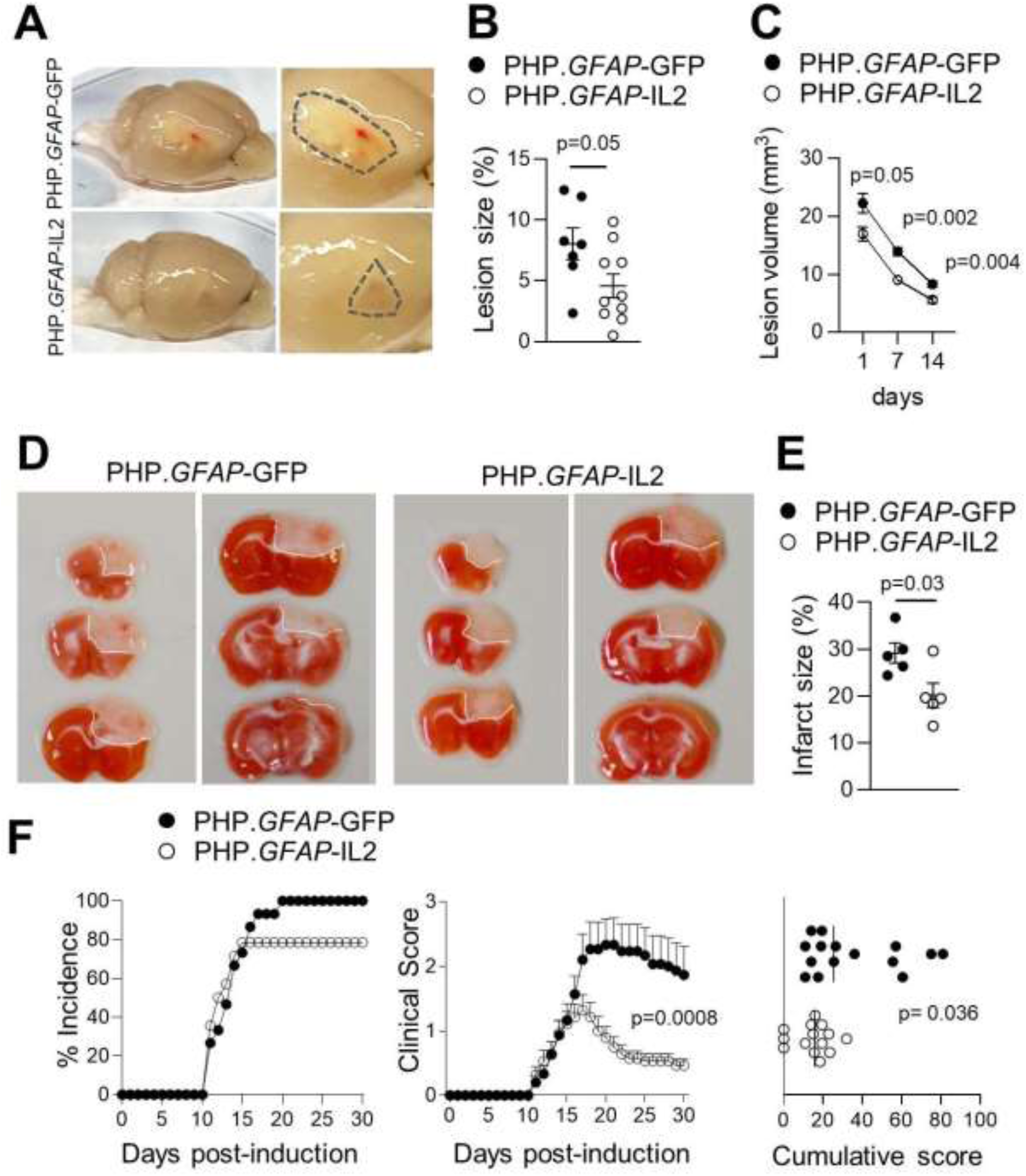
Neuroprotective utility for dual-lock IL2 gene delivery across multiple neuroinflammatory pathologies. **(A)** Wildtype mice, treated with control PHP.*GFAP*-GFP or PHP.*GFAP*-IL2 on day -14, were given a dMCAO stroke and examined at 15 days post-stroke for macroscopic damage (outlined) with **(B)** TTC-aided quantification of stroke damage (n = 7, 10) and **(C)** longitudinal MRI-based quantification of lesion size (n=11,17). **(D)** Wildtype mice, treated with control PHP.*GFAP*-GFP or PHP.*GFAP*-IL2 on day -14 (n = 5, 5), were given a photothrombotic stroke and examined one-day post-stroke for macroscopic damage (representative images, with lesion outlined) and **(E)** TTC-aided quantification of stroke damage. **(F)** EAE was induced in wildtype mice, following treatment with control vector (PHP.*GFAP*-GFP) or PHP.*GFAP*-IL2 on day -14 (n= 15, 14). Incidence, daily clinical score (mean ± SEM) and cumulative mean clinical score.

To investigate the translational potential of this approach, we tested PHP.*GFAP*-IL2 treatment in the curative context. First, we used the controlled cortical impact model, where pre-treatment with PHP.*GFAP*-IL2 was shown to reduce the size of the developing lesion (**Figure 5B,C**). Taking a curative approach, we first subjected mice to TBI and then treated the animals post-injury with PHP.*GFAP*-IL2. As with the preventative treatment, the curative approach reduced the size of the developing lesion compared to that observed in control treated mice (**Figure 8A**). In stroke, however, despite the positive benefits of a pre-treatment regime, curative PHP.*GFAP*-IL2 treatment after stroke induction did not reduce the resulting lesion size in either the dMCAO (**Figure 8B**) or photothrombotic (**Figure 8C**) models. This suggests either that the stroked brain is refractory to treatment, or that unfavorable kinetics of IL2 production preclude efficacy during the rapid damage that occurs following stroke. To discriminate between these possibilities we developed a model of secondary stroke. Photothrombotic stroke was induced in one hemisphere, mice were treated with PHP.*GFAP*-IL2 or control, and then 14 days later photothrombotic stroke was induced in the opposite hemisphere. In this context, treatment with IL2 following the primary stroke resulted in significant reduction of lesion size in the secondary stroke (**Figure 8D**), demonstrating both the limitations of the treatment window following stroke, and the potential clinical benefit this approach could have in treating secondary strokes. For MS, we again used the EAE model; however, we waited until the mice developed clinical manifestations and then treated with control PHP.B or the dual-lock PHP.*GFAP*-IL2. Strikingly, the protective effect of PHP.*GFAP*-IL2 was still observed, with separation of the clinical time-course by day 15 and a sharp reduction in the cumulative clinical score (**Figure 8E**).

**Figure 8.**
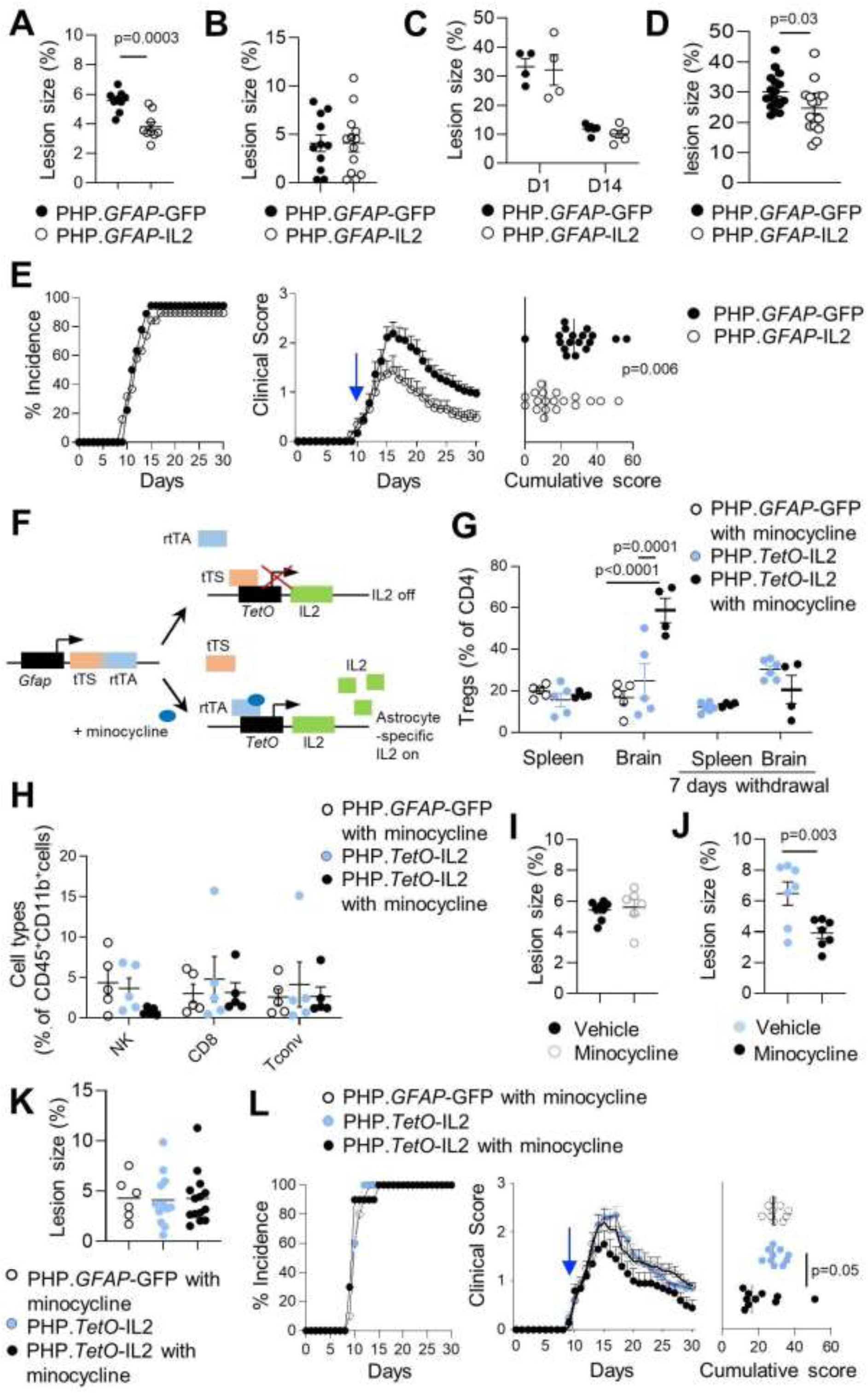
Neuroprotective utility for dual-lock IL2 gene delivery across multiple neuroinflammatory pathologies. **(A)** Mice were given controlled cortical impacts to induce moderate TBI, directly followed by treatment with PHP.*GFAP*-IL2 (or PHP.*GFAP*-GFP control vector), with examination 14 days post-TBI (n = 9,9). Quantification of cortical area lost due to lesion by histological analysis. **(B)** Mice were given a dMCAO stroke and immediately treated with control PHP.*GFAP*-GFP or PHP.*GFAP*-IL2. Mice were examined at 15 days post-stroke for TTC-aided quantification of stroke damage (n = 11,13). **(C)** Mice were given a photothrombotic stroke and immediately treated with control PHP.*GFAP*-GFP or PHP.*GFAP*-IL2. Mice were examined one day (n = 4,4) and 14 days (n = 6,5) post-stroke for TTC-aided quantification of stroke damage. **(D)** Mice were given a primary photothrombotic stroke, treated with control PHP.*GFAP*-GFP or PHP.*GFAP*-IL2, then given a secondary photothrombotic stroke in the opposing hemisphere. Mice were examined 14 days post-secondary stroke for TTC-aided quantification of stroke damage (n=17,17). **(E)** EAE was induced in wildtype mice and 10 days post-induction were treated with control vector (PHP.*GFAP*-GFP) or PHP.*GFAP*-IL2 (n = 18, 19) (blue arrow): incidence, daily clinical score (mean ± SEM) and cumulative mean clinical score were measured (n = 15, 14). Design of the ‘Tet On’ vector (PHP.TetO-IL2.*GFAP*-rtTA), with default suppression of IL2 converted into expression only in the presence of minocycline in transduced astrocytes. Wildtype mice were administered GFP control vector or PHP.TetO-IL2.*GFAP*-rtTA and gavaged daily with vehicle (PBS) or minocycline. Mice were assessed for the number of brain Tregs using flow cytometry 11 days after treatment. Additional groups were assessed 1 week after minocycline withdrawal (n = 4 - 5/group). **(H)** Frequency of CD8, NK and CD4 Tconv cells in the brain following GFP control vector or PHP.TetO-IL2.*GFAP*-rtTA and daily gavage with vehicle (PBS) or minocycline. **(I)** Wildtype mice were given controlled cortical impacts to induce moderate TBI, immediately followed by control vector, with or without minocycline. Quantification of cortical area lost due to lesion by histological analysis on day 14 (n = 8, 6). **(J)** Wildtype mice were given controlled cortical impacts to induce moderate TBI, immediately followed by treatment with PHP.TetO-IL2.*GFAP*-rtTA, with or without minocycline. Quantification of cortical area lost due to lesion by histological analysis on day 14 (n = 7,7). **(K)** Mice were given a dMCAO stroke and immediately treated with control vector plus minocycline, or PHP.TetO-IL2.*GFAP*-rtTA, without or with minocycline. Mice were examined at 15 days post-stroke for TTC-aided quantification of stroke damage (n = 6, 12, 14). **(L)** EAE was induced in wildtype mice, following treatment with control vector (PHP.TetO-GFP.*GFAP*-rtTA) or PHP.TetO-IL2.*GFAP*-rtTA, with or without minocycline on day 10 (n = 10/group) post-EAE-induction. Incidence, daily clinical score (mean ± SEM) and cumulative mean clinical score.

The development of a brain-specific IL2 delivery system provides the potential for use in clinical neuroinflammation contexts, where peripheral immune suppression and susceptibility to peripheral infections would prohibit a systemic delivery of IL2. Translation to the clinic requires, however, the ability for dose modification and withdrawal capacity. To add these clinically-desirable features, we altered the ‘dual-lock’ system to include a third layer of control through inclusion of a ‘Tet-On’ system. In this system, IL2 expression was shifted under the control of a TetO-dependent promoter, with the rtTA activator controlled by the *GFAP* promoter (**Figure 8F**). Furthermore, the rtTA was modified to allow response to the blood-brain barrier permeable drug minocycline ^26^, and was fused with a TetR to reduce baseline expression ^27^. This ‘triple-lock’ AAV provided the same brain-specific expansion of Tregs as the ‘dual-lock’ system, but only in the presence of minocycline (**Figure 8G**). Other major leukocyte populations present in the brain were unaffected in frequency (**Figure 8H**). Following the withdrawal of minocycline, brain Treg numbers returned to baseline levels within one week (**Figure 8G**). To validate this ‘triple lock’ system in the disease context, we first used TBI. Despite its reported anti-inflammatory properties, minocycline by itself did not alter the lesion size (**Figure 8I**). However, the combination of the ‘triple lock’ and minocycline treatment, given post-injury, substantially reduced the size of the resulting lesion (**Figure 8J**). In stroke, by contrast, the treatment did not alter lesion size, supporting an incompatibility between the kinetics of pathology and Treg expansion (**Figure 8K**). Finally, in EAE the combination of the ‘triple lock’ system and minocycline treatment, given after disease symptoms developed, resulted in earlier plateau of symptoms and lower cumulative pathology (**Figure 8L**). These results validate our triple-lock gene delivery system for brain-restricted IL2 expression as having both of the key clinical requirements of efficacy and dose-control, with potential use in neuroinflammatory pathologies with compatible disease progression kinetics, such as TBI and MS.

## Discussion

Here we demonstrated the utility of brain-specific IL2 as a neuroprotective agent and developed a delivery platform suitable for clinical application. While both direct and indirect mechanisms for IL2-mediated protection are plausible, the most parsimonious explanation remains the capacity of IL2 to expand the local Treg population. The brain provides a relatively IL2-deficient environment for Tregs, a state known to induce apoptosis and limit population size ^16^. With incomplete penetrance of brain Treg expansion following low-dose IL2 delivery, it is attractive to hypothesize that mice that failed to upregulate Treg numbers may correlate with clinical non-responders. As both assessments require whole brain, and therefore cannot be assayed simultaneously, this association remains speculative. Gene delivery resulted in increased brain IL2 concentration followed by a lagging increase in resident Treg frequency, with reproducible effects observed by day 5 in most mice. The delayed effect of IL2 supplementation on Treg numbers is consistent with the kinetics of clinical progression following treatment in the EAE context, where clinical scores of treated and control mice are initially concordant, with the beneficial effect observed from day 5 post-treatment. In the context of stroke, where current treatments sharply lose efficacy beyond 3 hours post-stroke ^28^, the delay in Treg expansion could account for the disparity between the protective pre-injury regime and the ineffective post-injury treatment. Likewise, the dependency of IL2 treatment on the presence of the adaptive immune system, as demonstrated through Rag-deficient mice given TBI and PHP.*GFAP*-IL2, is consistent with Tregs being the key mediator of the neuroprotective effect of brain-delivered IL2. Despite this supporting evidence, we do not discount the possibility that neurogenic IL2 works, at least in part, through effects on other cell types. While an alternative mechanism for IL2 neuroprotection is not identified, it is important to note that there is not a strict concordance between brain Treg frequency and treatment efficacy. In each neuroinflammatory model, the pathology-induced immune changes outweigh the treatment-induced immune changes at the assessed time-points. In part, these discrepancies may lie in technical limitations; both histological and brain Treg quantification approaches require the use of the whole brain, so we were unable to assess both measures simultaneously. However it is also important to consider that pathological modification and impact on clinical progression are not necessarily temporally coupled. Thus, a transient pulse of brain Treg expansion early on during disease or injury may drive long-lasting improvements in pathology at time-points where the treatment effect has been washed out. For example, in stroke, where efficacy is only observed in a pre-injury context allowing brain Treg numbers to be elevated at the point of injury, the critical treatment window is likely within hours. In this case, the treatment-induced Treg increase during the treatment window would be more relevant than the number of Tregs present at the histological analysis weeks later. By contrast a late inflammatory-response surge of Tregs, while comparable in size to the early treatment-response increase, is associated with poorer pathological outcomes. If local Treg expansion is the sole mechanism of action for neuroprotective IL2, the potential for efficacy must therefore be contingent on the size of the pathology-specific treatment window, the kinetics of treatment-induced Treg expansion, and potential interplay between pathology and treatment kinetics. Alternatively, neuroprotective IL2 may function via alternative mechanisms, perhaps in synergistic effect with the Treg expansion.

Without discounting the possibility of non-canonical effects of IL2, the expansion of brain-resident Tregs provides a functionally-dynamic mediator for local immune modulation. Tregs are capable of producing multiple anti-inflammatory agents, from cell-surface proteins, such as CTLA4, to secreted cytokines, including IL10 and Amphiregulin, as well as small molecules, for instance cAMP ^29^. Many of these immunosuppressive mediators are used in clinical practice as anti-inflammatory agents and have been proposed to treat neuroinflammation ^30,31^. Tregs also possess key reparative functions, beyond their direct immunosuppressive role ^9,32^. The use of IL2 to expand the Treg population bypasses the problem of identifying the ideal immunosuppressive mediator, and instead piggy-backs on the adaptive properties of Tregs, which are capable of sensing and responding to local microenvironmental cues and initiating effective immunosuppressive programs ^29^. The small number of brain-resident Tregs, however, even following local expansion, suggests a more common cell type is likely to be required as an effect amplifier. Microglia are an attractive candidate for this putative intermediatory, being both the primary immunological cell of the brain and being heavily implicated in neuroinflammation ^33^. In the context of TBI, while both treated and untreated mice demonstrated extensive microglial activation, the transcriptional profiles of the activation states were heavily modified by treatment. In particular, while activated microglia in control-treated TBI gained the classical inflammatory transcriptional profile, a substantial subset of activated microglia in IL2-treated mice sharply upregulated MHCII expression, without additional inflammatory markers. This upregulation correlated with localization along the injury border, indicating a potential function as a buffer to the expanding zone of neurotoxicity. The association between MHCII upregulation and restraint in inflammatory marker expression is intriguing. Early upregulation of MHCII on microglia has previously been associated with enhanced protection and repair of the injured CNS, in the context of optic nerve crush injury ^34^. MHCII expression indicates enhanced capacity for direct cognate interaction between microglia and Tregs, and may allow increased local production of multiple anti-inflammatory mediators, such as IL10 (upregulated during EAE) and amphiregulin (upregulated during TBI). Alternatively, CD74, a chaperone for MHCII, is highly upregulated in our system, in parallel with MHCII, and has been demonstrated to directly impede the polarization of microglia to the inflammatory state ^35^. Within the DAM compartment, IL2-treatment also resulted in greater upregulation of SPP1/osteopontin, which has been similarly demonstrated to impede inflammatory polarization of microglia ^36^, and has been proposed as a therapeutic in TBI and other neuroinflammatory diseases ^37^. We caution, however, against overly simplistic models of a single molecular mediator responsible for the neuroprotective effect of brain-resident Tregs, with the multipotent functions of these cells more compatible with complex synergistic effects, changing between models and inflammatory stages.

Despite the biological potency of IL2, incorporation into therapeutics has been slow. The short half-life of only 15 minutes necessitates either high doses, which in turn alter the biological targets ^38^ (including direct effects on the blood-brain barrier ^39^), or constant delivery ^40^ (to prevent the IL2 withdrawal-mediated contraction phase of Treg homeostasis ^16^). Proof-of-principle studies have demonstrated the capacity of IL2 to delay neuroinflammatory or neurodegenerative disease ^20-22^, including via AAV-mediated systemic delivery of IL2 ^19^. However the biological potency of Tregs precludes such approaches from clinical translation: systemic expansion of Tregs impedes potentially advantageous systemic immune responses. Global immunosuppression is not a viable strategy to combat neuroinflammatory disease, especially considering that such patients show a pre-existing enhanced susceptibility to infections ^41,42^. Indeed, expansion of the circulatory Treg population by systemic IL2 delivery likely means that off-target peripheral immune suppression would be greater than the desired on-target CNS immune suppression. The recent identification of brain-resident Tregs ^7^ provides an alternative approach: if sustained local IL2 production can be achieved it could expand the brain-resident Treg population, bypassing the unpalatable consequences of systemic IL2 delivery.

Here we used αCamKII^+^ neurons as the source of local IL2 production in the proof-of-concept phase, based on the observation that neurons are the primary source of IL2 in the healthy brain ^43^. In a therapeutic setting, however, astrocytes potentially have superior properties as a delivery source, not least of which is their higher robustness to injury and expanded productive capacity following insult. Astrocytes are a major non-neuronal cell type found throughout the CNS, including the spinal cord, and their non-overlapping ‘tiled’ arrangement allows for dispersed spatial distribution of production. As key players in CNS homeostasis, astrocytes provide trophic support to neurons ^3^, facilitated by their highly efficient secretory system ^44^. Here we demonstrate that this system can be effectively ‘hijacked’ and exploited for local IL2 production and secretion, with the additional advantage that astrocytic endfeet are in close proximity to the vasculature zones where brain-resident T cells are concentrated ^7^. The involvement of astrocytes in the pathophysiology of neurological injury and disease potentially acts as a biological amplification process. Reactive gliosis is poorly understood at the molecular level but, when unresolved, appears largely deleterious, being concomitant to neuronal loss ^45^. However, for our purposes, the upregulation of the *GFAP* promoter and astrogliosis typically seen following trauma, served to concentrate IL2 production near to regions of reactive astrogliosis and so-called glial scar. The system, therefore, contains the features of a natural ‘rheostat’, using the molecular signature of neurological damage to both amplify and anatomically direct the therapeutic response.

Gene delivery systems, such as the triple-lock IL2 system developed and validated here, have high potential for translation to the patient context. While early setbacks delayed the clinical uptake of AAV-based systems, improved vectors with superior safety profiles are gaining regulatory approval ^46^, including Luxturna for inherited retinal disease (suborbital injection) and Zolgensma (intravenous) for spinal muscular atrophy, with other CNS diseases under intensive investigation ^47^. The ability of AAV-based vectors to transduce both dividing and (critical for the CNS) non-dividing cells, with non-replicative and largely episomal gene delivery, combined with low inherent immunogenicity avoids many of the limitations of alternative vector types, such as adeno- and lentivirus-based systems. Furthermore, as AAV-based vectors provide strong and sustained transgene expression (over 4 years in human patients ^48^ and 15 years in non-human primate ^49^), use of such a system should ensure long-term therapeutic benefits - an attractive proposition in progressing or relapsing diseases such as MS, or injuries with a chronic component, such as TBI. Even in stroke, where the narrow treatment window seemingly precludes efficacious post-injury treatment, a lasting gene delivery of IL2 may have clinical benefit; 10% of stroke patients experience a secondary stroke within 90 days ^50^, with poor treatment options available ^51^. While PHP.B was used here for its superior transduction of the murine CNS following intravenous injection ^23^, poorer results have been observed in non-human primates ^52^ and are expected, by extension, in humans, due to lack of the LY6A receptor on brain microvascular endothelial cells ^53^. Both direct injections or intrathecal delivery can potentially overcome this problem ^54^ but remain highly invasive procedures associated with significant clinical complications. The adoption of alternative AAV capsids, showing more efficient CNS penetration in humans following systemic delivery, would retain the clinically non-invasive nature of the system described here and, in the case of engineered capsids, likely reduce issues related to pre-existing immunity reducing transduction levels ^55^, as well as reducing potential off-target toxicity. As the delivery system developed here relies on a modified version of the endogenous promoter, rather than capsid, for lineage specification, it can be readily adapted to alternative AAV capsids for human use. More efficient CNS penetration by novel engineered capsids would also allow substantially lower doses to be used, again reducing potential toxicity, as well as reducing manufacturing costs, which remain high at this time. Additional refinements may arise from the recent findings of molecular heterogeneity among astrocytes ^56,57^, providing promoter elements allowing specific microanatomical targeting of therapeutic delivery. The coupling to small-molecule inducers, such as validated here, provides both a dose-escalation function and a safety-withdrawal capacity. Minocycline-based systems combine the appropriate biodistribution with the intriguing potential for synergy, as minocycline itself is a mild neuroprotective agent, with potential efficacy in MS, TBI and stroke ^58-60^. The lack of viable alternatives to treat the neuroinflammatory component of CNS pathologies warrants the further investigation of the triple-lock IL2 delivery system for clinical development.

## Acknowledgements

This manuscript is dedicated to the memory of Russell Liston, who died too young from traumatic brain injury. The work was supported by the VIB, an ERC Consolidator Grant TissueTreg (to A.L.), an ERC Proof of Concept Grant TreatBrainDamage (to A.L.), an ERC Starting Grant AstroFunc (to M.G.H.), the ERC Proof of Concept Grant AD-VIP (to M.G.H.), FWO Research Grant 1513616N (to M.G.H.), Thierry Latran Foundation Grant SOD-VIP (to M.G.H.), an ERNAET Chair (H2020-WIDESPREAD-2018-2020-6; NCBio: 951923) (to M.G.H.), FWO Research Grant 1503420N (to EP), an SAO-FRA pilot grant (20190032, to E.P.) and the Biotechnology and Biological Sciences Research Council through Institute Strategic Program Grant funding BBS/E/B/000C0427 and BBS/E/B/000C0428, and the Biotechnology and Biological Sciences Research Council Core Capability Grant to the Babraham Institute. E.P., V.L., M.M., P.G. and A.d.B. were supported by fellowships from the FWO. RL is a senior clinical investigator of FWO Flanders. PB and CPF were supported by an ERA-NET-NEURON grant EJTC 2016 to CPF and by The Netherlands Organization for Scientific research (NWO). The VIB and Babraham Institute are owners of a patent based on work included in the manuscript, with L.Y., E.P., J.D., M.G.H. and A.L. potential financial beneficiaries of commercialization. The authors acknowledge the important contributions of Jeason Haughton (VIB) for mouse husbandry, Melvin Rincon (VIB) for advice on AAV design and production, Krist’l Vennekens for technical support, Pier-Andrée Penttila and the KUL FACS Core, Jens Wouters and the KUL Molecular Small Animal Imaging Center (MoSAIC), Simon Walker and the Babraham Institute Imaging Core, the VIB Bio-Imaging Core, the VIB Single Cell Sequencing Core, and Ronald Breedijk, Mark Hink and the Leeuwenhoek Center for Advanced Microscopy at the University of Amsterdam.

