## Supplementary methods and figures for "Astrocyte-targeted gene delivery of interleukin 2 specifically increases brain-resident regulatory T cell numbers and protects against pathological neuroinflammation"

### Materials and methods

#### *Mice*

Foxp3-Cre transgenic mice [1], αCamKII-CreERT2 transgenic mice [2], PLP1-CreERT transgenic mice [3], IL2-GFP mice [4], and Rag1 knockout mice [5] were used on the C57BL/6 background. RosaIL2 mice were generated through the insertion of a cassette containing a floxed-STOP sequence followed by an IL2-IRES-GFP sequence into the Rosa26 locus, using the endogenous Rosa26 promoter [6], and were used on the C57BL/6 background. Mice were housed under SPF conditions, under a 12 hour light/dark cycle in a temperature and humidity-controlled room with *ad libitum* access to food and water. All animal procedures were approved by the KU Leuven Animal Ethics Committee (P035/2015, P015/2014, P209/2015, P043/2016, P082/2018, P124/2019,), the University of Amsterdam (CCD 4925, AVD1110020184925), or the Babraham Institute Animal Welfare and Ethics Review Body (PP3981824) taking into account relevant national and European guidelines. Both male and female mice (8-12 weeks old) were used in this study, unless otherwise specified. Age and sex of study mice and treatment regime was selected in consultation with the Animal Ethics Committees. Tamoxifen (Sigma T5648) was solubilized in corn oil (Sigma) at 10 mg/ml. Five to seven weeks old mice were injected 3 times, via intraperitoneal injection, at 48 h intervals using a dose of 100 mg/kg body weight. Minocycline (PBS vehicle) was administrated at 50 mg/kg body weight, through daily oral gavage. Functional quantification of the blood-CSF barrier and blood-brain barrier permeability was performed through *in vivo* dye application and *ex vivo* dye penetration assessment, as previously described [7]. Sample sizes for mouse experiments were chosen based on power calculations and pilot data, in conjunction with the Animal Ethics Committee, to allow for robust sensitivity without excessive animal use. Mice were selected randomly for inclusion into the various experimental groups, with the animal technicians performing experimental procedures and clinical measurements blinded as to the identity of experimental groups.

#### *Parabiosis*

For parabiosis, pairs of 7-10 week-old female mice were co-housed for 14-21 days prior to surgery. C57BL/6.SJL-*Ptprc*$^{a}$/BoyJ mice (CD45.1) were parabiosed to αCamKII$^{IL2}$ mice (CD45.2), pre-treated with tamoxifen, for 10 weeks. Pairs of mice were anesthetized with inhaled isoflurane, 3.5% v/v induction, 2.5-3.0% v/v maintenance. Carprofen and buprenorphine were delivered intraperitoneally at a dose of 10 mg/kg and 0.1 mg/kg prior to surgery. Fur was removed from the surgical site. Mice were laid supine and the surgical site was disinfected with betadine solution, followed by 70% EtOH. Longitudinal skin incisions were made to the shaved sides of each animal, starting at 0.5cm above the elbow and extending all the way to 0.5cm below the knee joint. The skin was gently detached from the subcutaneous fascia to create 0.5 cm of free skin, and sutured together to generate a parabiotic pair.

*Traumatic Brain Injury*

A moderate cortical traumatic brain injury (TBI) was induced using the controlled cortical impact model as described previously [8], but with minor adjustments. In brief, male mice were treated with tamoxifen (6 weeks old) or vector (10 weeks old). At 12 weeks, mice were anesthetized using 5% isoflurane and placed in a stereotaxic frame. Mice were kept anesthetized using 2% isoflurane throughout the procedure. A craniotomy was performed creating a window over the left hemisphere, ranging from lambda to bregma. The impact piston (Leica Impact One) had a 3mm metal tip and was placed on top of the left cortex at a 20-degree angle. An impact was performed using the following settings: 5.5 m/s velocity, 1 mm impact depth, 300 ms dwell time. Immediately after the impact the skull bone was replaced and attached using superglue. The skin was stitched to close the wound and mice were allowed to recover on a heat pad until fully awake. The sham group underwent craniotomy but did not receive the impact. The TBI-induced perfusion deficit in the brain was measured by magnetic resonance imaging (MRI) at 24 hours and on days 7-14 after impact. Flow cytometry profiling and immunohistochemistry analysis were performed 14 days after impact.

*Experimental Autoimmune Encephalomyelitis*

Experimental autoimmune encephalomyelitis (EAE) was induced in female mice, aged 8-12 weeks. To induce active EAE, mice were immunized with 50 μg of $MOG_{35–55}$ peptide (Covalab) emulsified with Complete Freund Adjuvant (CFA) containing 2 mg/ml of *Mycobacterium tuberculosis* (Sigma). 200 ng/ml pertussis toxin (List Biochemicals) was given on day 0 and day 2 after immunization. Clinical score was evaluated blind by a technician on a scale from 0 to 5 [9].

*Photothrombotic stroke*

Focal cortical ischemia was induced using the photothrombotic lesion model, as previously described [10]. Briefly, mice (male, 12 weeks old) were anesthetized with 2.5% isoflurane (Halocarbon) in an oxygen/air mixture, respiration was monitored and rectal temperature was maintained at 37 ± 0.5°C with a heating plate (TCAT-2LV Controller, Physitemp Instruments). After fixation in a stereotactic frame attached to a digital display (David Kopf Instruments), the skull was exposed by a 1 cm midline incision of the skin. 100 µl rose bengal (Sigma) at a concentration of 3 mg/ml in saline, was injected via the tail vein. For illumination, a 2.4 mm laser beam with a wavelength of 565 nm (L4887-13, Hamamatsu Photonics) was focused on the motor cortex responsible for forepaw function (0.5 mm rostral, 1.8 mm lateral of bregma). 5 seconds after rose bengal injection, the brain was illuminated though the intact skull for a total duration of 5 minutes. After illumination, the incision was sutured and animals were given 500 µl saline and 0.05 mg/kg vetergesic (ecuphar) subcutaneously. During recovery, mice were placed in a separate cage with half of the cage placed under an ultraviolet lamp before returning to their home cage and housing facility. After stroke, mice were monitored on a daily basis during the first week; after that animal health was checked weekly. In mice with a secondary stroke, treatment with AAV occurred i.v. after the primary stroke. Mice were then allowed to recover for two weeks before receiving a secondary stroke on the opposite hemisphere, using the same procedure. To prevent unnecessary suffering of animals, mice were euthanized if they showed severe weakness or lost 20% of their bodyweight within 5 days.

24 hours after stroke, mice were anesthetized with an overdose of dolethal (20 mg/ml, vetoquinol) and transcardially perfused with PBS. Brains were collected and cut in 1 mm sections using a mouse brain matrix (Zivic Instruments). For each animal, a total of 6 sections surrounding the infarct were collected and incubated in a 1% (w/v) 2,3,5-triphenyl tetrozolium chloride (TTC, Sigma-Aldrich) in PBS solution for 25 minutes at room temperature, while protected from light. When stained, sections were put on glass plates and pictures were taken with a regular photo camera (Nikon). To correct for edema developed during the early phase of ischemia, stroke area was calculated according to the method

provided by Swanson and colleagues [11]: [lesion area] = [area of the contralateral side] - [total undamaged area] and presented as a percentage of the contralateral hemisphere. The identification of the undamaged ipsilateral area involved mapping the zone surrounding the combined missing scar tissue and TCC-stained ischemic zones.

*Permanent Distal Middle Artery Occlusion*

10 weeks old male mice were subjected to permanent distal middle cerebral artery occlusion (dMCAO), essentially as described [12], but with minor modifications. Briefly, the mice were anesthetized with 2% (v/v) isoflurane and placed in a lateral position on a heat pad to maintain body temperature. Eye ointment (duratears®, Alcon) was applied to prevent dehydration of the eyes. The surgical site was shaved and disinfected with ethanol and a vertical skin incision was made between the left ear and eye. Next, a surgical window was opened in the skin and the temporal muscle was separated, dorsal and apical, with surgical scissors to expose the temporal bone without removing the muscle. The MCA was identified and a hole was burred with a microdrill (Stoelting) at the site of MCA bifurcation. The remaining bone and the overlying dura mater were subsequently removed with forceps. Permanent occlusion of the MCA was then performed with bipolar coagulation forceps (0.4 mm tip; ERBE) with the electrosurgical unit (ERBE ICC 50) set at 8 W. During surgery, the surgical site was kept hydrated using saline. After visual confirmation MCA occlusion (reduced blood flow), the muscle was put back into place and the wound was sutured, disinfected, and the animals were allowed to recover in a pre-heated environment. Mice that developed subarachnoid haemorrhage during surgery were excluded from the study. Sham animals followed the same surgical procedure except for the final coagulation step.

*Magnetic resonance imaging*

Mice with TBI were scanned 1 day, 1 week, 2 weeks, 4 weeks and 3 months after injury. Mice with dMCAO were scanned 1 day, 1 week and 2 weeks after injury. MRI measurements were performed on a 9.4 T Bruker Biospec small animal MR system (20 cm horizontal bore, Bruker Biospin), using a quadrature resonator with an inner diameter of 7.2 cm for transmission and an actively-decoupled mouse brain surface coil for receiving (Rapid Biomedical). The scanner was equipped with an actively shielded gradient set of 600 mT/m. Mice were scanned under isoflurane anesthesia (1-2% (v/v) isoflurane in 100% (v/v) $O_2$

administered through a snout mask). Rectal temperature and respiratory rate were continuously monitored (SAII), and isoflurane levels were adjusted to maintain a respiratory rate of 80-100 breaths per minute. Rectal temperature was maintained at 37°C (36-37.5°C).

Following an initial localizer scan, the MRI protocol used an axial $T_2$-weighted spin echo sequence with a repetition time (TR) of 4.5 s, effective echo time (TE) of 40 ms, rare factor 8, 1 average, matrix 256x256, field of view (FOV) 20x20 mm, and 24 slices of 500 µm thickness. A multi-slice-multi-echo sequence was acquired for the calculation of parametric T2-maps using the same slice orientation as for the $T_2$-weighted MRI and the following parameters: TR 4.0 s, 12 TE increments of 12 ms, 1 average, matrix 128x128, FOV 20x20 mm, 24 slices of 500 µm thickness. A diffusion weighted MRI was acquired for the calculation of parametric apparent diffusion coefficients (ADC map) using the following parameters: TR 2.0 s, TE 20 ms, 1 average, matrix 128x128, FOV 20x20 mm, 20 slices of 500 µm thickness with a 200 µm gap in between slices, b-values of 0, 100, 300, 500, 800, 1000 and 1500. Finally, a three-dimensional gradient echo sequence (FLASH) with the following parameters was acquired: TR 30 ms, TE 7 ms, 20˚ pulse, matrix 160x160x96, FOV 20x20x12 mm, resulting in an isotropic resolution of 125 µm. Operators were masked to experimental group. All MR images were processed using the Bruker Biospin software Paravision 6.1. Parametric $T_2$ and ADC-maps were calculated in Paravision 6.1, through a pixel wise mono-exponential fit. For quantification of lesion volumes, Paravision 6.1 software was used.

*Behavioral experiments*

Behavioral experiments in healthy mice were performed in 13-18 weeks old female mice. Mice were habituated to their new environment for at least 5 days and tests were conducted during the light phase of their activity cycle. Behavioral experiments in TBI mice were initiated on day 16 post-TBI and were conducted during the dark phase of their activity cycle. The general health and weight of the mice were routinely recorded during the testing. Tests were performed and analyzed by an observer blind to the experimental group. If tracking error(s) occurred, mice were excluded. Behavioral experiments were recorded using infrared cameras. Sham operated littermates were used as controls.

*Open-field:* Open-field exploration was tested in a 40 cm × 40 cm × 30 cm (w×l×h) square arena illuminated by indirect light. Animals were dark adapted for 30 min and tested in the arena for 10 min. Movements of the mice in the arena were video-tracked for 10 minutes with Ethovision software (Noldus).

*Nest building:* For the nest building test, mice were single-housed with 1.5 g of cocoon nesting pellets and left undisturbed for 24 hours. Nest building was scored according to published protocols [13].

*Sociability:* Sociability was evaluated using the three-chamber test [14, 15]. The set-up consisted of a rectangular transparent plexiglass box divided into three compartments, which were separated by two partitions. The central chamber (42 x 26 cm) was connected to a left and right chamber (26 x 26 cm) connected by guillotine doors (6 x 8 cm). The test consisted of two consecutive stages: acclimation stage and sociability stage. After habituation to the central compartment (5 min), wire cages (11 x 10 cm) were placed in the external chambers, which either contained a 'stranger' mouse (same sex, 'stranger 1') or were left empty: approach behavior to either of the wire cages was recorded for 5 min (sociability). Animal behavior was recorded using a webcam and ANY-maze™ Video Tracking System software (Stoelting Europe).

*Forced swim test:* Mice were individually placed into a glass cylinder (20 × 14 cm) filled with water (16 cm depth, 25 ± 1°C). Individual mice were placed gently in the water and left for 6 min: the degree of immobility (passive floating, no active swimming or escape behavior) was scored for the final 4 min using ANY-maze™ Video Tracking System software (Stoelting Europe).

*Light/dark test:* The apparatus used for the light/dark test consisted of a cage (40 × 40 × 30 cm) divided into two compartments of equal size by a partition with a door. One compartment was brightly illuminated, while the other was dark. Mice were placed in the dark compartment and allowed to move freely between the two chambers for 10 min. Mouse activity in the light compartment was tracked with Ethovision equipment and software (Noldus).

*Rotarod test:* Motor coordination and equilibrium were tested with a rotarod setup (MED Associates Inc.). Mice were first trained at constant speed (4 rpm for 2 min) and then tested on four trials (inter-trial interval, 10 min). During the test trials, the animals had to balance on a rotating rod that accelerated from 4 rpm to 40 rpm over a period of 300 seconds. The latency to fall off the rod was recorded (up to a 5 min cut-off point).

*Contextual discrimination fear conditioning:* Two identical conditioning boxes (25 × 25 × 25 cm) with stainless steel grid floors to deliver shocks were located in sound-attenuating dark cubicles. Animal movement was monitored by motion-sensitive platforms connected to an interfaced computer using Panlab Freezing v1.2.0 software. The degree of motion measured ranged from 0 to 100 (arbitrary units). Freezing, defined as the absence of movement except for breathing, was recorded when activity remained below a validated threshold of 2.5 arbitrary units for at least 1 second. Contextual fear conditioning was measured in three different contexts. Two contexts (A and C) were very similar in tactile (grid floor), olfactory (odor) and visual (dark) dimensions, but C had in addition an inserted 'A-frame' roof (adapted from [16]). In contrast, context B was different in all dimensions. On days 1 - 3, mice were placed in context A, and after 3 minutes a single mild foot shock (2 s, 0.5 mA) was applied to induce contextual fear conditioning. After another minute, the animals were placed back in their home cage. On days 4 and 5, fear memory to A, B, or C was measured by placing the animal for 4 minutes in each of the contexts and measuring the amount of freezing. The amount of freezing in C compared to A indicated the amount of contextual generalization.

*Morris water maze (MWM):* Spatial learning and cognitive flexibility were tested in the hidden platform MWM. For the healthy mouse experiments, a circular pool (150 cm diameter) was filled with opaquified (0.01% Acusol OP301, Dow Chemicals) water (26 ± 1°C). The platform (15 cm diameter) was hidden 1 cm underneath the surface of the water. For spatial learning, the mice were trained for 10 days to a fixed platform position [17, 18]. To evaluate reference memory, probe trials (100 seconds) were conducted on days 6 and 11 during acquisition learning. During probe trials, floater mice were excluded. For cognitive assessment in TBI mice, a modified MWM was performed on days 19-26 post-TBI or sham surgery, as previously described [19]. In brief, on day 19 mice were habituated to the water maze over 2 sessions lasting 1 minute each. Starting on day 20, the escape platform was positioned in the target quadrant and spatial cues were placed on the wall surrounding the maze. All animals underwent 2 sessions per day, with a 30 minute interval, for 5 consecutive days to assess spatial acquisition. During each session, mice were placed in randomized starting quadrants facing the maze wall. Trials lasted a maximum of 60 seconds, after which mice were placed on the escape platform if they did not succeed themselves. On day 26, a probe trial was carried out to assess reference memory. The escape platform was removed from the pool and mice were allowed to explore the maze for 60 seconds. For assessments in both healthy and TBI mice, swim paths were tracked with Ethovision software (Noldus).

*Novel object recognition:* The novel object recognition test was performed on mice on day 17 post-TBI or sham surgery, as previously described [20]. In brief, mice were habituated to the test environment on day 15, followed by training on day 16 (free exploration of the environment containing two identical objects for five minutes) and testing on day 17 (replacement of one familiar object by a novel object with five minutes of exploration). The ratio of time spent exploring the novel object over the familiar object was calculated and used as a proxy for learning and memory. Activity was tracked and semi-automatically quantified using Ethovision software (Noldus).

*AAV vector production and purification*

AAV-PHP.B production was performed by Vigene Sciences (Rockville, MD, USA) or VectorBuilder (Neu-Isenburg, Germany), using the classical tri-transfection method, with subsequent vector titration performed using a qPCR-based methodology [21, 22]. For AAV-PHP.B.*GFAP*-IL2 and PHP.B.*αCamKII*-IL2, the mouse IL2 coding sequence, together with 5' and 3' UTR (accession number BC116845) was cloned into a single stranded AAV2-derived expression cassette, containing a 2.2kb human *GFAP* promoter [23] or full length murine *αCamKII* promoter (Gene ID: 12322), woodchuck hepatitis post-transcriptional regulatory element (WPRE) and bovine growth hormone polyadenylation (bGH polyA) sequence. Control vectors were prepared by swapping the IL2 coding sequence for that encoding enhanced green fluorescent protein (EGFP, Vector Biolabs).

PHP.B.TetO:IL2.sGFAP:TetR-T2A-rtTA(V7/V14) was constructed with a 7xTetO sequence and minimal CMV promoter driving IL2, and a short human *GFAP* promoter driving a TetR-T2A-rtTA fusion protein modified to include the V7/V14 mutations for enhanced minocycline response.

In both cases, vector (100 µl total volume) was administered to mice via the intravenous route at $1x10^9$ vector genomes/dose, unless otherwise specified. Batch concentration was normalized using brain Treg expansion as a biological read-out. Mice were used for experimental procedures at least 14 days after AAV injection, unless otherwise indicated.

*Flow cytometry*

Mice were deeply anaesthetized with intraperitoneal injection of a ketamine (87 mg/kg), xylazine (13 mg/kg) mixture. Blood was collected from the right ventricle prior to transcardial perfusion with ice cold PBS. Blood was prepared by red blood cell lysis; single-cell suspensions from lymphoid organs were prepared by mechanical dissociation; single-cell suspensions from brain tissue were prepared by digestion for 30 minutes at 37ºC with 1 mg/ml collagenase IV (Thermo Fisher), 300 µg/ml hyaluronidase (Sigma-Aldrich) and 40 µg/ml DNase I (Sigma-Aldrich) in RPMI 1640 supplemented with 2 mM $MgCl_2$, 2mM $CaCl_2$ 20% FBS and 2 mM HEPES (Gibco), followed by mechanical disruption, filtration (through 100 µm mesh) and enrichment for leukocytes by gradient centrifugation (40% Percoll GE Healthcare, 600 x g, 10 min, no brake). For estimation of absolute cell numbers, counting beads were 'spiked in' at the initial step, allowing for calculation of cell loss during preparation. Non-specific binding was blocked using 2.4G2 supernatant. To assess intracellular cytokine production, cells were cultured for 4h in the presence of phorbol mysristate acetate (1 µg/ml, Sigma-Aldrich), ionomycin (1 µg/ml, Sigma-Aldrich), and brefeldinA (BD). Cells were fixed and permeabilized with the eBioscience Foxp3 staining kit (eBioscience). Cellular phenotypes were assessed using high parameter flow cytometry panels, containing markers to identify cell types and markers to assess activation states. Data were acquired on a BD FACSymphony, with panels covering (i) CD45, CD4, CD8, CD3, CD19, NK1.1, Foxp3, eBioscience™ Fixable Viability Dye eFluor™ 780, CD103, CD62L, GITR, CD25, Neuropilin, ST2, PD-1, KLRG1, Helios, CD69, ICOS, CD44, T-bet, TCRγδ and Ki67 or (ii) CCR6, CD80, TCRγδ, CD45, Foxp3, MHCII, eBioscience™ Fixable Viability Dye eFluor™ 780, IL1β, CD25, Ly6G, ST2, CX3CR1, PD-L1, TNF, CD44, Ki67, CD4, Ly6C, TrkB, CD19, CD69, CD8α, LAMP1, CD64, CD11b, CD3, TGFβ, or (iii) Foxp3, eBioscience™ Fixable Viability Dye eFluor™ 780, IL5, IL6, IL17, CD4, IFNγ, CD8α, TNFα, CD3, Amphiregulin, IL10, IL4, CD11b, CD19, GM-CSF, TCRγδ, pro-IL1β, TCRβ, IL2, NK11. For brain panels, the entire brain sample was acquired. Data was compensated using AutoSpill [24]. Examples for murine cells are always presented as concatenated biological replicates. Cell sorting was performed using a BD FACSAria III with a panel including CD4, CD11b, CD45, TCRβ, eBioscience™ Fixable Viability Dye eFluor™ 780 and CD25. Representative gating for brain Treg quantification is shown in **Supplementary Figure 25**.

*Fluorescence immunostaining*

Mice were deeply anaesthetized with intraperitoneal injection of a ketamine (87 mg/kg) / xylazine (13 mg/kg) mixture and transcardially perfused with PBS followed by 4% buffered formalin solution. The brain was removed and fixed in 10% buffered formalin solution overnight and stored in 30% sucrose until preservation in tissue freezing medium (Shandon™ Cryomatrix™ embedding resin, Thermo Scientific), and stored at -80°C. Sections (20-50 µm) were washed 15 minutes in 50 mM $NH_4Cl$-PBS and pre-blocked with 10% normal donkey serum in 0.5% Triton-X-100-PBS for 1h at room temperature. Aldh1l1 immunofluorescence required heat-induced epitope retrieval at 80°C for 30 minutes in a 10 mM sodium citrate buffer (pH 6.0). Sections were incubated overnight at 4ºC with primary antibodies directed against Foxp3 (1:500, MAB8214, R&D systems), CD4 (1:250, 100506, Biolegend), Iba1 (1:1000, 014-19741, Wako), GFAP (1:500, ab4674, Abcam), CD31 (1:100, MA3105, Invitrogen), S100ß (1:1000, S2532, Sigma-Aldrich), APC (1:250, ab16794, Abcam), NeuN (1:500, ABN90P, Millipore), GFP (1:300, 132002, Synaptic Systems; 1:1000, 600-401-215, Rockland; 1:500, 600-101-215, Rockland), GFAP (1:1000, 173004, Synaptic Systems), PDGRα (1:200, APA5, BD Pharmingen), Aldh1l1 (1:200, ab87117, Abcam), MHCII (1:400, eBiosciences, 14-5321-82). Subsequently, the sections were incubated for 90 minutes at room temperature with appropriate fluorophore-conjugated secondary antibodies (Thermo Scientific, Biolegend). After each antibody incubation, slices were washed 3 times for 10 minutes with 0.1% Triton-X-100-PBS. All sections were incubated with DAPI (1:1000) for 15 min and mounted with ProlongGold (Invitrogen) or Fluoromount-G (Southern Biotech). Images were obtained using a Zeiss LSM780 confocal microscope (60X Apochromat/NA 1.4), automated upright Leica DM5500 B microscope (20X HC Plan-Apochromat/NA 0.70), Nikon A1R Eclipse Ti confocal (60X Apochromat/NA 1.4), or a Zeiss Axioscan Z.1 slide-scanner (20X Plan-Apochromat/NA 0.8) equipped with a Hamamatsu Orca Flash 4.0 V3 camera. Axioscan images were exported and compressed to 10% file size. Fluorescence measurements were corrected for background. Subsequent image processing was performed using ImageJ (https://imagej.nih.gov/ij/download.html).

Structural integrity of the blood-brain barrier was assessed following transcardial perfusion of mice with ice-cold 4% PFA-PBS. Subsequently, brains were extracted from the skull and split into two halves (mid-sagittal). The right hemispheres were embedded in Frozen Section Medium (Thermo Fisher) immediately in cryomolds (Sakura) which were frozen on dry ice and stored at -80°C until further use. The left hemispheres were post-fixed overnight 4% PFA-PBS at 4°C. After dehydration, samples were embedded in paraffin in cryomolds and stored at room temperature until further use. The brains were cut into 5 µm slices for paraffin sections (HM 340 E, Thermo Fisher) or 20 µm slices for cryosections (CryoStar NX70, Thermo Fisher). Cryopreserved sections were used to stain for ZO-1 (1:500, 617300, Invitrogen), Claudin-1 (1:200, 51-9000, Thermo Fisher), E-cadherin (1:500,610181, BD) and CD31 (1:100, DIA-310, Dianova). Paraffin sections were used to stain for Occludin (1:100, 33-1500, Invitrogen) and CD31 (1:100, DIA-310, Dianova). Sections were permeabilized in 0.3% Triton X-100-PBS. Following blocking with 5% normal goat serum in 0.3% Triton X-100-PBS at room temperature for 1 h, sections were incubated with primary antibodies in blocking solution and left at 4°C overnight. After washing with PBS, sections were stained with fluorophore-conjugated secondary antibodies (Alexa Fluor-488 goat anti-rabbit /Alexa Fluor-488 goat anti-mouse (1:400, A11008/A11001, Thermo Fisher) or Alexa Fluor-633 goat anti-rat (1:400, A21094, Thermo Fisher)) in PBS or 0.1% Triton X-100-PBS at room temperature for 1-1.5 h. Counterstaining was done with Hoechst reagent (Sigma-Aldrich, 1:1000 in PBS). Confocal laser scanning microscopy was performed using a Zeiss LSM780 confocal microscope equipped with a 40x objective. Images files were exported and further analysis was carried out using NIH Image J software.

*Surface morphology imaging*

To visualize potential deformations and assess the overall shapes of mouse brains, the entire organ was imaged in a Bioptonics 3001 OPT Scanner was used. Samples were imaged using autofluorescence f-OPT [25] and reflected light [26]. Four hundred images with a 0.9° angle pitch were acquired to image a full rotation of the sample. Consequently, 3D volumes were reconstructed using NRecon software (version 1.7.1.6; Bruker) and visualized with Arivis (version 2.12.5; Rostock).

*Single-cell RNA sequencing*

Single-cell suspensions were prepared as described in the flow cytometry section. Male mice aged between 12-16 weeks and from the same litter were used. Live $CD11b^+CD45^+$ and either $CD4^+CD45^+CD11b^-$ or $TCR\beta^+CD45^+CD11b^-$ cells were sorted using a BD FACSAria III, and suspended in 0.04% BSA-PBS. Post-sorting, the cell number and viability were confirmed using a LUNA-FL dual fluorescence cell counter (Logos Biosystems). For each experiment, approximately 8,700 cells were added to each channel for a targeted cell recovery of 5,000 cells. Post-cell count and quality control, the samples were immediately loaded onto the 10X Genomics Chromium Controller and library preparation performed using the Single Cell 3' Kit v3, according to manufacturer's instructions. Library quality was checked at the recommended points using a Qubit 2 Fluorometer (Thermo Fisher) and a Bioanalyzer HS DNA kit (Agilent). Libraries were sequenced on an Illumina Novaseq 6000 or Illumina HiSeq platform using the recommended paired-end sequencing workflow (v3 read parameters, 28-8-0-91 cycles). On average, libraries were sequenced to a depth of 50,000 reads per cell.

Data were preprocessed with Cell Ranger V.3.1 ($\alpha CamKII^{IL2}$ dataset) or V6.0 (PHP.*GFAP*-IL2 dataset) from 10x Genomics. The resulting count matrices, showing the number of transcripts (UMIs) for each gene in a given cell, were analyzed with R V.3.6.3 (www.r-project.org) [27] and Seurat (https://satijalab.org/seurat/) V.3.1.5 [28] ($\alpha CamKII^{IL2}$ dataset), or V.4.0.1 and V.4.0.5 (PHP.*GFAP*-IL2 dataset), following the standard pipeline with default parameters, unless stated otherwise. Prior to analysis, the data was filtered based on different quality metrics calculated to only include *bona fide* single cells of high quality. Genes detected in less than five cells were filtered out. Low-quality cells or empty droplets (identified as those with less than 200 genes) and cells with less than 500 UMI counts were also filtered out. Finally, libraries with extensive mitochondrial reads, indicative of dying cells, were filtered out. The feature expression measurements for each cell within the combined datasets were normalized by the total expression and log-transformed. Before clustering, unwanted variation due to UNI number and mitochondrial gene expression was removed. A linear transformation (‘scaling’) of gene expression was also performed, normalizing across cells for variations in gene expression.

For the identification of various cell populations, dimension reduction approaches were applied on gene expression data using the Seurat analysis package. Initially, linear dimension reduction was applied in the form of Principal Component Analysis (PCA), using the PCElbowPlot() function, to obtain the principal components, followed by dimension reduction approaches, based on similarities in the expression data. tSNE (t-distributed stochastic neighbor embedding) and UMAP (uniform manifold approximation and projection) reduction was used for non-linear dimensionality reduction on the subset of PCAs representing the most variation in the gene expression data (determined via elbow plots). The A k-nearest neighbor algorithm was then applied to the projection to generate the shared nearest neighbor graph, which was used to generate the clusters using FindClusters() function with the Louvain algorithm, with a resolution of 0.4 and 1,000 iterations. Thus, cells which are similar in gene expression cluster together in these "communities". Once clusters were generated, expression of known marker genes was used to assign cell-type identity. Comparison of cell proportions were calculated using t-test with Bonferroni correction. Pathway analysis was performed using GAGE V.2.40.2 [29], Pathview V.1.30.1 [30], and clusterProfiler V.3.18 [31]. The full *αCamKII*$^{IL2}$ analysis code is available in **Supplementary Resource 2**, with the data available on GEO as dataset GSE153427. The full PHP.*GFAP*-IL2 analysis code is available in **Supplementary Resource 3**, with the data available on GEO as dataset GSE179176.

*High-Sensitivity mouse IL-2 Immunoassay*

Serum was obtained from whole blood by incubation of blood at room temperature for 30 min followed by centrifugation at 2000 x g for 10 min. Serum samples were diluted (1/40) in assay dilution buffer (Life Technologies). Tissue samples (5 mg) were placed in 300 µl protein quant sample lysis buffer (Life Technologies) containing protein inhibitor cocktail (Life Technologies). Tissues were homogenised in a FastPrep instrument (MP Biomedicals) with Lysing Matrix D, according to the manufacturer’s recommendation, and then incubated on a shaker for 20 min at 4°C. The lysate obtained was centrifuged at 16000 x g for 1 min at 4°C. IL2 levels from tissue lysates and serum were examined using a ProQuantum High-

Sensitivity mouse IL-2 immunoassay, according to the manufacturer's instructions (Life Technologies).

*Functional imaging in acute brain slices*

Recordings were done in (i) αCamKII$^{IL2}$ mice and littermate controls, or (ii) age-matched AAV injected mice (3-4 months old). Preparation of acute brain slices was adapted from a published protocol [32]. Briefly, animals were anesthetized using intraperitoneal administration of nembutal (50 mg/kg). Transcardial perfusion was performed using 20 ml of ice-cold N-methyl-D-glucamine-based artificial cerebrospinal fluid dissection solution (NMDG-ACSF), containing (in mM): 93 NMDG, 2.5 KCl, 1.25 $NaH_2PO_4$, 30 $NaHCO_3$, 10 $MgSO_4$, 0.5 $CaCl_2$, 20 HEPES, 25 D-glucose, 5 L-ascorbic acid, 2 thiourea, 3 sodium pyruvate, 10 N-acetyl-L-cysteine. The solution was adjusted to 305–310 mOsm/l, pH 7.4 (HCl) and bubbled in 95% $O_2$/5% $CO_2$ gas for 20 min before use and throughout the experiment. Following decapitation, the brain was swiftly removed. Coronal slices containing the visual cortex (350 µm thick) were obtained using a VT1200s vibratome (Leica Biosystems). Slices were further cut along the midline to separate the cerebral hemispheres, and placed in a chamber containing NMDG-ACSF at 33°C. Slices were maintained under these conditions for 25 min, with the controlled reintroduction of $Na^+$ achieved by gradual addition of 2 M NaCl to the chamber. Slices were then transferred to a storage chamber containing room temperature holding ACSF (in mM): 92 NaCl, 2.5 KCl, 1.25 $NaH_2PO_4$, 30 $NaHCO_3$, 2 $MgSO_4$, 2 $CaCl_2$, 15 D-Glucose, 20 HEPES, 5 L-ascorbic acid, 2 thiourea, 3 sodium pyruvate, 10 N-acetyl-L-cysteine (pH 7.4 HCl; 95% $O_2$/5% $CO_2$ gas; osmolarity 305–310 mOsm/l), until they were used for dye loading and recordings. For recordings, we used recording ACSF (in mM): 1.24 NaCl, 2.5 KCl, 1.25 $NaH_2PO_4$, 26 $NaHCO_3$, 2 $MgSO_4$, 2 $CaCl_2$, 15 D-Glucose (pH 7.4 HCl; 95% $O_2$/5% $CO_2$ gas; osmolarity 305–310 mOsm/l). For imaging, slices were transferred to a specialized recording chamber and superfused with normal ACSF (including pharmacological reagents where appropriate) at 22°C.

To identify astrocytes, cells were labeled with sulforhodamine 101 (SR101). To avoid potential issues associated with high levels of SR101 loading (such as induction of seizure like activity [33]), slices were incubated for 20 min in a six-well culture dish containing 1 µM SR101 (Sigma-Aldrich) in holding ACSF at 33°C. Next, slices were loaded in a solution containing Fluo4-AM. The dye was supplied as a 50 µg ampoule (Thermo Fisher) and was solubilized using a mixture of 7 µl dimethyl sulfoxide (DMSO), 2 µl 20% Pluronic F-127 (Tocris Biosciences) in DMSO and 1 µl 0.5% Kolliphor EL (Sigma-Aldrich) in DMSO. The ampoule was then incubated at 41°C with constant agitation (1400 RPM) for 15 min using a thermomixer. Concentrated Fluo4-AM was then added to a well containing 3 ml of standard ACSF, giving a final concentration of 15.2 µM of Fluo4-AM. Slices were loaded in this solution for 45–60 min at 35°C. At the end of this period, excess AM dye was removed by washing in room temperature holding ACSF for at least 1 hour prior to imaging.

All functional imaging recordings followed the same protocol. A field of view containing the primary visual cortex was chosen. First, baseline activity was recorded for 60s. Next, (R)-(-)-phenylephrine hydrochloride, 50 µM (PHE, Tocris Bioscience) was bath applied. Slices were imaged over a total period of 300 s. Live imaging of cells in acute slices was performed using a two-photon imaging system (VIVO 2-Photon platform, Intelligent Imaging Innovations GmbH), equipped with a tunable multiphoton laser (MaiTai laser, Spectra-Physics). Imaging was performed using a Zeiss Axio Examiner Z1, equipped with a W PlanApochromat 20×/NA1.0 objective. To excite both SR101 and Fluo4, the excitation wavelength was tuned to 840 nm. Signals were detected using two fast-gated GaAsP PMTs (Hamamatsu Photosensor Modules H11706), equipped with either a 525/40 nm or 612/69 nm emission filter. The system was further equipped with a 580 nm edge BrightLine® single-edge imaging-flat dichroic beamsplitter (Semrock FF580-FDi01-28x38). Images were 512 × 512 pixels in size and acquired at a frequency between 3 and 3.5 Hz. Acquisition was controlled using Slidebook 6 software (Intelligent Imaging Innovations GmbH). Laser power was limited to a maximum of 30 mW at the specimen. The focal plane used was usually 30–100 µm deep within the slice.

Images were initially processed using Fiji software with standard plugins to correct for image drift and noise. First, a sum projection of all frames was used to generate one image of the SR101 channel, which was then used to draw a region of interest (ROI) surrounding astrocyte cell bodies. The average fluorescence for each ROI per frame of the Fluo4 channel was then measured and exported to MATLAB (The Mathworks). Next, we used custom-written scripts to evaluate the relative variations in intracellular $Ca^{2+}$, estimated as changes in Fluo4 signal relative to the baseline ($\Delta F/F0$). The baseline (F0) was defined for each ROI to be the average fluorescence before phenylephrine was added to the recording chamber. We calculated the peak amplitude of the response to PHE, as well as the total area under the curve (AUC) of the ΔF/F0 trace using custom MATLAB scripts.

*Multi-electrode array electrophysiology*

LTP recordings were done in littermate αCamKII$^{IL2}$ and control mice, or age-matched AAV-injected mice (3-4 months old). For tissue preparation, mice were anesthetized with isoflurane and decapitated. Brains were rapidly removed and 300 µm thick parasagittal brain slices prepared using a Leica VT1200 vibratome. Slicing was performed in a sucrose-based cutting solution (ACSF) containing (in mM): 87 NaCl, 2.5 KCl, 1.25 $NaH_2PO_4$, 10 glucose, 25 $NaHCO_3$, 0.5 $CaCl_2$, 7 $MgCl_2$, 75 sucrose, 1 kynurenic acid, 5 ascorbic acid, 3 pyruvic acid (pH 7.4 (HCl) ; 95% $O_2$/5% $CO_2$ gas). Slices were allowed to recover at 34°C for 35 min, and then maintained at room temperature in the same solution for at least 30 min before using. For recordings, slices were placed onto a multielectrode array (MEA 2100, Multi Channel Systems) and continuously perfused with artificial cerebrospinal fluid (aCSF) solution containing (in mM): 119 NaCl, 2.5 KCl, 1 $NaH_2PO_4$, 11 glucose, 26 $NaHCO_3$, 4 $MgCl_2$ and 4 $CaCl_2$ (pH 7.4 (HCl) ; 95% $O_2$/5% $CO_2$ gas; 34°C). Field excitatory post-synaptic potentials (fEPSPs) were recorded from Schaffer collateral-CA1 synapses by stimulating and recording from the appropriate (visually identified) electrodes. Input-output curves were recorded for each slice by applying single-stimuli ranging from 500 to 2750 mV with 250 mV increments. A stimulus strength that corresponds to 35% of the maximal response in the input-output curve was used for recordings. For long-term potentiation (LTP) experiments, stable fEPSPs were recorded for 30 minutes to establish a baseline. Next, we applied three high frequency trains (100 stimuli; 100 Hz) with 5 minutes intervals. Subsequently, post-LTP fEPSPs were measured every 5 minutes (average of three consecutive stimuli at 15 seconds intervals) for 55 minutes. Recordings were processed and analyzed using Multi Channel Experimenter software (Multi Channel Systems).

*Quantification of blood-CSF barrier and blood-brain barrier permeability*

Blood-CSF barrier and blood-brain barrier permeability were determined as previously described [7]. Briefly, 4 kDa FITC-dextran (Sigma) was injected intravenously 1 h before CSF collection. CSF was obtained from the fourth ventricle using the cisterna magna puncture method. Subsequently, mice were perfused with 0.2% heparin-PBS and brain tissue was isolated. CSF samples were diluted 100-fold in sterile PBS, and blood-CSF barrier leakage was determined by measurement of fluorescence at $\lambda_{ex}$ 485 nm; $\lambda_{em}$ 520 nm. Brain samples were cut into small pieces, incubated overnight at 37°C in formamide while shaking. Supernatant was collected after centrifugation for 15 min at maximum speed. Brain fluid was diluted 2-fold in sterile PBS, and blood-brain barrier leakage was determined by measurement of fluorescence at $\lambda_{ex}$ 485 nm; $\lambda_{em}$ 520 nm.

*Statistics*

Comparisons between two groups were performed using unpaired two-tailed Student's t tests. Post hoc Holm's or Sidak's multiple comparisons tests were performed, when required. Two-way ANOVA was used when appropriate. Non-parametric testing was performed when data was not normally distributed (QQ plot for visual check and Shapiro-Wilk normality test on pooled residuals). The value of n reported within figure legends represents the number of animals, unless otherwise specified. Values are represented as mean ± SEM, with differences considered significant when $p < 0.05$.

tSNE, FlowSOM and heatmap analysis were performed in R (version 3.6.2) using an in-house script (manuscript in preparation). FlowSOM clusters are formed based on multi-marker similarity in a non-supervised manner. Clusters were annotated based on post-clustering comparison of marker expression, aligning the unique marker profile of each cluster to literature-based nomenclature. Key annotations for Treg clusters included naïve ($CD62L^{hi}CD44^{low}$), activated ($CD62L^{low}CD44^{high}$), resident (activated, plus enriched for expression of CD69, CD103, KLRG1, ST2) and pTreg ($Nrp1^{-}$), with additional clusters annotated based on the unique marker distribution. Differences between tSNE plots were calculated following the same approach as in the tSNE algorithm (manuscript in preparation). From these point probabilities, the distribution of cross-entropy in the tSNE space relative to the original space was obtained per plot. Then all pair-wise comparisons between plots were evaluated with Kolmogorov-Smirnov tests on the differences between the cross-entropy distributions. Resulting p-values were corrected with the Holm method. Dendrograms were obtained from hierarchical clustering, using as distance the Kolmogorov-Smirnov statistic (manuscript in preparation).

**Supplementary Resource 1. High resolution Treg localization in coronal sections of wildtype and αCamKII$^{IL2}$ mice.** Healthy perfused mouse brains from wildtype and αCamKII$^{IL2}$ mice were compared by immunofluorescent confocal imaging. CD4 (green), Foxp3 (red), laminin α4 (vascular basement membrane, white) and DAPI (blue). Left, wildtype mouse. Right, αCamKII$^{IL2}$ mouse. High resolution allows zooming and panning for localization of Tregs across the entire coronal section.

**Supplementary Resource 2. Code for single-cell RNA-seq analysis for wildtype vs *αCamKII*$^{IL2}$ mice.** Code is provided in .pdf and .Rmd formats.

**Supplementary Resource 3. Code for single-cell RNA-seq analysis for PHP.GFAP-GFP- vs PHP.GFAP-IL2-treated mice.** Code is provided in .pdf and .Rmd formats.

**Supplementary Video 1. CD4 T cells in wildtype brain.** 3D surface rendering of a wildtype perfused brain, stained for CD4 (green), Foxp3 (red), CD31 (white) and DAPI (blue). Representative video of a conventional CD4 T cell and a Treg, in the mid-brain region.

**Supplementary Video 2. Brain Tregs in αCamKII$^{IL2}$ brain.** 3D surface rendering of an αCamKII$^{IL2}$ perfused brain, stained for CD4 (green), Foxp3 (red), CD31 (white) and DAPI (blue). Representative video of a CD4 T cell cluster, consisting of two conventional CD4 T cell and four Tregs, in the mid-brain region.

**Supplementary Video 3. CD4 T cells in control brain.** 3D surface rendering of a PHP.GFAP-GFP-treated perfused brain, stained for CD4 (green), Foxp3 (red), CD31 (white) and DAPI (blue). Representative video of a conventional CD4 T cell and a Treg, within the mid-brain.

**Supplementary Video 4. Brain Tregs in PHP.GFAP-IL2-treated brain.** 3D surface rendering of a PHP.GFAP-IL2-treated perfused brain, stained for CD4 (green), Foxp3 (red), CD31 (white) and DAPI (blue). Representative video of a CD4 T cell cluster consisting of three Tregs, in the mid-brain region.

**Supplementary Figure 1. IL2 reporter expression in neurons.** Healthy perfused mouse brains from $IL2^{GFP}$ mice and non-transgenic controls assessed for GFP reporter expression by immunohistochemistry. **(A)** anti-GFP IL2 reporter (green), NeuN (red), GFAP (purple) and DAPI (blue). Single and combined channel confocal images of GFP-expressing NeuN+ cells in the mid-brain, compared to non-transgenic control. **(B)** anti-GFP IL2 reporter (green), Iba1 (red), GFAP (purple) and DAPI (blue). Single and combined channel confocal images of GFP-expressing cells in the mid-brain, compared to non-transgenic control. All images are of representative sections. Scale bar, 50 μm.

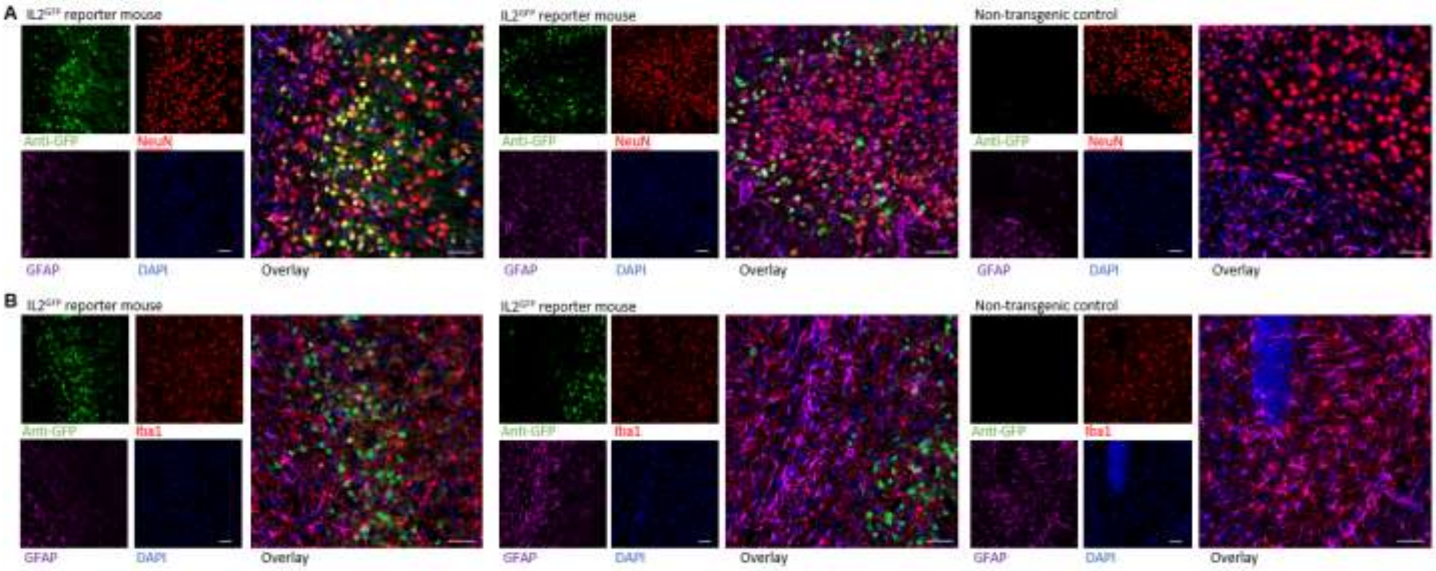

**Supplementary Figure 2. Brain regulatory T cell-specific effects of neuronal IL2 production. (A)** Healthy spleen from wildtype and Foxp3$^{IL2}$ mice were compared by high-dimensional flow cytometry (n = 4, 6). Frequency quantification of key markers (CD25. CD44. CD62L, CD103, CTLA4, Helios, ICOS, Ki67, KLRG1, Neuropilin1, PD1, ST2, Tbet) on Tregs in spleen. **(B)** Healthy perfused brain from wildtype and αCamKII$^{IL2}$ mice were compared by high-dimensional flow cytometry (n = 4, 3). Frequency quantification of key markers (CD25. CD44. CD62L, CD103, CTLA4, Helios, ICOS, Ki67, KLRG1, Neuropilin1, PD1, ST2, Tbet) on Tregs. **(C)** tSNE of blood, spleen and brain Tregs built on key markers (CD25. CD44. CD62L, CD103, CTLA4, Helios, ICOS, Ki67, KLRG1, Neuropilin1, PD1, ST2, Tbet). Colors indicate annotated FlowSOM clusters, with quantification in main Figure 1N. tSNE run on samples pooled post-acquisition, with quantification performed on individual samples (n = 4, 3). The residential cluster is characterized as CD25$^{hi}$CD69$^{+}$PD1$^{+}$CD103$^{+}$. **(D)** tSNE of total leukocytes from healthy perfused brain from wildtype and αCamKII$^{IL2}$ mice, built on lineage markers (CD4, CD8, NK1.1, CD44, CD62L, CD69, CD25, FoxP3) with quantification. tSNE run on samples pooled post-acquisition, with quantification performed on individual samples (n = 4, 3). **(E)** NK, CD4 and CD8 T cells, (left) as a proportion of CD45$^{+}$CD11b$^{-}$ cells in the brain of wildtype and αCamKII$^{IL2}$ mice, and (right) in absolute numbers, together with Tregs. **(F)** tSNE of brain CD4 conventional T cells built on key markers (CD62L, CD44, CD103, CD69, CD25, PD-1, Nrp1, ICOS, KLRG1, ST2, Ki67, Helios, T-bet, CTLA4). t-SNE run on samples pooled post-acquisition, with quantification performed on individual samples (n = 4, 3). Colors indicate annotated FlowSOM clusters, with quantification and **(G)** frequency of marker expression. **(H)** t-SNE of brain CD8 T cells built on key markers (CD62L, CD44, CD103, CD69, CD25, PD-1, Nrp1, ICOS, KLRG1, ST2, Ki67, Helios, T-bet, CTLA4). t-SNE run on samples pooled post-acquisition, with quantification performed on individual samples (n = 4, 3). Colors indicate annotated FlowSOM clusters, with quantification and **(I)** frequency of marker expression. **(J)** t-SNE of brain NK cells built on key markers (CD62L, CD44, CD103, CD69, CD25, PD-1, Nrp1, ICOS, KLRG1, ST2, Ki67, Helios, T-bet, CTLA4). t-SNE run on samples pooled post-acquisition, with quantification performed on individual samples (n = 4, 3). Colors indicate annotated FlowSOM clusters, with quantification and **(K)** frequency of marker expression.

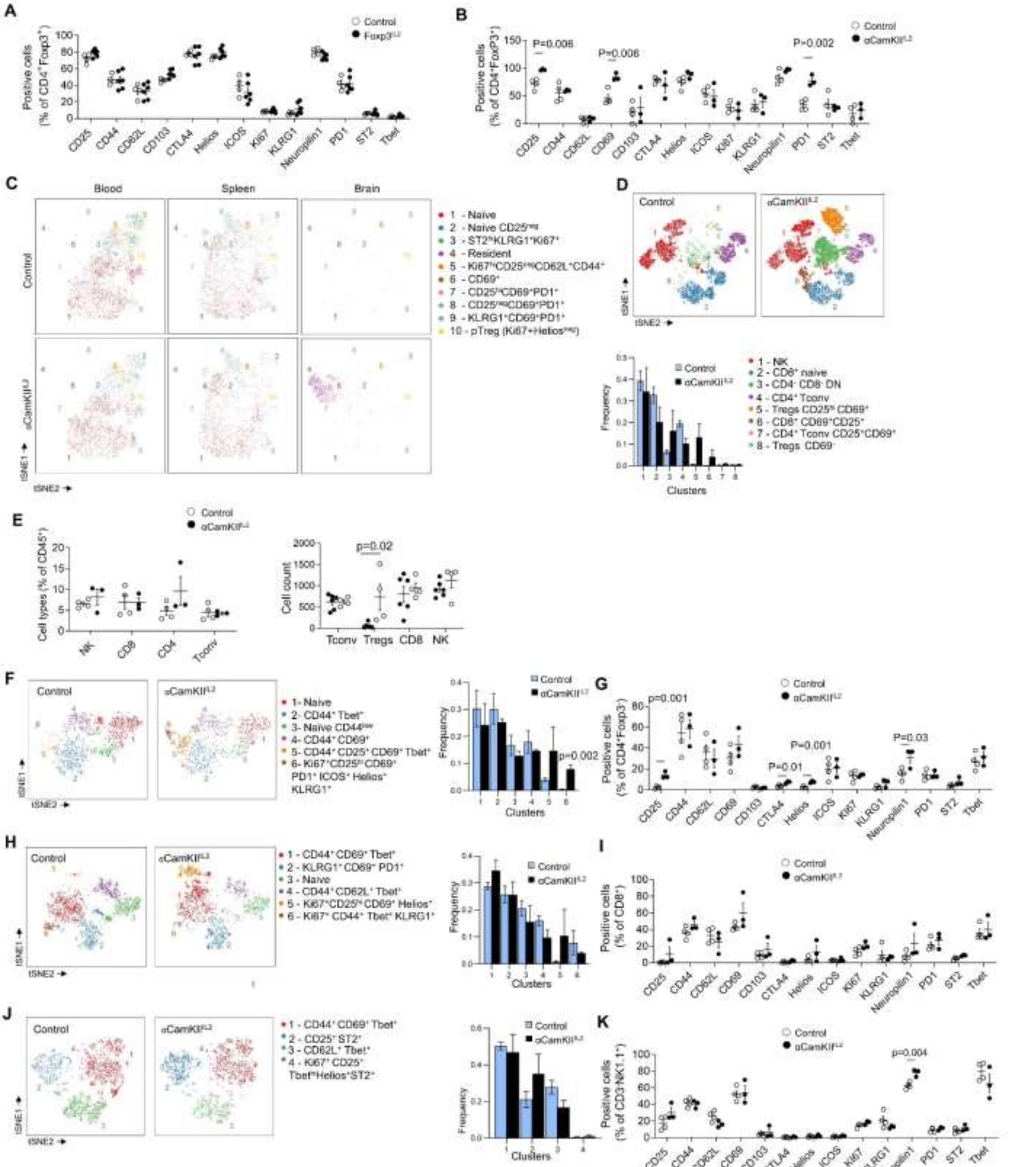
A
Control
Foxp3^IL2
Positive cells (% of CD4+Foxp3+)
CD25 CD44 CD62L CD103 CTLA4 Helios ICOS Ki67 KLRG1 Neuropilin1 PD1 ST2 Tbet
B
Control
αCamKII^IL2
Positive cells (% of CD4+FoxP3+)
P=0.006
P=0.006
P=0.002
CD25 CD44 CD62L CD69 CD103 CTLA4 Helios ICOS Ki67 KLRG1 Neuropilin1 PD1 ST2 Tbet
C
Blood
Spleen
Brain
Control
αCamKII^IL2
tSNE1
tSNE2
1 - Naive
2 - Naive CD25neg
3 - ST2hi KLRG1+Ki67+
4 - Resident
5 - Ki67hi CD25neg CD62L+CD44+
6 - CD69+
7 - CD25hi CD69+PD1+
8 - CD25neg CD69+PD1+
9 - KLRG1+CD69+PD1+
10 - pTreg (Ki67+Heliosneg)
D
Control
αCamKII^IL2
tSNE1
tSNE2
Frequency
Clusters
1 - NK
2 - CD8+ naive
3 - CD4- CD8- DN
4 - CD4+ Tconv
5 - Tregs CD25hi CD69+
6 - CD8+ CD69+CD25+
7 - CD4+ Tconv CD25+CD69+
8 - Tregs CD69-
E
Control
αCamKII^IL2
Cell types (% of CD45+)
NK CD8 CD4 Tconv
p=0.02
Cell count
Tconv Tregs CD8 NK
F
Control
αCamKII^IL2
tSNE1
tSNE2
1- Naive
2- CD44+ Tbet+
3- Naive CD44low
4- CD44+ CD69+
5- CD44+ CD25+ CD69+ Tbet+
6- Ki67+CD25hi CD69+ PD1+ ICOS+ Helios+ KLRG1+
Frequency
Clusters
p=0.002
G
Control
αCamKII^IL2
Positive cells (% of CD4+Foxp3-)
p=0.001
P=0.01
P=0.001
P=0.03
CD25 CD44 CD62L CD69 CD103 CTLA4 Helios ICOS Ki67 KLRG1 Neuropilin1 PD1 ST2 Tbet
H
Control
αCamKII^IL2
tSNE1
tSNE2
1 - CD44+ CD69+ Tbet+
2 - KLRG1+ CD69+ PD1+
3 - Naive
4 - CD44+ CD62L+ Tbet+
5 - Ki67+CD25hi CD69+ Helios+
6 - Ki67+ CD44+ Tbet+ KLRG1+
Frequency
Clusters
I
Control
αCamKII^IL2
Positive cells (% of CD8+)
CD25 CD44 CD62L CD69 CD103 CTLA4 Helios ICOS Ki67 KLRG1 Neuropilin1 PD1 ST2 Tbet
J
Control
αCamKII^IL2
tSNE1
tSNE2
1 - CD44+ CD69+ Tbet+
2 - CD25+ ST2+
3 - CD62L+ Tbet+
4 - Ki67+ CD25+ Tbethi Helios+ ST2+
Frequency
Clusters
K
Control
αCamKII^IL2
Positive cells (% of CD3-NK1.1+)
p=0.004
CD25 CD44 CD62L CD69 CD103 CTLA4 Helios ICOS Ki67 KLRG1 Neuropilin1 PD1 ST2 Tbet

**Supplementary Figure 3. Confocal identification of brain Tregs in αCamKII$^{IL2}$ mice.** Healthy perfused mouse brains from wildtype and αCamKII$^{IL2}$ mice were compared by immunofluorescent confocal imaging. CD4 (green), Foxp3 (red), CD31 (vasculature, white) and DAPI (blue). Single and combined channel representative images of CD4 T cells in the mid-brain, with close-up imaging of identified CD4 T cells. Scale bar, 10 µm.

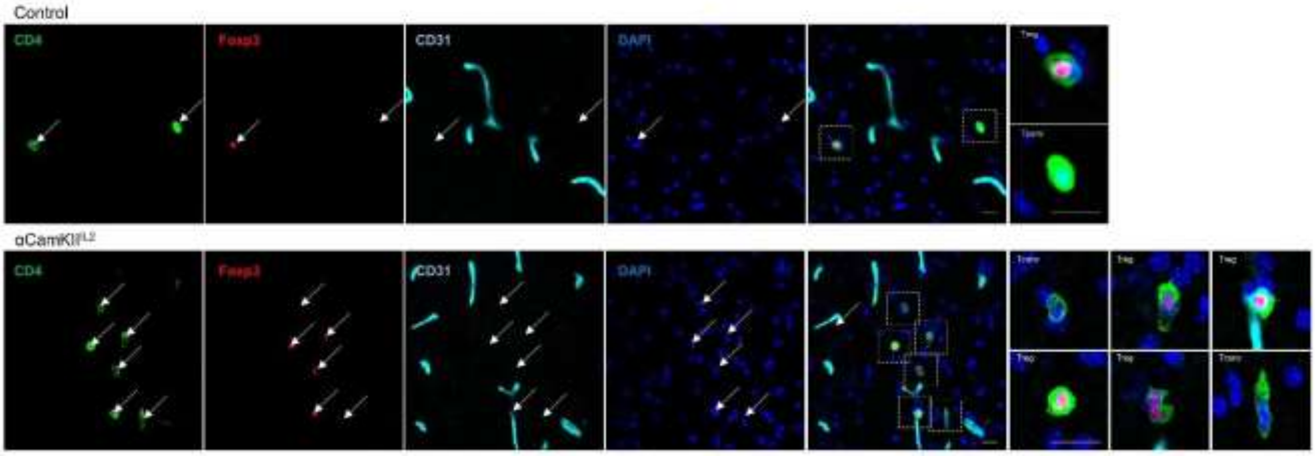

**Supplementary Figure 4. Brain Treg localization in sagittal sections of αCamKII$^{IL2}$ mice.** Healthy perfused mouse brains from **(A)** wildtype and **(B)** αCamKII$^{IL2}$ mice were compared by immunofluorescent confocal imaging. CD4 (green), Foxp3 (red), CD31 (vasculature, white) and DAPI (blue). Sagittal sections (top) were used to select large brain regions (main image). Arrow represent CD4 Tconv cells (blue) and Tregs (red). Letters indicate selected cells shown in insets. Scale bar main image, 100 µm. Scale bar insets, 10 µm.

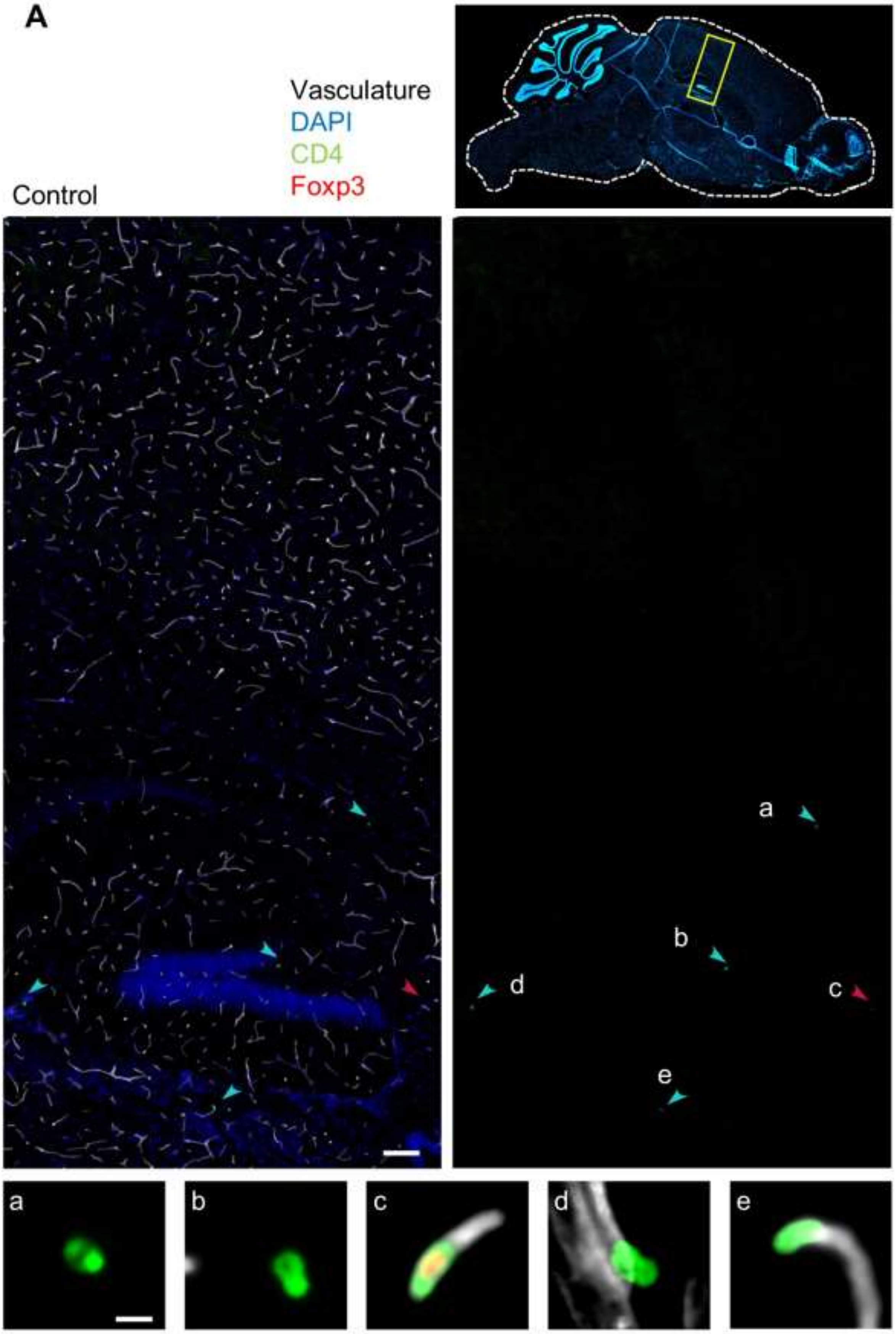

A
Vasculature
DAPI
CD4
Foxp3
Control
a
b
c
d
e

**B**

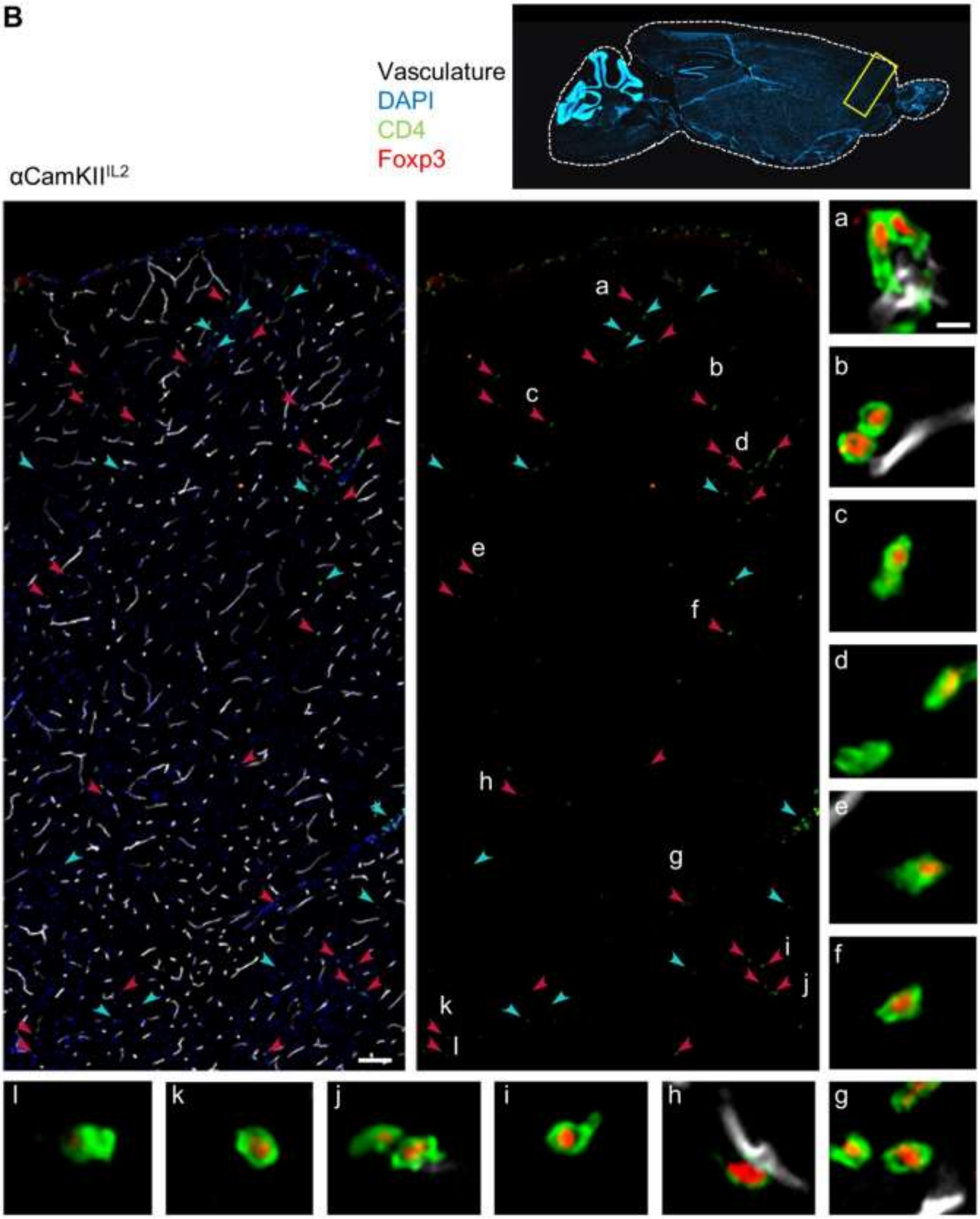

**Supplementary Figure 5. Brain Treg localization in coronal sections of αCamKII$^{IL2}$ mice.** Healthy perfused mouse brains from **(A)** wildtype and **(B)** αCamKII$^{IL2}$ mice were compared by immunofluorescent confocal imaging. CD4 (green), Foxp3 (red), laminin α4 (vascular basement membrane, white) and DAPI (blue). Entire coronal sections (left) were imaged, with the high resolution (single cell level) images available in Supplementary Resource 1. Five representative large regions were selected (middle), with identified CD4 Tregs in these regions annotated by meningeal/perivascular (orange arrow) versus parenchymal (white arrow) localization. Insets (right) visualize identified cells. Scale bar coronal section, 100 µm. Scale bar regional image, 100 µm. Scale bar insets, 10 µm.

A

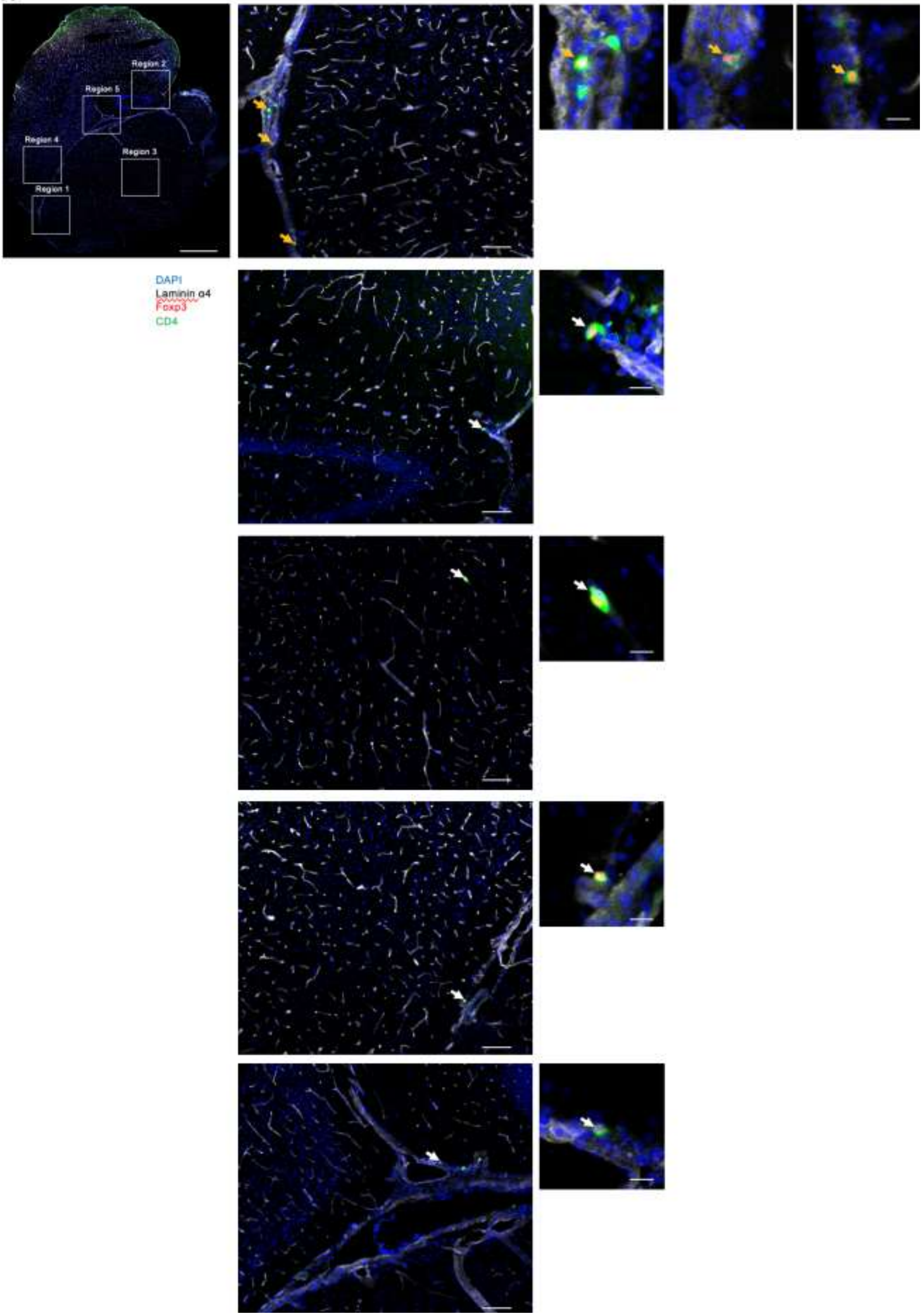
Region 2
Region 5
Region 4
Region 3
Region 1
DAPI
Laminin α4
Foxp3
CD4

**B**

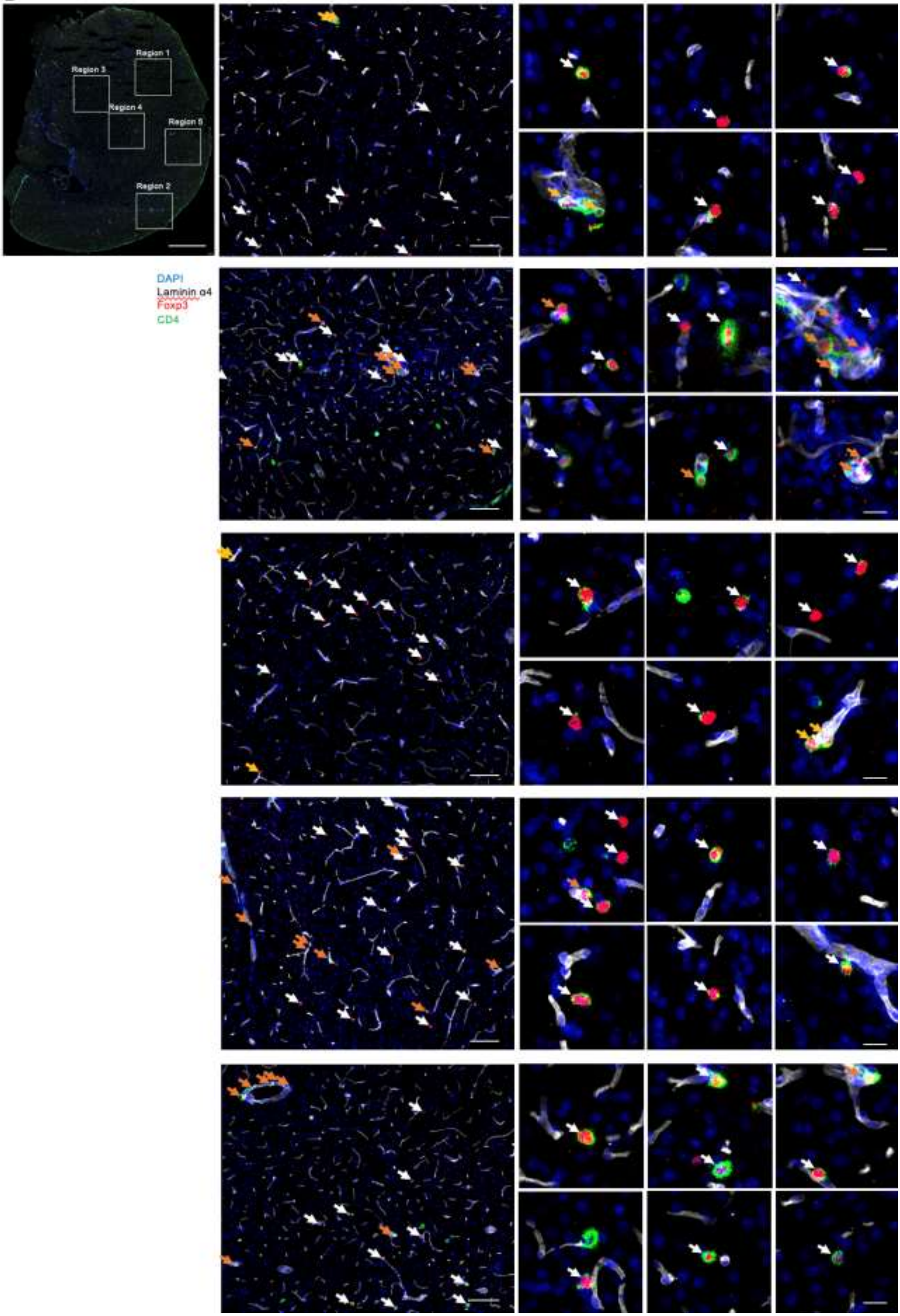
Region 1
Region 3
Region 4
Region 5
Region 2
DAPI
Laminin α4
Foxp3
CD4

**Supplementary Figure 6. Tregs in brain regions in αCamKII$^{IL2}$ mice.** Healthy perfused brain from wildtype and αCamKII$^{IL2}$ mice was dissected into major regions for flow cytometric analysis of Tregs (n = 5,3). **(A)** Frequency of Tregs within the CD4 population, and **(B)** absolute numbers of Tregs.

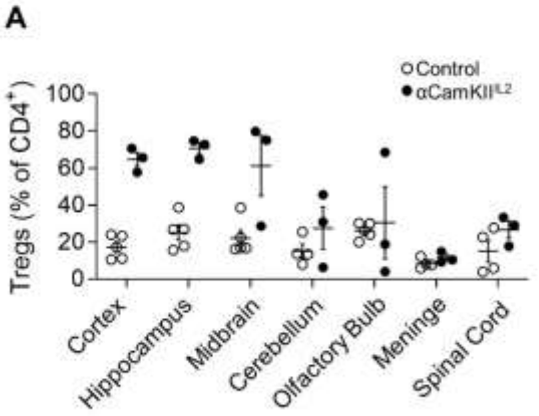


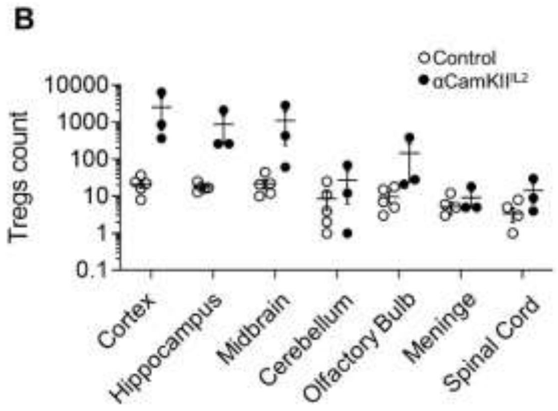

**Supplementary Figure 7. Synthetic expansion of brain regulatory T cells preserves the transcriptional profile.** Brain $CD11b^+$ cells, brain $CD45^+CD4^+$ cells and blood $CD45^+CD4^+$ positive cells were sorted from wildtype or $\alpha CamKII^{IL2}$ mice for 10X single-cell sequencing. Data processing and filtering identified 28,590 cells, of which 20,021 were classified as microglia, 6332 were classified as T cells and 2237 were classified as non-target leukocytes. **(A)** Using the full data, combining blood and brain per genotype, clusters were identified, with annotations **(B)** based on the expression of key lineage markers. Expression of key lineage markers visualized as colored single cells on tSNE plots. **(C)** Clusters containing $CD4^+$ conventional and regulatory T cells were reclustered, with expression of key lineage markers visualized as colored single cells on tSNE plots for use in cluster annotation. **(D)** Differential gene expression displayed for brain Tregs from wildtype vs $\alpha CamKII^{IL2}$ mice, **(E)** blood Tregs from wildtype vs $\alpha CamKII^{IL2}$ mice, **(F)** brain conventional T cells from wildtype vs $\alpha CamKII^{IL2}$ mice, **(G)** blood conventional T cells from wildtype vs $\alpha CamKII^{IL2}$ mice, or **(H)** blood vs brain Tregs. Vertical lines mark fold changes 0.4 and -0.4 and horizontal lines mark the adjusted P value of 0.05. Selected significant changes are annotated.

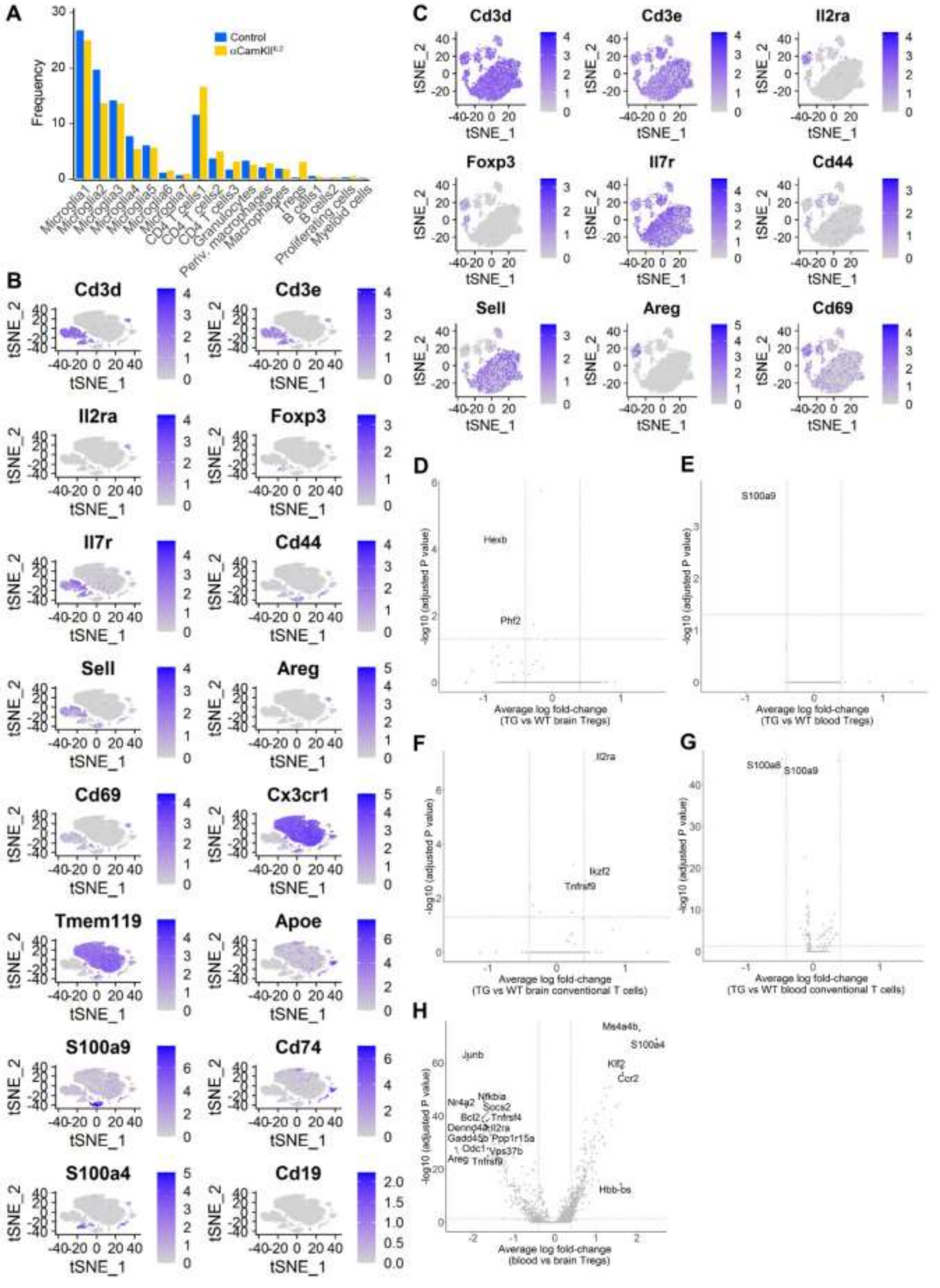
A
B
C
D
E
F
G
H
Control
αCamKII[IL2]
Frequency
Microglia1
Microglia2
Microglia3
Microglia4
Microglia5
Microglia6
Microglia7
CD4 T cells1
CD4 T cells2
CD4 T cells3
Granulocytes
Periv. macrophages
Macrophages
Tregs
B cells1
B cells2
Proliferating cells
Myeloid cells
Cd3d
Cd3e
Il2ra
Foxp3
Il7r
Cd44
Sell
Areg
Cd69
Cx3cr1
Tmem119
Apoe
S100a9
Cd74
S100a4
Cd19
tSNE_1
tSNE_2
Hexb
Phf2
Average log fold-change (TG vs WT brain Tregs)
-log10 (adjusted P value)
S100a9
Average log fold-change (TG vs WT blood Tregs)
Il2ra
Ikzf2
Tnfrsf9
Average log fold-change (TG vs WT brain conventional T cells)
S100a8
S100a9
Average log fold-change (TG vs WT blood conventional T cells)
Junb
Nr4a2
Nfkbia
Socs2
Bcl2
Tnfrsf4
Dennd4a
Il2ra
Gadd45b
Ppp1r15a
Odc1
Vps37b
Areg
Tnfrsf9
Ms4a4b
S100a4
Klf2
Ccr2
Hbb-bs
Average log fold-change (blood vs brain Tregs)

**Supplementary Figure 8. Normal long-term potentiation in αCamKII$^{IL2}$ mice.** Field excitatory post-synaptic potentials (fEPSPs) were recorded from Schaffer collateral-CA1 neuronal synapses in brain slices from wildtype and αCamKII$^{IL2}$ littermates. Input-output curves were recorded for each slice by applying single-stimuli ranging from 500 to 2750 mV with 250 mV increments. **(A)** Slope and **(B)** amplitude were analyzed (n=4,4). Long-term potentiation (LTP) was induced by applying three high frequency trains (theta-burst stimulation (TBS): 100 stimuli; 100 Hz) with 5 minutes intervals between trains. After baseline determination, fEPSPs were measured for 55 minutes. Changes in the **(C)** slope and **(D)** amplitude have been analyzed across time. Boxplots represent quantification of the baseline (left) and final LTP (right). Mean ± SEM (n=4,4).

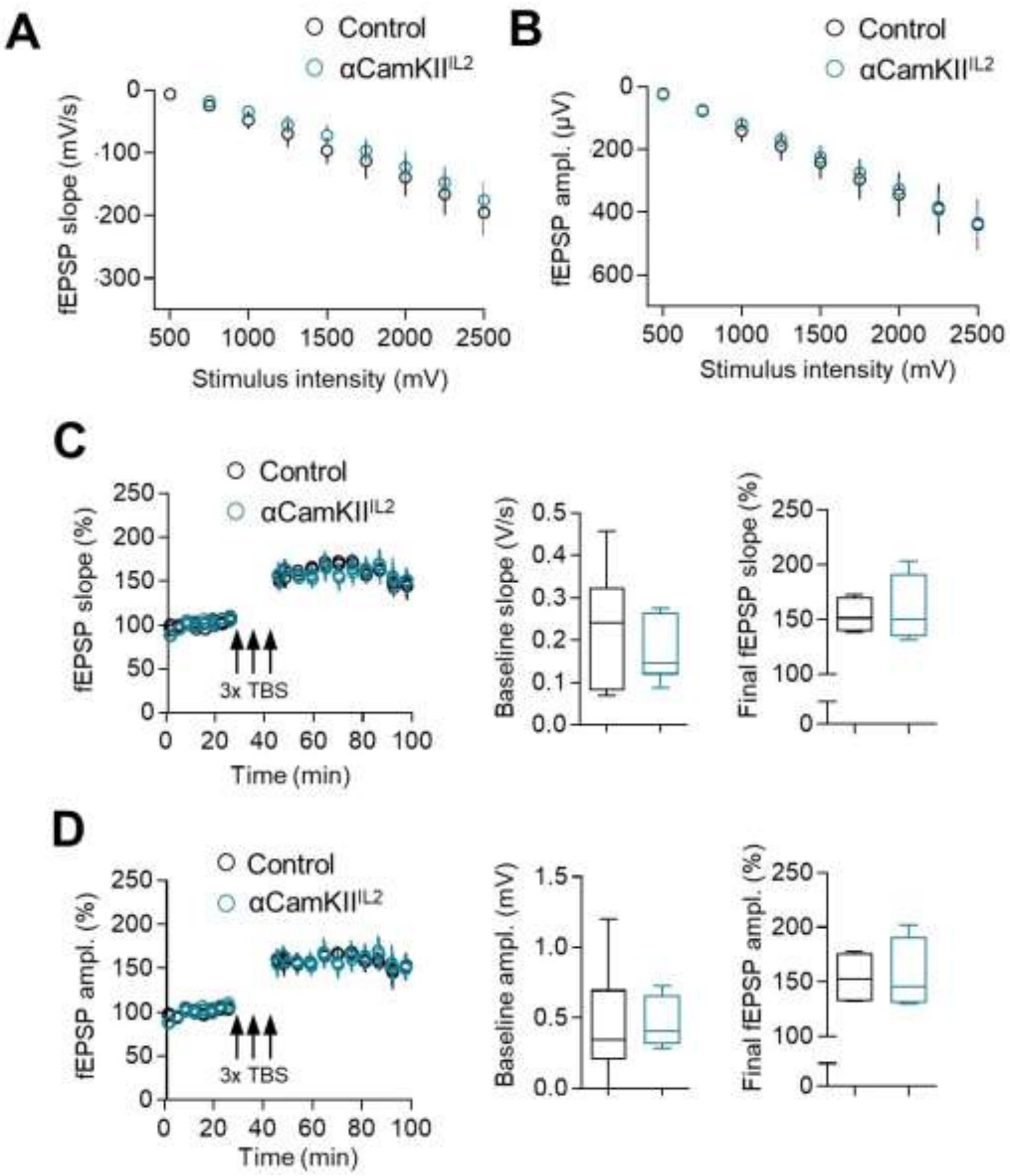

**Supplementary Figure 9. Normal behavior in mice with expanded brain regulatory T cells.** Behavioral assessment of αCamKII$^{IL2}$ and littermate control mice. **(A)** Time spent on the rod, average of 4 repeated tests of 300 seconds (n = 23, 17). **(B)** Open field, total distance moved and **(C)** time spent in the corners (n = 23, 16). **(D)** Nest building scoring (n = 24, 18). **(E)** Light-dark test latency to enter light zones and **(F)** time spent in the light zone (n = 20, 17). **(G)** Time immobile during forced swim test (n = 24, 16). **(H)** Sociability test trials to monitor the interaction with a stranger mouse (S) compared to an empty chamber (E) (n = 28, 18). **(I)** Freezing behavior over time during context acquisition conditioning (n = 28, 18). Mean ± SEM. **(J)** Contextual discrimination during generalization test. Mean ± SEM (n = 28, 18) **(K)** Spatial learning in the Morris water maze. Path length to finding the hidden platform (n = 28, 18), probe tests after 5 days and 10 days and after reversal learning (n = 28, 18). Mean ± SEM.

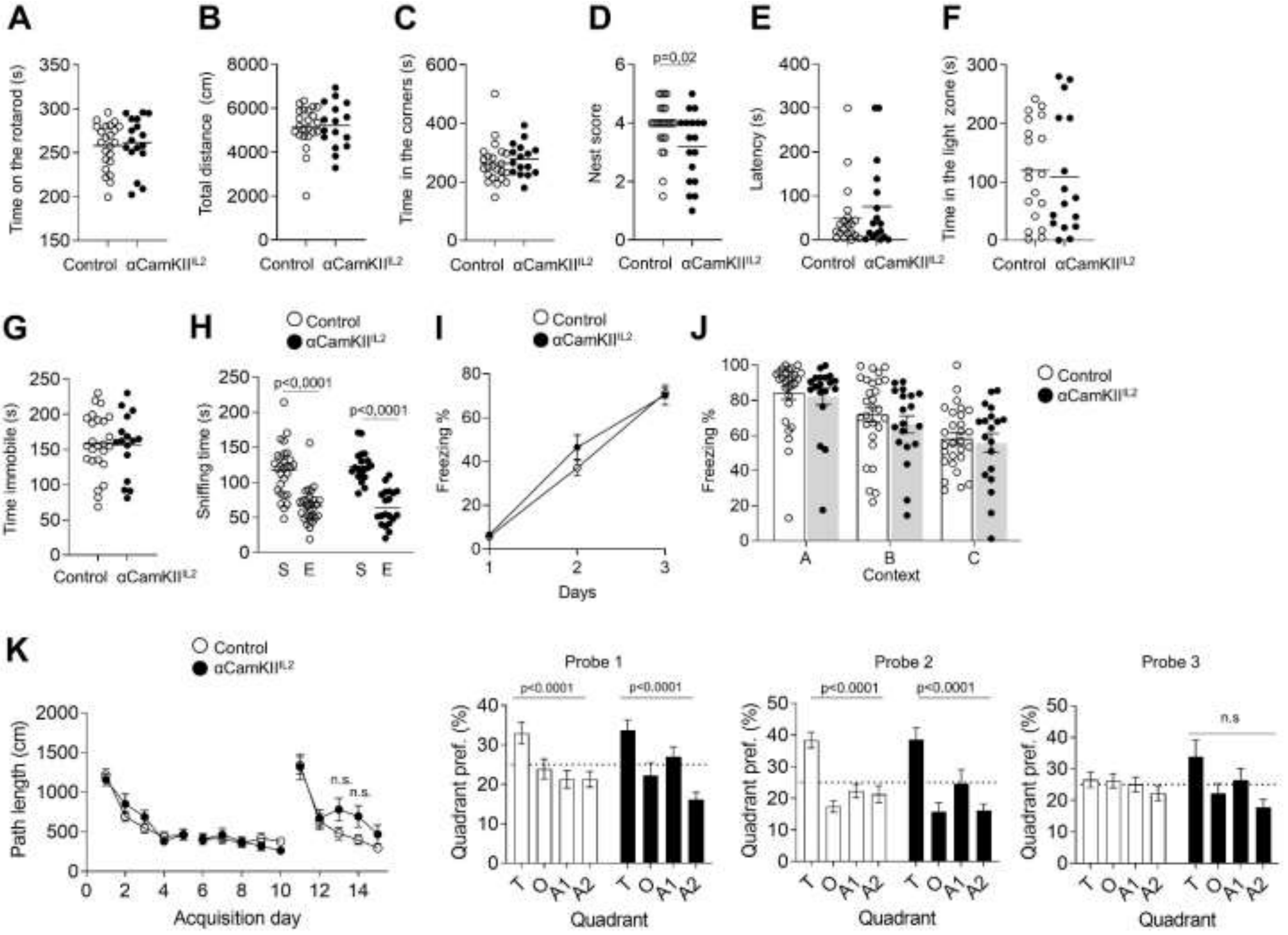

**Supplementary Figure 10. Intact microglial transcriptional signature in αCamKII$^{IL2}$ mice. (A)** tSNE projection of 20,021 microglial cells from wildtype or αCamKII$^{IL2}$ mice. Following alignment, each cell was grouped into clusters denoted by number and color. **(B)** tSNE projection of microglial cells labelled as originating from control or αCamKII$^{IL2}$ mice, with **(C)** the proportion of cells from control or αCamKII$^{IL2}$ mice belonging to the identified populations. **(D)** Volcano plots showing differentially expressed genes in wildtype vs αCamKII$^{IL2}$ microglial cells as annotated in panel A. Vertical lines mark fold changes 0.4 and -0.4 and horizontal line mark the adjusted P value of 0.05, genes above these thresholds are annotated. **(E)** Visualization of expression of key lineage markers (colored single cells) on tSNE plots. **(F)** Microglia from perfused brains of αCamKII$^{IL2}$ transgenic mice and littermate controls were assessed by high parameter flow cytometry. tSNE of microglia, gated on CD11b$^{+}$ CX3CR1$^{+}$ CD64$^{+}$ CD45$^{mod}$ Ly6G$^{-}$ (n = 4/group). Microglia clusters, based on FlowSOM expression analysis, were assessed for expression of CD64, MHCII, TGFβ, LAMP1, CD44, CD69, PDL1, ST2, Ki67, CD80 and IL1β using a heatmap and dendrogram. **(G)** Expression of key markers MHCII, PDL1 and CD80 visualized using tSNE projections. **(H)** Frequency of expression of MHCII, TNF, TGFβ, LAMP1, PDL1, Ki67, CD80 and IL1β in control αCamKII$^{IL2}$ microglia (n = 6, 11). Mean ± SEM.

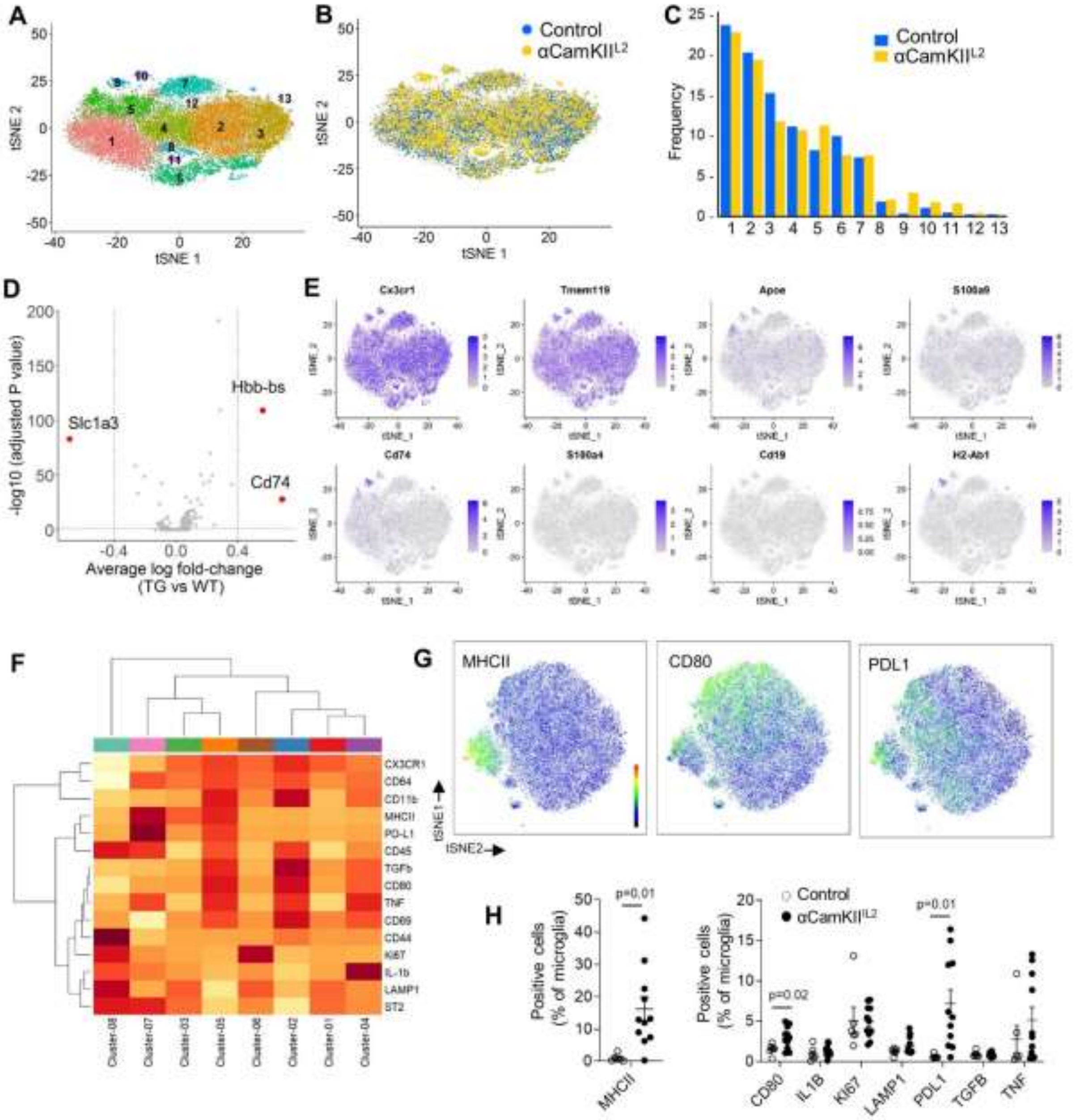
A
B
Control
αCamKII[L2]
C
Control
αCamKII[L2]
Frequency
D
Slc1a3
Hbb-bs
Cd74
-log10 (adjusted P value)
Average log fold-change
(TG vs WT)
E
Cx3cr1
Tmem119
Apoe
S100a9
Cd74
S100a4
Cd19
H2-Ab1
F
CX3CR1
CD64
CD11b
MHCII
PD-L1
CD45
TGFb
CD80
TNF
CD69
CD44
KI67
IL-1b
LAMP1
ST2
G
MHCII
CD80
PDL1
tSNE1
tSNE2
H
Positive cells
(% of microglia)
p=0.01
MHCII
Control
αCamKII[L2]
p=0.02
p=0.01
CD80
IL1B
KI67
LAMP1
PDL1
TGFB
TNF

**Supplementary Figure 11. Progressive neurological damage following traumatic brain injury.** Controlled cortical impact was performed to induce moderate TBI, with examination on days 1, 2, 3 and 7 post-TBI (n = 3, 3, 1, 3). **(A)** Macroscopic damage to the surface of the brain at the injury site, representative photo. Scale bar, 0.5 cm. **(B)** Representative immunofluorescence staining of the cortical tissue after controlled cortical impact. GFAP (astrocytes), Iba1 (microglia), DAPI (nuclei). Scale bars, 500 µm. **(C)** Measure of total integrated GFAP intensity in the cortical area adjacent to the impact site at 1, 2, 3 and 7 days post-TBI.

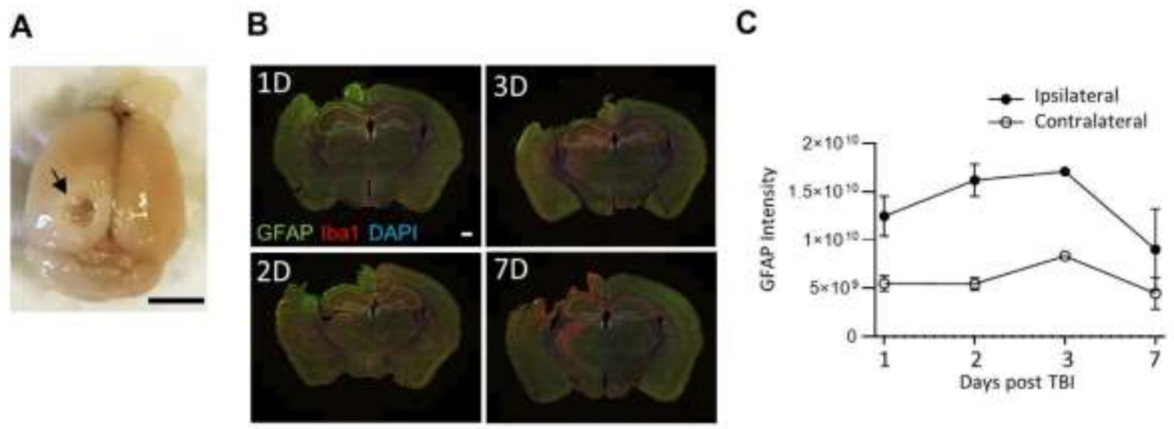

**Supplementary Figure 12. Normal peripheral influx following traumatic brain injury in αCamKII$^{IL2}$ mice.** Control (wildtype) littermates and αCamKII$^{IL2}$ mice were tamoxifen-treated at 6 weeks, and controlled cortical impacts to induce moderate TBI were given at 12 weeks. Mice were examined at 15 days post-TBI (n=4,4). TBI-induced perfused brains from littermate control and αCamKII$^{IL2}$ mice were compared by high-dimensional flow cytometry. **(A)** Absolute number of CD4, Tregs, CD8 and γδ T cells. **(B)** Expression of CD25, CD44, CD69, Ki67 and PDL1 markers, or **(C)** amphiregulin (Areg), IL10 and IL17 within the CD4 conventional T cell population. **(D)** Expression of CD25, CD44, CD69, Ki67 and PDL1 markers, or **(E)** amphiregulin, IL10 and IL17 within the Treg population. Mean ± SEM.

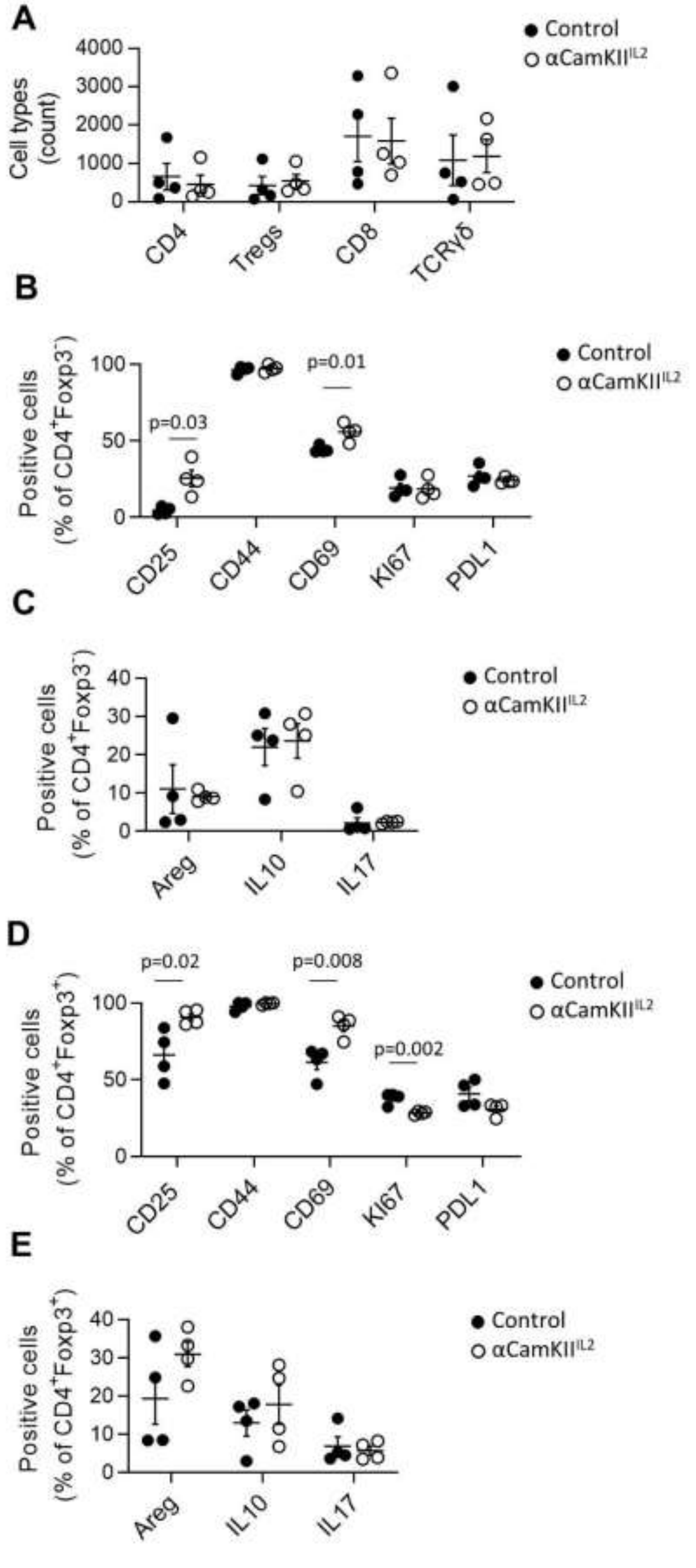
A
Control
αCamKII^IL2
Cell types (count)
4000
3000
2000
1000
0
CD4
Tregs
CD8
TCRγδ
B
Positive cells (% of CD4+Foxp3-)
p=0.03
p=0.01
100
50
0
CD25
CD44
CD69
KI67
PDL1
Control
αCamKII^IL2
C
Positive cells (% of CD4+Foxp3-)
40
30
20
10
0
Areg
IL10
IL17
Control
αCamKII^IL2
D
p=0.02
p=0.008
p=0.002
Positive cells (% of CD4+Foxp3+)
100
50
0
CD25
CD44
CD69
KI67
PDL1
Control
αCamKII^IL2
E
Positive cells (% of CD4+Foxp3+)
40
30
20
10
0
Areg
IL10
IL17
Control
αCamKII^IL2

**Supplementary Figure 13. PHP.*GFAP*-IL2 expands regulatory T cells in the brain without impacting draining lymph nodes.** Mice were treated with PHP.*GFAP*-GFP or PHP.*GFAP*-IL2 and assessed for Treg numbers by flow cytometry of perfused mice (n=4-5, 4-9). **(A)** Frequency of Tregs, as a proportion of CD4 T cells in the superficial cervical lymph nodes, and **(B)** deep cervical lymph nodes. **(C)** Absolute number of Tregs in superficial cervical lymph nodes, and **(D)** deep cervical lymph nodes. **(E)** Frequency and **(F)** absolute number of Tregs in the pia mater, 14 days after PHP.*GFAP*-GFP or PHP.*GFAP*-IL2 treatment (n=5,5). **(G)** Blood, spleen and perfused mouse brain from PHP.*GFAP*-GFP and PHP.*GFAP*-IL2-treated mice were compared by high-dimensional flow cytometry for Treg numbers (n=11-12/group). **(H)** Perfused organs from PHP.*GFAP*-GFP and PHP.*GFAP*-IL2-treated mice were compared by flow cytometry for Treg frequency (n=5/group). mLN, mesenteric lymph nodes; SC, spinal cord; IEL, intraepithelial leukocytes; LPL, lamina propria leukocytes; PP, Peyer's Patch.

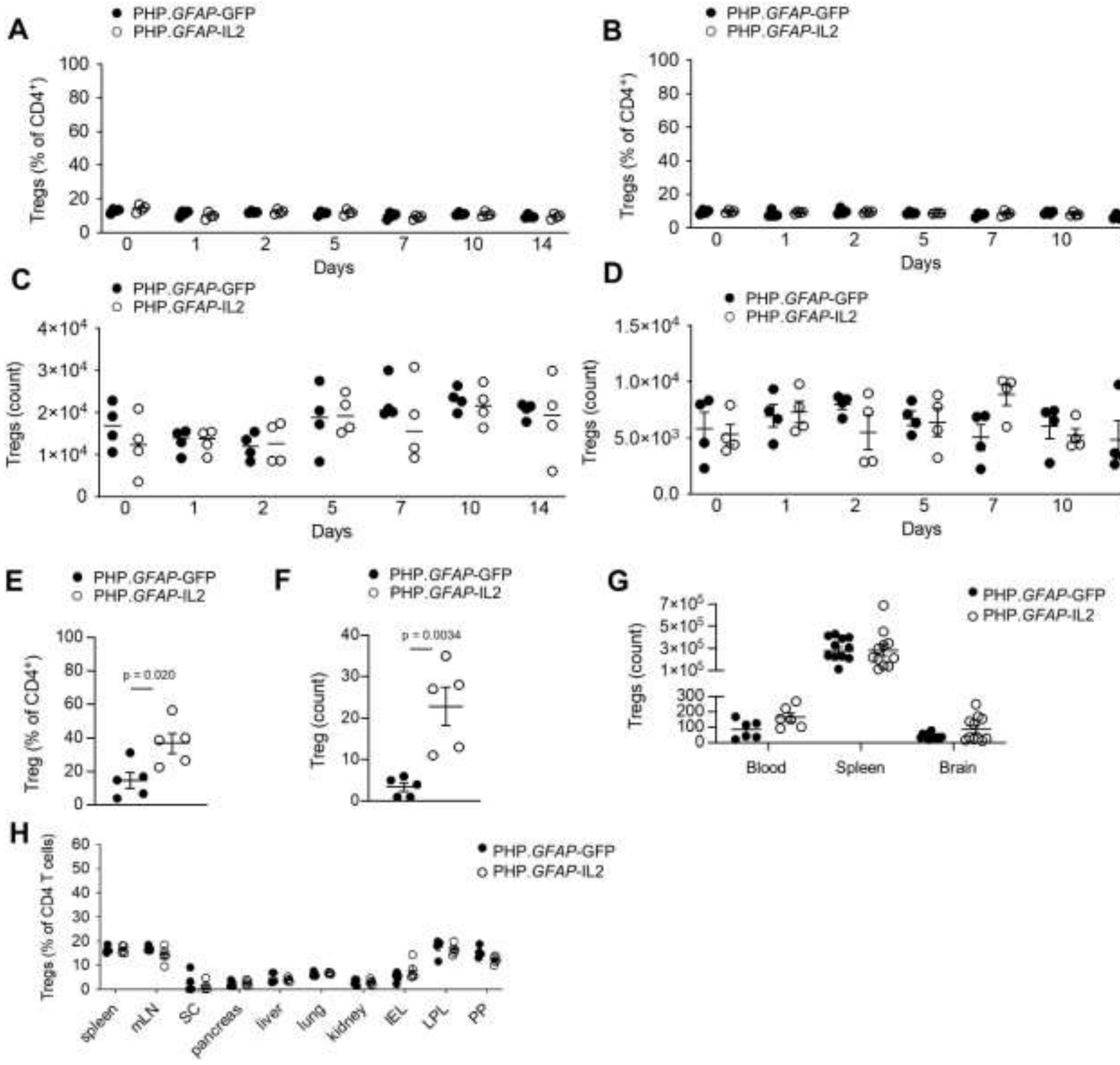
A
PHP.GFAP-GFP
PHP.GFAP-IL2
Tregs (% of CD4+)
Days
B
C
Tregs (count)
D
E
Treg (% of CD4+)
p = 0.020
F
Treg (count)
p = 0.0034
G
Tregs (count)
Blood
Spleen
Brain
H
Tregs (% of CD4 T cells)
spleen
mLN
SC
pancreas
liver
lung
kidney
IEL
LPL
PP

**Supplementary Figure 14. Confocal identification of brain Tregs in PHP.*GFAP*-IL2-treated mice.** Healthy perfused mouse brains from wildtype mice treated with PHP.*GFAP*-GFP or PHP.*GFAP*-IL2 were compared by immunofluorescent confocal imaging. CD4 (green), Foxp3 (red), CD31 (white) and DAPI (blue). Single and combined channel representative images of CD4 T cells in the mid-brain, with close-up imaging of identified CD4 T cells. Scale bar, 10 µm.

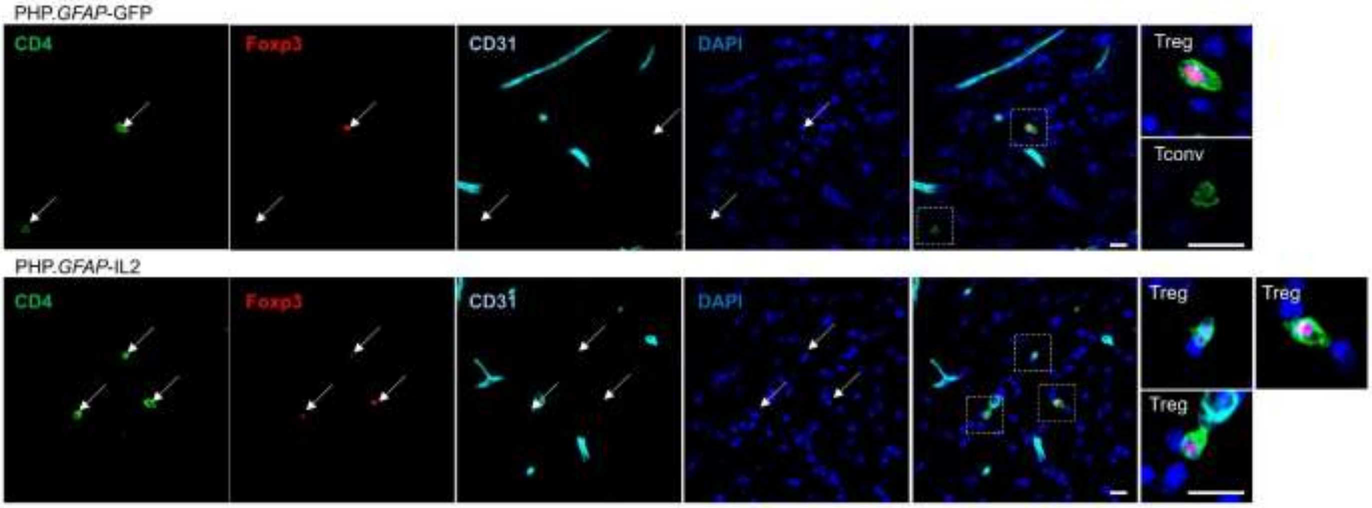

**Supplementary Figure 15. Off-target effects of PHP.*GFAP*-IL2 treatment are not observed. (A)** Healthy perfused brain, spleen and blood from PHP.*GFAP*-GFP control and PHP.*GFAP*-IL2-treated mice were compared by high-dimensional flow cytometry (n = 3, 5). tSNE plots of leukocytes built on lineage markers (CD4, CD8, NK1.1, CD44, CD62L, CD69, CD25, Foxp3) with quantification of major populations. **(B)** Absolute numbers of key leukocyte populations. **(C)** tSNEs of brain CD4 conventional T cells built on key markers (CD62L, CD44, CD103, CD69, CD25, PD-1, Nrp1, ICOS, KLRG1, ST2, Ki67, CTLA4). Colors indicate annotated FlowSOM clusters, with quantification and **(D)** frequency of marker expression. **(E)** tSNEs of brain CD8 T cells built on key markers (CD62L, CD44, CD103, CD69, CD25, PD-1, Nrp1, ICOS, KLRG1, ST2, Ki67, CTLA4). Colors indicate annotated FlowSOM clusters, with quantification and **(F)** frequency of marker expression. **(G)** tSNEs of brain NK cells built on key markers (CD62L, CD44, CD103, CD69, CD25, PD-1, Nrp1, ICOS, KLRG1, ST2, Ki67, CTLA4). Colors indicate annotated FlowSOM clusters, with quantification and **(H)** frequency of marker expression. **(I)** tSNEs of brain Tregs built on key markers (CD62L, CD44, CD103, CD69, CD25, PD-1, Nrp1, ICOS, KLRG1, ST2, Ki67, CTLA4). Colors indicate annotated FlowSOM clusters, with quantification and **(J)** frequency of marker expression. Mean ± SEM.

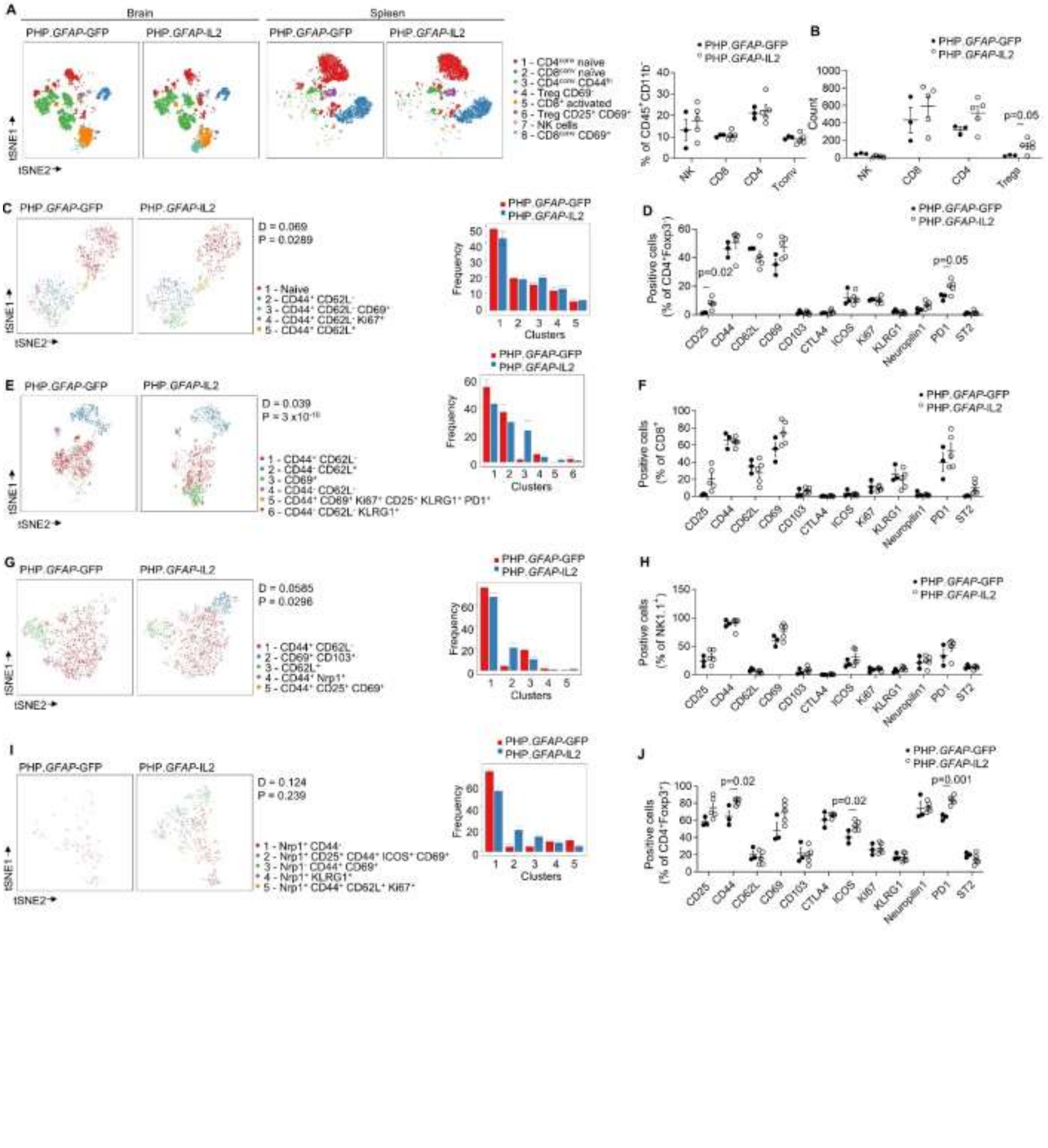
A
Brain
Spleen
PHP.GFAP-GFP
PHP.GFAP-IL2
PHP.GFAP-GFP
PHP.GFAP-IL2
1 - CD4conv naive
2 - CD8conv naive
3 - CD4conv CD44hi
4 - Treg CD69-
5 - CD8+ activated
6 - Treg CD25+ CD69+
7 - NK cells
8 - CD8conv CD69+
tSNE1
tSNE2
PHP.GFAP-GFP
PHP.GFAP-IL2
% of CD45+ CD11b-
NK
CD8
CD4
Tconv
B
PHP.GFAP-GFP
PHP.GFAP-IL2
Count
p=0.05
NK
CD8
CD4
Tregs
C
PHP.GFAP-GFP
PHP.GFAP-IL2
D = 0.069
P = 0.0289
1 - Naive
2 - CD44+ CD62L-
3 - CD44+ CD62L- CD69+
4 - CD44+ CD62L- Ki67+
5 - CD44+ CD62L+
Frequency
Clusters
D
Positive cells (% of CD4+Foxp3+)
p=0.02
p=0.05
CD25
CD44
CD62L
CD69
CD103
CTLA4
ICOS
Ki67
KLRG1
Neuropilin1
PD1
ST2
E
PHP.GFAP-GFP
PHP.GFAP-IL2
D = 0.039
P = 3 x10-10
1 - CD44+ CD62L-
2 - CD44- CD62L+
3 - CD69+
4 - CD44- CD62L-
5 - CD44+ CD69+ Ki67+ CD25+ KLRG1+ PD1+
6 - CD44- CD62L- KLRG1+
F
Positive cells (% of CD8+)
G
PHP.GFAP-GFP
PHP.GFAP-IL2
D = 0.0585
P = 0.0296
1 - CD44+ CD62L-
2 - CD69+ CD103+
3 - CD62L+
4 - CD44+ Nrp1+
5 - CD44+ CD25+ CD69+
H
Positive cells (% of NK1.1+)
I
PHP.GFAP-GFP
PHP.GFAP-IL2
D = 0.124
P = 0.239
1 - Nrp1+ CD44-
2 - Nrp1+ CD25+ CD44+ ICOS+ CD69+
3 - Nrp1- CD44+ CD69+
4 - Nrp1+ KLRG1+
5 - Nrp1+ CD44+ CD62L+ Ki67+
J
Positive cells (% of CD4+Foxp3+)
p=0.02
p=0.02
p=0.001

**Supplementary Figure 16. Elevated IL2 did not produce detectable effects on astrocyte or neuron function.** Acute brain slices were prepared from either αCamKII$^{IL2}$ or PHP.*GFAP*-IL2 treated mice. $Ca^{2+}$ imaging in astrocytes was performed using Fluo4. **(A)** (Left, red) Representative image of $SR101^{+}$ astrocytes from the visual cortex. (Right) Pseudo-color images taken from the Fluo4 channel at baseline and following application of the α1-noradrenergic receptor agonist phenylephrine (PHE) to induce $Ca^{2+}$ release in astrocytes. Pseudo-color images were obtained by summing the florescence intensities measured in each region of interest (dotted circles) for each frame of a 10 s recording period (centered on the indicated timepoints). **(B)** Averaged ΔF/F0 traces (solid line) and corresponding SEM (shaded area) for $Ca^{2+}$ levels in astrocytes from controls (wildtype littermates) or αCamKII$^{IL2}$ mice. **(C)** Boxplots of the ΔF/F0 amplitude (left) and Area Under the Curve (AUC) (right) for both groups. n = 3 mice in each group and $n_{astrocytes}$ = 710 and 774 for control and αCamKII$^{IL2}$ groups, respectively. Whiskers of the boxplots represent the maximal and minimal values. **(D)** Averaged ΔF/F0 traces (solid line) and corresponding SEM (shaded area) for astrocytes from PHP.*GFAP*-GFP (control) or PHP.*GFAP*-IL2 treated mice. **(E)** Boxplots of the ΔF/F0 amplitude (left) and AUC (right) for both groups. n = 3 mice in each group, and $n_{astrocytes}$ = 609 and 646 for the PHP.*GFAP*-GFP and PHP.*GFAP*-IL2 groups, respectively. Whiskers of the boxplots represent the maximal and minimal values. **(F)** Neuronal function was measured using field excitatory post-synaptic potentials (fEPSPs) in PHP.*GFAP*-GFP and PHP.*GFAP*-IL2 treated mice. Input-output curves were recorded for each slice by applying single-stimuli ranging from 500 to 2750 mV with 250 mV increments. Slope and **(G)** amplitude were analyzed (n=4,4). Long-term potentiation (LTP) was induced by applying three high frequency stimulus trains (theta-burst stimulation (TBS): 100 stimuli; 100 Hz) with 5 minutes intervals between trains. After baseline determination, fEPSPs were measured for 55 minutes. Changes in the **(H)** slope and **(I)** amplitude were analyzed across time. Boxplots represent quantification of the baseline (left) and final LTP (right). Mean ± SEM (n=4,4). Whiskers of the boxplots represent the maximal and minimal values.

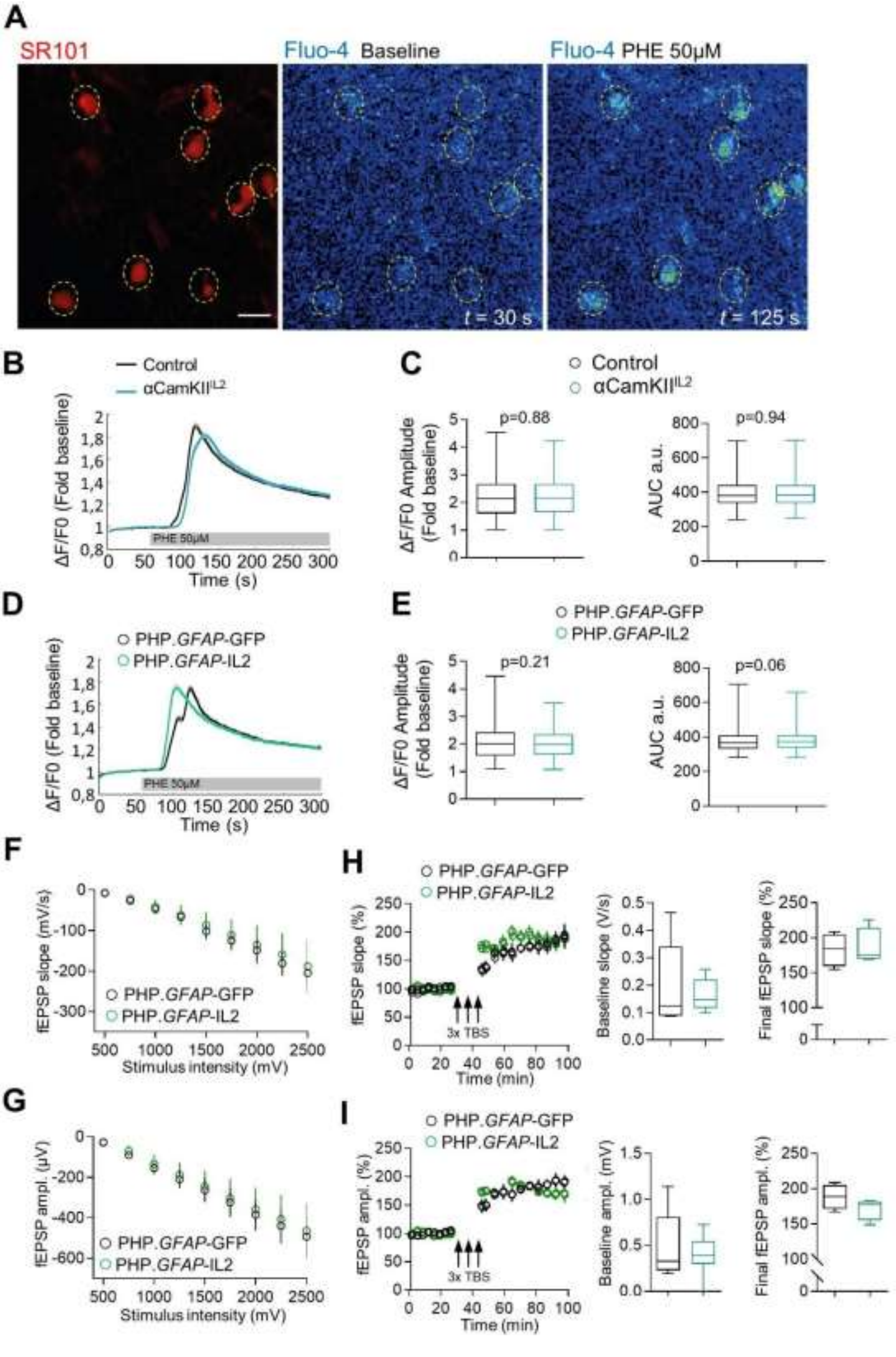
A
SR101
Fluo-4 Baseline
Fluo-4 PHE 50µM
t = 30 s
t = 125 s
B
Control
αCamKII[IL2]
ΔF/F0 (Fold baseline)
PHE 50µM
Time (s)
C
Control
αCamKII[IL2]
ΔF/F0 Amplitude (Fold baseline)
p=0.88
AUC a.u.
p=0.94
D
PHP.GFAP-GFP
PHP.GFAP-IL2
ΔF/F0 (Fold baseline)
PHE 50µM
Time (s)
E
PHP.GFAP-GFP
PHP.GFAP-IL2
ΔF/F0 Amplitude (Fold baseline)
p=0.21
AUC a.u.
p=0.06
F
fEPSP slope (mV/s)
PHP.GFAP-GFP
PHP.GFAP-IL2
Stimulus intensity (mV)
G
fEPSP ampl. (µV)
PHP.GFAP-GFP
PHP.GFAP-IL2
Stimulus intensity (mV)
H
PHP.GFAP-GFP
PHP.GFAP-IL2
fEPSP slope (%)
3x TBS
Time (min)
Baseline slope (V/s)
Final fEPSP slope (%)
I
PHP.GFAP-GFP
PHP.GFAP-IL2
fEPSP ampl. (%)
3x TBS
Time (min)
Baseline ampl. (mV)
Final fEPSP ampl. (%)

**Supplementary Figure 17. Normal behavior in PHP.*GFAP*-IL2-treated mice.** Behavioral assessment of wildtype mice treated with PHP.*GFAP*-GFP or PHP.*GFAP*-IL2 **(A)** Time spent on the rod, average of 4 repeated tests of 300 seconds (n=15,15). **(B)** Open field, total distance moved and time spent in the corners (n = 15,15). **(C)** Nesting behavior (n = 15,15). **(D)** Spatial learning in the Morris Water Maze. Path length to finding the hidden platform (n=15,15). **(E)** Probe tests after 5 days and 10 days and after reversal learning (n=15,15). **(F)** Freezing behavior over time during context acquisition conditioning (n=15,15). **(G)** Contextual discrimination during generalization test (n=15,15). **(H)** Sociability test trials to monitor the interaction with a stranger mouse (S1) compared to an empty chamber (E) (n=15,15). Mean ± SEM.

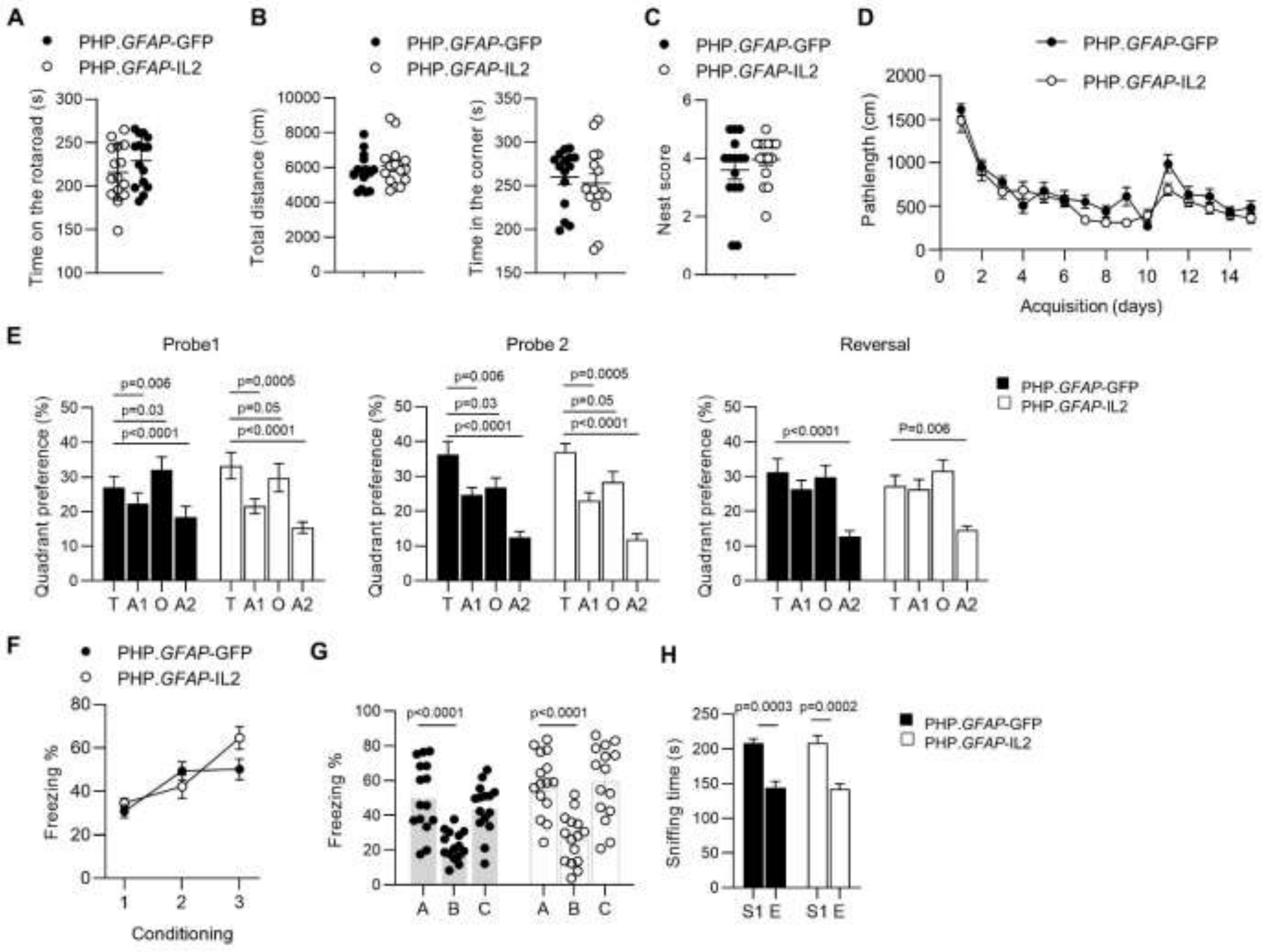

**Supplementary Figure 18. Intact blood-brain barrier integrity following PHP treatment.** Wildtype mice treated with PBS, PHP.*GFAP*-GFP or PHP.*GFAP*-IL2 were assessed at post-injection day 14 for blood-brain barrier (BBB) integrity. **(A)** Histological assessment for CD31, ZO-1 and DAPI or **(B)** CD31, OCLN and DAPI. Scale bar, 10 µm. **(C)** Histological assessment for CLDN1 and DAPI or E-cadherin/CDH1 and DAPI. Scale bar, 50 µm. Insert scale bar, 10 µm. **(D)** Mice were injected i.v. with 4 kDa FITC-dextran, followed by quantification in the cerebral spinal fluid (CSF), cerebellum, cortex and hippocampus (n=4-6 mice/group). Mean ± SEM.

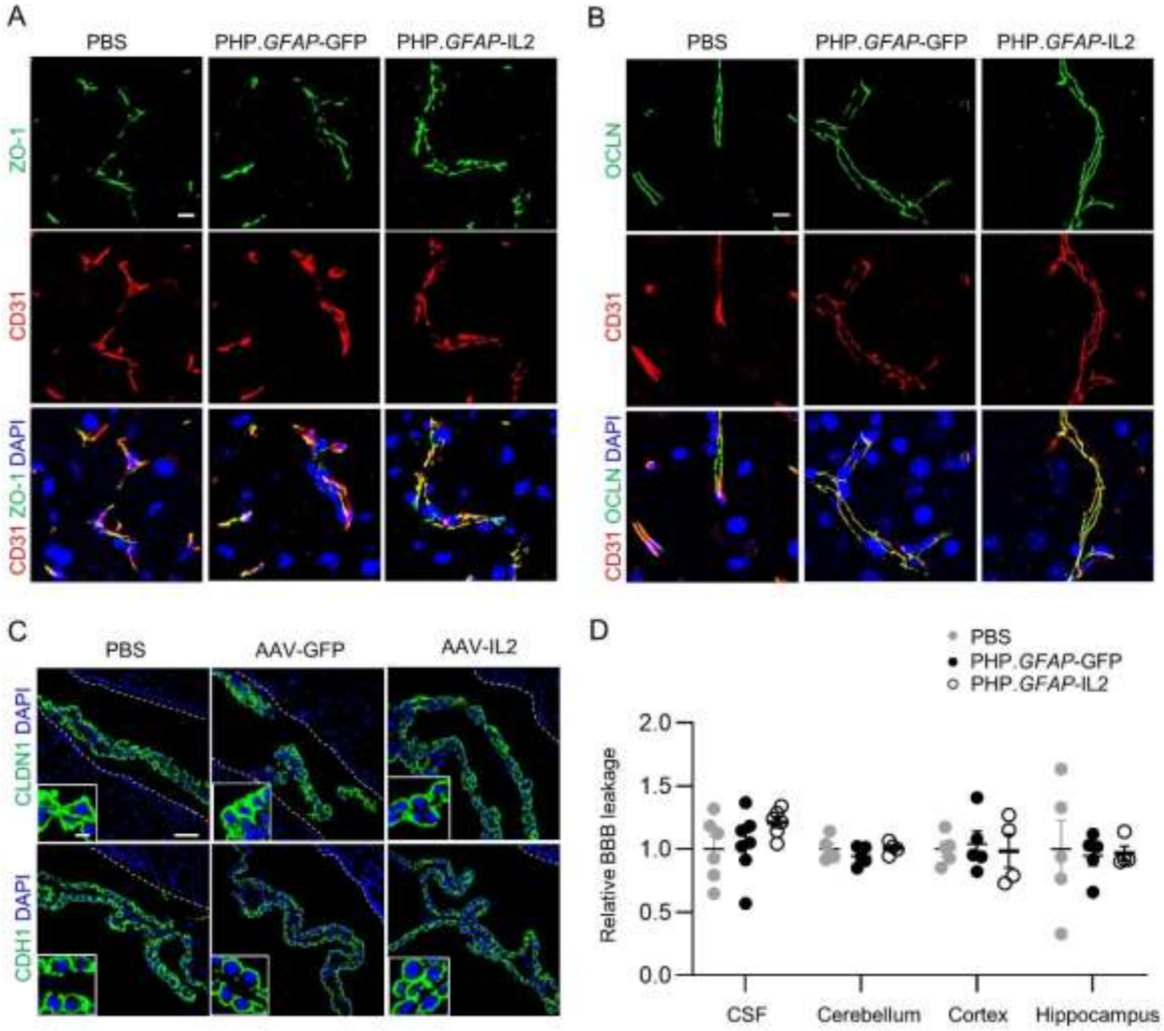

**Supplementary Figure 19. Brain-specific Treg expansion following PHP.*CamKII*-IL2 treatment. (A)** The *CamKII* promoter restricts gene expression (as assessed using GFP scoring) to neurons (NeuN positive) in adult mouse brain. Off-target expression was not detected when slices were counter-stained for GFAP (astrocytes). Left panel, hippocampus; right panel, cortex. Scale bar, 20 μm. Data are representative images seen in 2 slices from each of 4 independent mice receiving a PHP.CamKII-GFP (control) vector. **(B)** Quantification of GFP colocalization with NeuN and GFAP in PHP.*CamKII*-GFP-treated mice. **(C)** Levels of IL2 were measured from tissue samples obtained from WT mice administered with $1x10^9$, $1x10^{10}$ or $1x10^{11}$ total vector genomes of PHP.*CamKII*-GFP (control) or PHP.CamKII-IL2 (n=7,7,5,8,8,8). **(D)** Wildtype mice were administered, intravenously, $1x10^9$, $1x10^{10}$ or $1x10^{11}$ vector genomes (total dose) of PHP.*CamKII*-GFP or PHP.*CamKII*-IL2 and assessed for the number of conventional T cells (left) and Tregs (right), 14 days after treatment (n=5 group) in the perfused brain or **(E)** spleen. **(F)** Tregs as a percentage of CD4 T cells in the brain and **(G)** spleen.

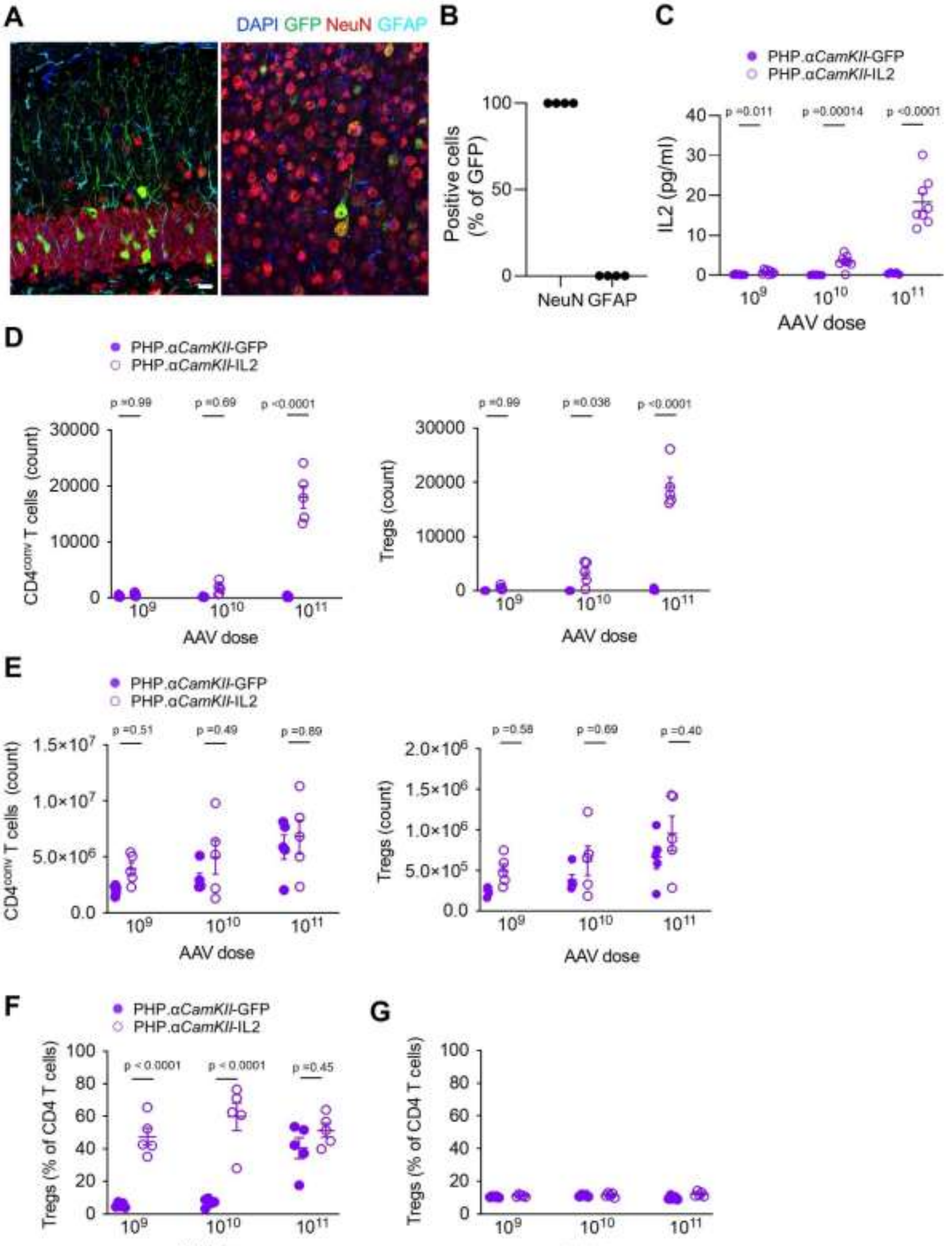

A
DAPI GFP NeuN GFAP
B
Positive cells (% of GFP)
100
50
0
NeuN GFAP
C
PHP.αCamKII-GFP
PHP.αCamKII-IL2
IL2 (pg/ml)
p =0.011
p =0.00014
p <0.0001
AAV dose
D
PHP.αCamKII-GFP
PHP.αCamKII-IL2
CD4conv T cells (count)
p =0.99
p =0.69
p <0.0001
Tregs (count)
p =0.99
p =0.036
p <0.0001
AAV dose
E
PHP.αCamKII-GFP
PHP.αCamKII-IL2
CD4conv T cells (count)
p =0.51
p =0.49
p =0.89
Tregs (count)
p =0.58
p =0.69
p =0.40
AAV dose
F
PHP.αCamKII-GFP
PHP.αCamKII-IL2
Tregs (% of CD4 T cells)
p < 0.0001
p < 0.0001
p =0.45
AAV dose
G
Tregs (% of CD4 T cells)
AAV dose

**Supplementary Figure 20. Normal peripheral influx following PHP.*GFAP*-IL2 treatment in traumatic brain injury mice.** Mice treated with PHP.*GFAP*-GFP control or PHP.*GFAP*-IL2 (day -14) were given controlled cortical impacts to induce moderate TBI and examined at 15 days post-TBI (n = 3, 4, 4); a sham TBI was included in the PHP.*GFAP*-GFP group. Perfused brains from sham, TBI and PHP.*GFAP*-IL2-treated TBI mice were compared by high-dimensional flow cytometry. **(A)** Absolute number of microglia, gated as $CD11b^{+}$ $CX3CR1^{+}$ $CD64^{+}$ $CD45^{mod}$ $Ly6G^{-}$ cells. **(B)** CD8 and CD4 T cells, as a proportion of $CD45^{+}CD11b^{-}$ cells. **(C)** Absolute number of Tregs, CD4 Tconv and CD8 T cells. **(D)** Frequency of CD25, CD44, CD69, Ki67 and PDL1 expressing-cells, and **(E)** frequency or **(F)** mean fluorescence intensity (MFI) of amphiregulin (Areg)-producing cells within the CD4 conventional T cell population. **(G)** Frequency of CD25, CD44, CD69, Ki67 and PDL1 expressing-cells, and **(H)** frequency of IL-10 or Areg-producing cells, within the CD4 conventional T cell population. **(I)** Mean expression of Areg, in Areg-producing CD4 conventional T cells. Mean ± SEM. **(J)** Representative histograms for CD25, CD44, CD69, Ki67, PDL1, IL-10 and Areg in CD4 conventional T cells, or **(K)** Tregs, from sham, TBI and PHP.*GFAP*-IL2-treated TBI mice. **(L)** Mice treated with PHP.*GFAP*-GFP control or PHP.*GFAP*-IL2 (day -14) were given controlled cortical impacts to induce moderate TBI and examined at 15 days post-TBI (n = 5/group). Superficial cervical lymph nodes were assessed for the percentage of CD4, CD8 and γδ T cells within the T cell compartment, and **(M)** Tregs within the CD4 T cell compartment, **(N)** with calculation of absolute numbers. **(O)** Cervical lymph nodes were assessed for the percentage of CD4, CD8 and γδ T cells within the T cell compartment, and **(P)** Tregs within the CD4 T cell compartment, **(Q)** with calculation of absolute numbers.

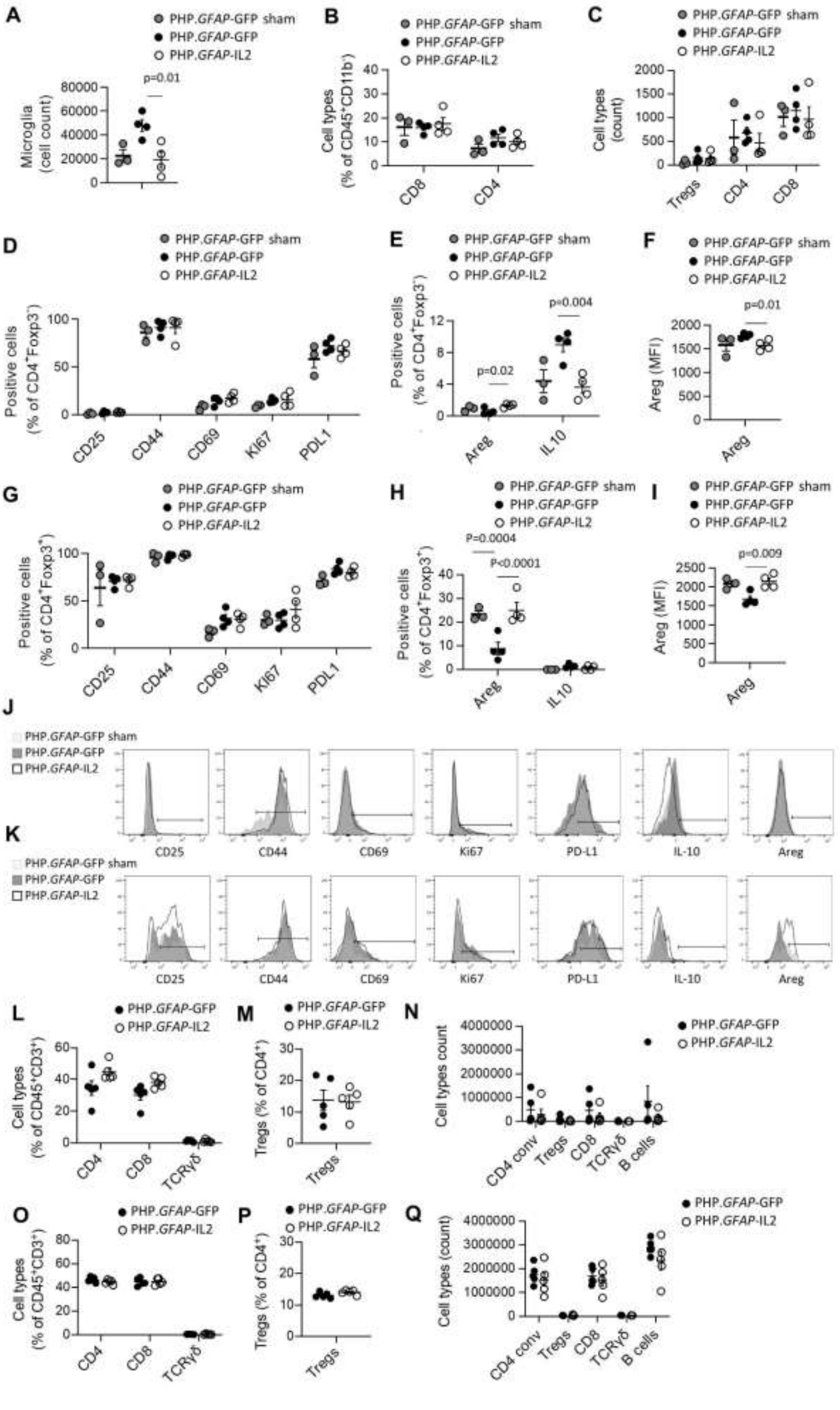
A
PHP.GFAP-GFP sham
PHP.GFAP-GFP
PHP.GFAP-IL2
p=0.01
Microglia (cell count)
80000
60000
40000
20000
0
B
Cell types (% of CD45+CD11b-)
CD8
CD4
C
Cell types (count)
Tregs
CD4
CD8
D
Positive cells (% of CD4+Foxp3-)
CD25
CD44
CD69
Ki67
PDL1
E
Positive cells (% of CD4+Foxp3-)
p=0.02
p=0.004
Areg
IL10
F
Areg (MFI)
p=0.01
Areg
G
Positive cells (% of CD4+Foxp3+)
CD25
CD44
CD69
Ki67
PDL1
H
P=0.0004
P<0.0001
Positive cells (% of CD4+Foxp3+)
Areg
IL10
I
p=0.009
Areg (MFI)
Areg
J
PHP.GFAP-GFP sham
PHP.GFAP-GFP
PHP.GFAP-IL2
CD25
CD44
CD69
Ki67
PD-L1
IL-10
Areg
K
PHP.GFAP-GFP sham
PHP.GFAP-GFP
PHP.GFAP-IL2
CD25
CD44
CD69
Ki67
PD-L1
IL-10
Areg
L
PHP.GFAP-GFP
PHP.GFAP-IL2
Cell types (% of CD45+CD3+)
CD4
CD8
TCRγδ
M
Tregs (% of CD4+)
Tregs
N
Cell types count
4000000
3000000
2000000
1000000
0
CD4 conv
Tregs
CD8
TCRγδ
B cells
O
Cell types (% of CD45+CD3+)
CD4
CD8
TCRγδ
P
Tregs (% of CD4+)
Tregs
Q
Cell types (count)
CD4 conv
Tregs
CD8
TCRγδ
B cells

**Supplementary Figure 21. Transcriptional analysis following brain-specific delivery of IL2 during TBI.** Wildtype mice, treated with PHP.*GFAP*-IL2 (or PHP.*GFAP*-GFP control vector) on day -14 were given controlled cortical impacts to induce moderate TBI or sham surgery. 14 days post-TBI, T cells and microglia were sorted from the perfused brains for 10x single-cell transcriptomics. **(A)** UMAP expression plot of T cell data, with expression patterns of *CD3d, CD4, CD8, Foxp3, IL2RA* and *Sel1* superimposed to identify various T cell populations. **(B)** T cell UMAP representation, showing the relative numbers of various T cell types across treatment groups. **(C)** Volcano plot showing differential gene expression in the CD4 Tconv cluster between PHP.*GFAP*-GFP- and PHP.*GFAP*-IL2-treated mice, for sham (left) and TBI (right). **(D)** Volcano plot showing differential gene expression in the CD8 T cell cluster between PHP.*GFAP*-GFP- and PHP.*GFAP*-IL2-treated mice, for sham (left) and TBI (right). **(E)** UMAP expression plot of microglia data, with differential expression patterns of *Lpl, Cst7, Axl, Itgax, Spp1, Ccl6, Csf1* and *H2-Aa* shown. **(F)** Volcano plot showing differential gene expression for total microglia between PHP.*GFAP*-GFP- and PHP.*GFAP*-IL2-treated mice, for sham (left) and TBI (right). **(G)** Volcano plot showing differential gene expression for homeostatic microglia between PHP.*GFAP*-GFP- and PHP.*GFAP*-IL2-treated mice, for sham (left) and TBI (right). **(H)** Filtered overview of KEGG pathways based on differential gene enrichment in microglia from PHP.*GFAP*-GFP- and PHP.*GFAP*-IL2-treated mice, for sham and TBI conditions. **(I)** Pathview plot for “Antigen processing and presentation” (KEGG mmu04612), using the average log-fold changes between the four differential gene expression comparisons indicated.

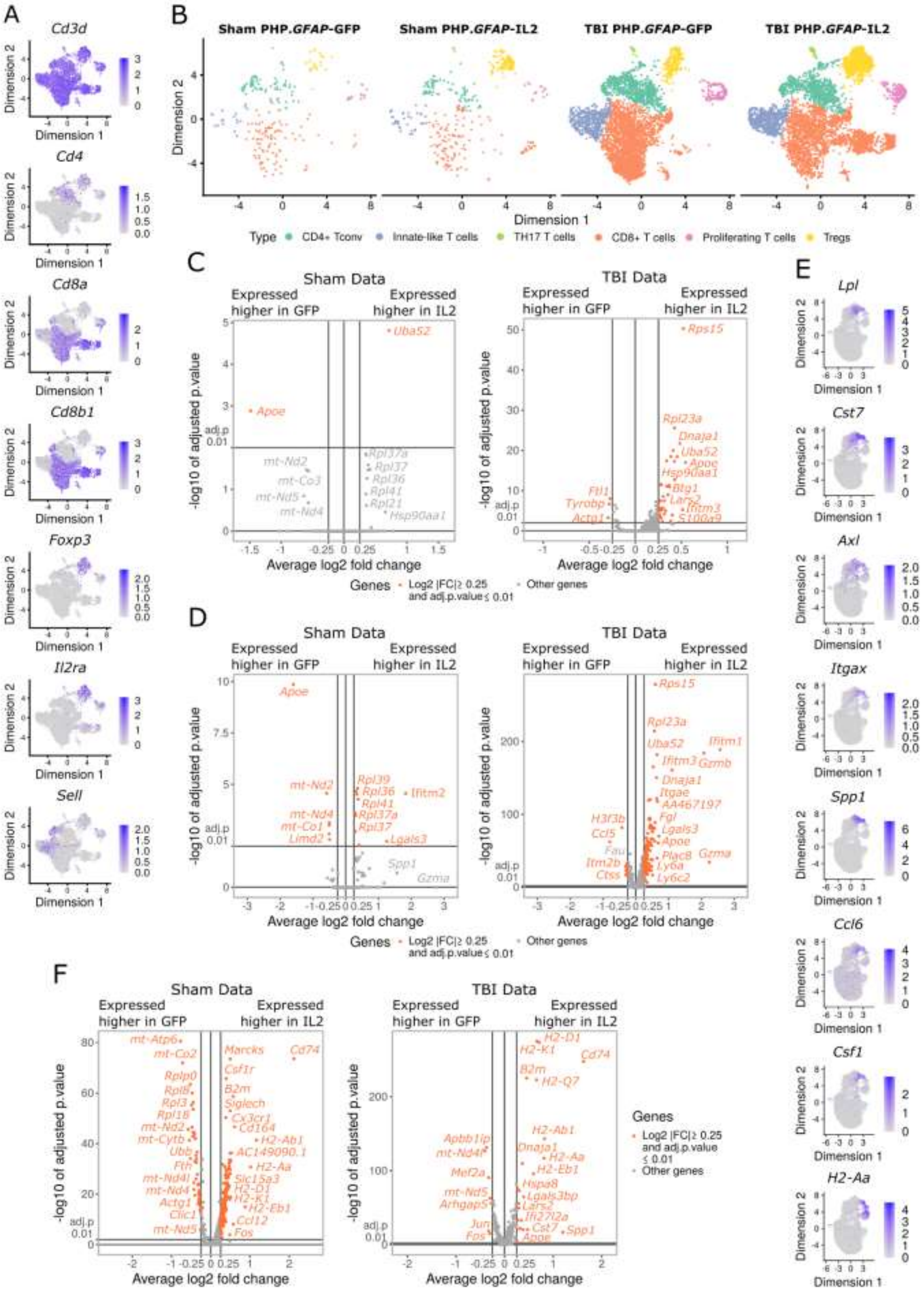
A
Cd3d
Cd4
Cd8a
Cd8b1
Foxp3
Il2ra
Sell
Dimension 1
Dimension 2
B
Sham PHP.GFAP-GFP
Sham PHP.GFAP-IL2
TBI PHP.GFAP-GFP
TBI PHP.GFAP-IL2
Type
CD4+ Tconv
Innate-like T cells
TH17 T cells
CD8+ T cells
Proliferating T cells
Tregs
C
Sham Data
TBI Data
Expressed higher in GFP
Expressed higher in IL2
-log10 of adjusted p.value
Average log2 fold change
adj.p 0.01
Uba52
Apoe
mt-Nd2
mt-Co3
mt-Nd5
mt-Nd4
Rpl37a
Rpl37
Rpl36
Rpl41
Rpl21
Hsp90aa1
Rps15
Rpl23a
Dnaja1
Uba52
Apoe
Hsp90aa1
Btg1
Lars2
Ifitm3
S100a9
Ftl1
Tyrobp
Actg1
Genes
Log2 |FC|≥ 0.25 and adj.p.value ≤ 0.01
Other genes
D
Sham Data
TBI Data
Apoe
mt-Nd2
mt-Nd4
mt-Co1
Limd2
Rpl39
Rpl36
Rpl41
Rpl37a
Rpl37
Ifitm2
Lgals3
Spp1
Gzma
Rps15
Rpl23a
Uba52
Ifitm1
Ifitm3
Gzmb
Dnaja1
Itgae
AA467197
Fgl
Lgals3
Apoe
Plac8
Ly6a
Ly6c2
Gzma
H3f3b
Ccl5
Fau
Itm2b
Ctss
E
Lpl
Cst7
Axl
Itgax
Spp1
Ccl6
Csf1
H2-Aa
F
Sham Data
TBI Data
mt-Atp6
mt-Co2
Rplp0
Rpl8
Rpl3
Rpl18
mt-Nd2
mt-Cytb
Ubb
Fth
mt-Nd4l
mt-Nd4
Actg1
Clic1
mt-Nd5
Marcks
Cd74
Csf1r
B2m
Siglech
Cx3cr1
Cd164
H2-Ab1
AC149090.1
H2-Aa
Slc15a3
H2-D1
H2-K1
H2-Eb1
Ccl12
Fos
H2-D1
H2-K1
Cd74
B2m
H2-Q7
H2-Ab1
Dnaja1
H2-Aa
H2-Eb1
Hspa8
Lgals3bp
Lars2
Ifi27l2a
Cst7
Spp1
Apoe
Apbb1ip
mt-Nd4l
Mef2a
mt-Nd5
Arhgap5
Jun
Fos
Genes
Log2 |FC|≥ 0.25 and adj.p.value ≤ 0.01
Other genes

G

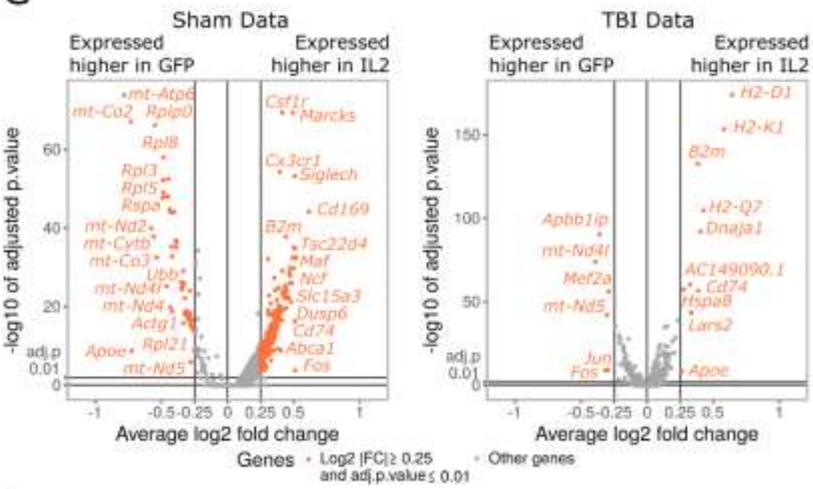
Sham Data
Expressed higher in GFP
Expressed higher in IL2
TBI Data
-log10 of adjusted p.value
Average log2 fold change
adj.p 0.01
mt-Atp6
mt-Co2
Rplp0
Rpl8
Rpl3
Rpl5
Rpsa
mt-Nd2
mt-Cytb
mt-Co3
Ubb
mt-Nd4l
mt-Nd4
Actg1
Apoe
Rpl21
mt-Nd5
Csf1r
Marcks
Cx3cr1
Siglech
Cd169
B2m
Tsc22d4
Maf
Ncf
Slc15a3
Dusp6
Cd74
Abca1
Fos
H2-D1
H2-K1
H2-Q7
Dnaja1
Apbb1ip
mt-Nd4l
Mef2a
mt-Nd5
AC149090.1
Hspa8
Lars2
Jun
Apoe
Genes
Log2 |FC| ≥ 0.25 and adj.p.value ≤ 0.01
Other genes

H

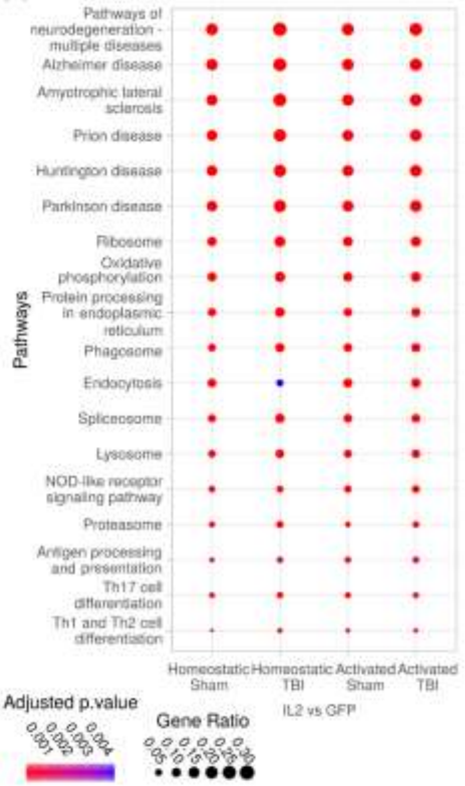
Pathways
Pathways of neurodegeneration - multiple diseases
Alzheimer disease
Amyotrophic lateral sclerosis
Prion disease
Huntington disease
Parkinson disease
Ribosome
Oxidative phosphorylation
Protein processing in endoplasmic reticulum
Phagosome
Endocytosis
Spliceosome
Lysosome
NOD-like receptor signaling pathway
Proteasome
Antigen processing and presentation
Th17 cell differentiation
Th1 and Th2 cell differentiation
Homeostatic Sham
Homeostatic TBI
Activated Sham
Activated TBI
IL2 vs GFP
Adjusted p.value
0.001
0.002
0.003
0.004
Gene Ratio
0.05
0.10
0.15
0.20
0.25
0.30

I

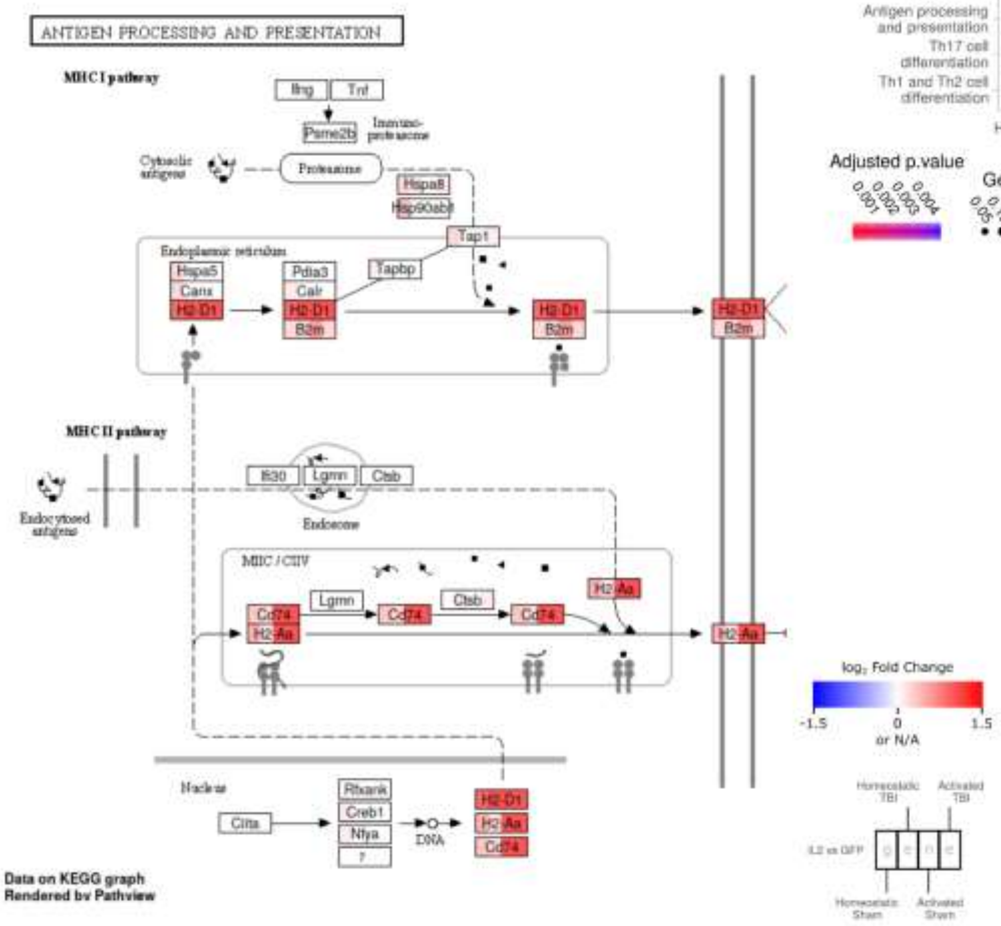
ANTIGEN PROCESSING AND PRESENTATION
MHC I pathway
Ifng
Tnf
Psme2b
Immuno-proteasome
Cytosolic antigens
Proteasome
Hspa8
Hsp90ab1
Tap1
Endoplasmic reticulum
Hspa5
Canx
H2-D1
Pdia3
Calr
B2m
Tapbp
MHC II pathway
Endocytosed antigens
Ifi30
Lgmn
Ctsb
Endosome
MIIC / CIIV
H2-Aa
Cd74
Nucleus
Ciita
Rfxank
Creb1
Nfya
DNA
log2 Fold Change
-1.5
0
1.5
or N/A
Homeostatic TBI
Activated TBI
IL2 vs GFP
Homeostatic Sham
Activated Sham
Data on KEGG graph
Rendered by Pathview

**Supplementary Figure 22. Normal peripheral influx during distal middle cerebral artery occlusion following PHP.GFAP-IL2 treatment.** Mice treated with PHP.*GFAP*-GFP control or PHP.*GFAP*-IL2 (day -14) were subject to dMCAO and examined at day 14 (n = 3, 5, 2). Perfused brains were compared by high-dimensional flow cytometry. **(A)** Absolute number of microglia, gated as $CD11b^+$ $CX3CR1^+$ $CD64^+$ $CD45^{mod}$ $Ly6G^-$ cells. **(B)** Frequency of CD4, CD8 and γδ T cells within $CD45^+$ cells, and **(C)** frequency of Tregs within $CD4^+$ T cells. **(D)** Absolute numbers of CD4 Tconv, Tregs CD8, and γδ T cells. **(E)** Expression of CD25, CD44, CD69, Ki67 and PDL1 markers, or **(F)** amphiregulin (Areg), IL10 and IL17 within the CD4 conventional T cell population. **(G)** Expression of CD25, CD44, CD69, Ki67 and PDL1 markers, or **(H)** Areg, IL10 and IL17 within the Treg population. **(I)** Superficial cervical lymph nodes were assessed for the frequency of CD4 Tconv, Tregs, CD8, Treg, B cells and γδ T cells within the $CD45^+$ cell population, with **(J)** absolute numbers. Mean ± SEM.

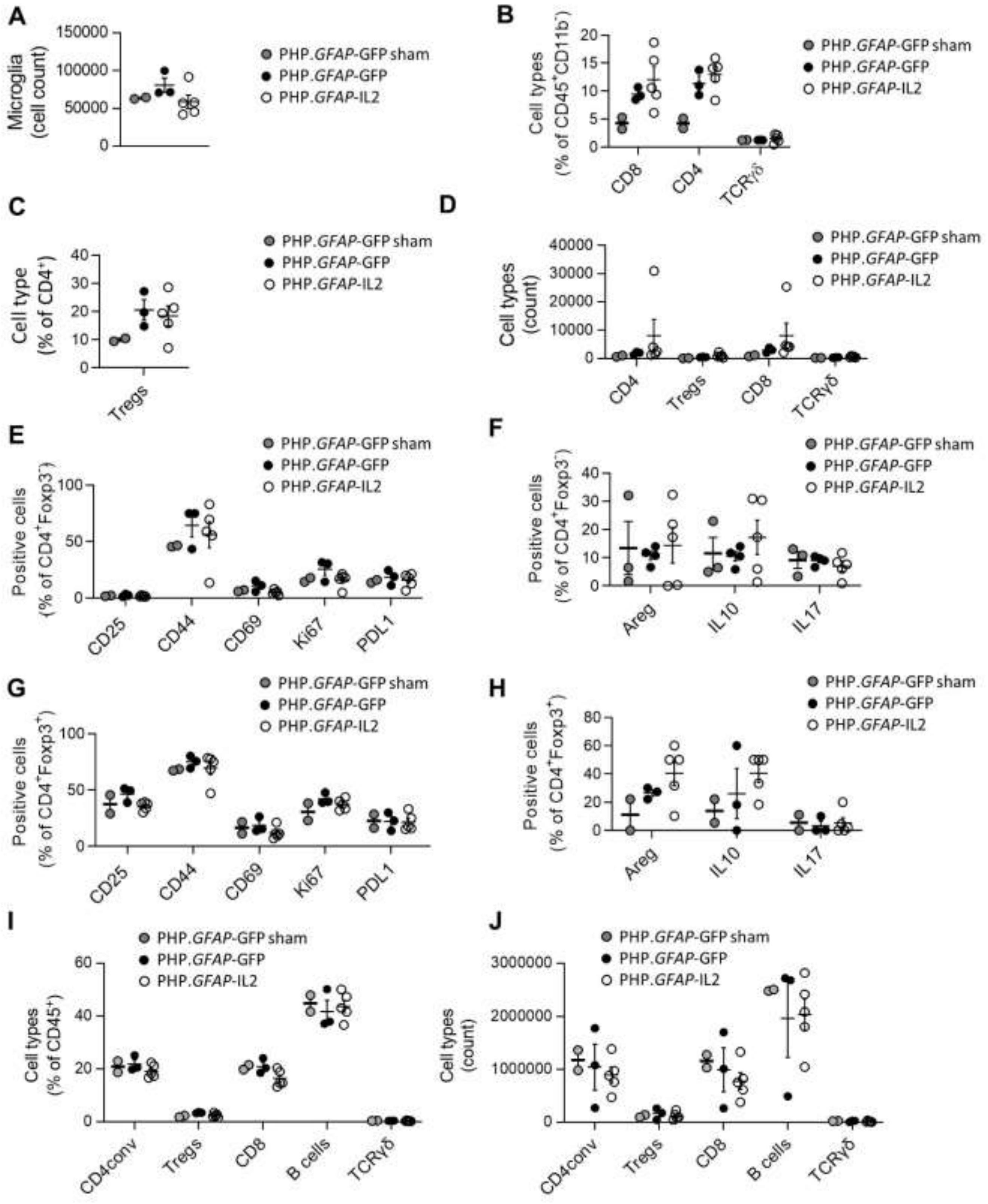
A
Microglia (cell count)
150000
100000
50000
0
PHP.GFAP-GFP sham
PHP.GFAP-GFP
PHP.GFAP-IL2
B
Cell types (% of CD45+CD11b-)
CD8
CD4
TCRγδ
C
Cell type (% of CD4+)
Tregs
D
Cell types (count)
CD4
Tregs
CD8
TCRγδ
E
Positive cells (% of CD4+Foxp3-)
CD25
CD44
CD69
Ki67
PDL1
F
Positive cells (% of CD4+Foxp3-)
Areg
IL10
IL17
G
Positive cells (% of CD4+Foxp3+)
CD25
CD44
CD69
Ki67
PDL1
H
Positive cells (% of CD4+Foxp3+)
Areg
IL10
IL17
I
Cell types (% of CD45+)
CD4conv
Tregs
CD8
B cells
TCRγδ
J
Cell types (count)
CD4conv
Tregs
CD8
B cells
TCRγδ

**Supplementary Figure 23. Normal peripheral influx during photothrombotic stroke following PHP.*GFAP*-IL2 treatment.** Mice treated with PHP.*GFAP*-GFP control or PHP.*GFAP*-IL2 (day -14) were induced with a photothrombotic stroke and examined at day 1 (n = 6, 6, 3). Perfused brains were compared by high-dimensional flow cytometry. **(A)** Absolute number of microglia, gated as $CD11b^+$ $CX3CR1^+$ $CD64^+$ $CD45^{mod}$ $Ly6G^-$ cells. **(B)** Frequency of CD4, CD8 and γδ T cells within $CD45^+$ cells, and **(C)** frequency of Tregs within $CD4^+$ T cells. **(D)** Absolute numbers of CD4 Tconv, Tregs, CD8, and γδ T cells. **(E)** Expression of CD25, CD44, CD69 and Ki67 markers, or **(F)** amphiregulin (Areg), IL10 and IL17 within the CD4 conventional T cell population. **(G)** Expression of CD25, CD44, CD69 and Ki67 markers, or **(H)** Areg, IL10 and IL17 within the Treg population. Mean ± SEM.

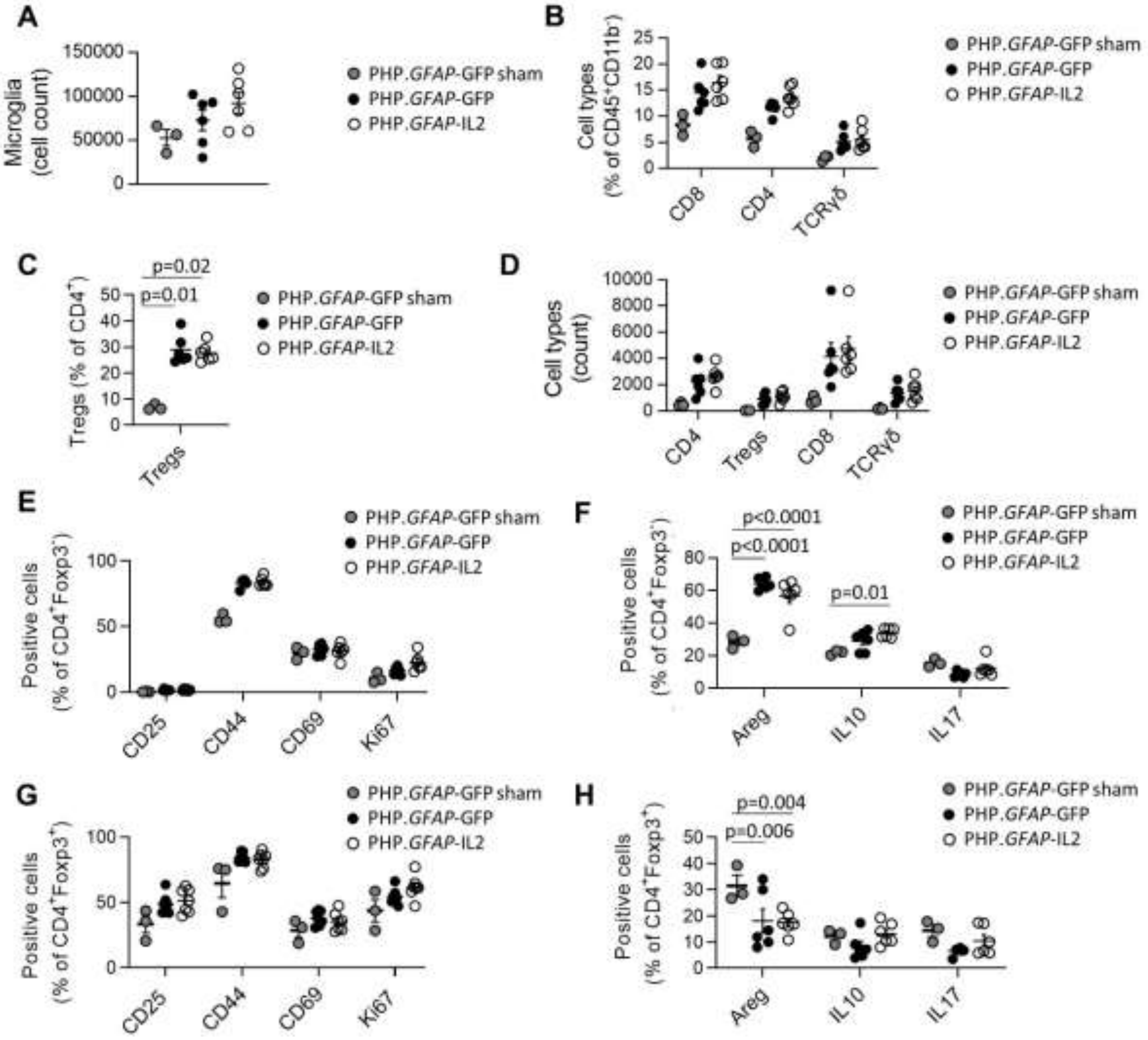

**Supplementary Figure 24. Normal peripheral influx during experimental autoimmune encephalomyelitis following PHP.*GFAP*-IL2 treatment.** Mice treated with PHP.*GFAP*-GFP control or PHP.*GFAP*-IL2 (day -14) were induced with experimental autoimmune encephalomyelitis (EAE) and examined at days 15, 21 and 30 (n = 5-7/group). Perfused CNS was compared by high-dimensional flow cytometry. **(A)** Frequency of CD8 T cells and **(B)** CD4 T cells within $CD45^+$ cells, and **(C)** frequency of Tregs within $CD4^+$ T cells. **(D)** Absolute number of CD8 T cells, **(E)** CD4 T cells and **(F)** Tregs. **(G)** Frequency of CD25, CD44, CD62L, CD69, CD103, CTLA4, GITR, Helios, ICOS, Ki67, KLRG1, Neuropilin1, PD1 and Tbet expression within the CNS-resident Treg population on day 15, **(H)** day 21 or **(I)** day 30. **(J)** IL10 and **(K)** amphiregulin (Areg) expression within CNS-resident Tregs. **(L)** IL17, **(M)** IFNγ, **(N)** GM-CSF, **(O)** TNFα, **(P)** IL2 and **(Q)** IFNγ-TNFα cytokine expression within the CNS-resident Tconv population at various times post-EAE induction. **(R)** 15 days post-induction, major leukocyte subsets were assessed by flow cytometry in the superficial cervical lymph nodes and **(S)** the deep cervical lymph nodes. **(T)** Absolute numbers of major leukocyte subsets in the superficial and **(U)** deep cervical lymph nodes (n=5,5). Mean ± SEM.

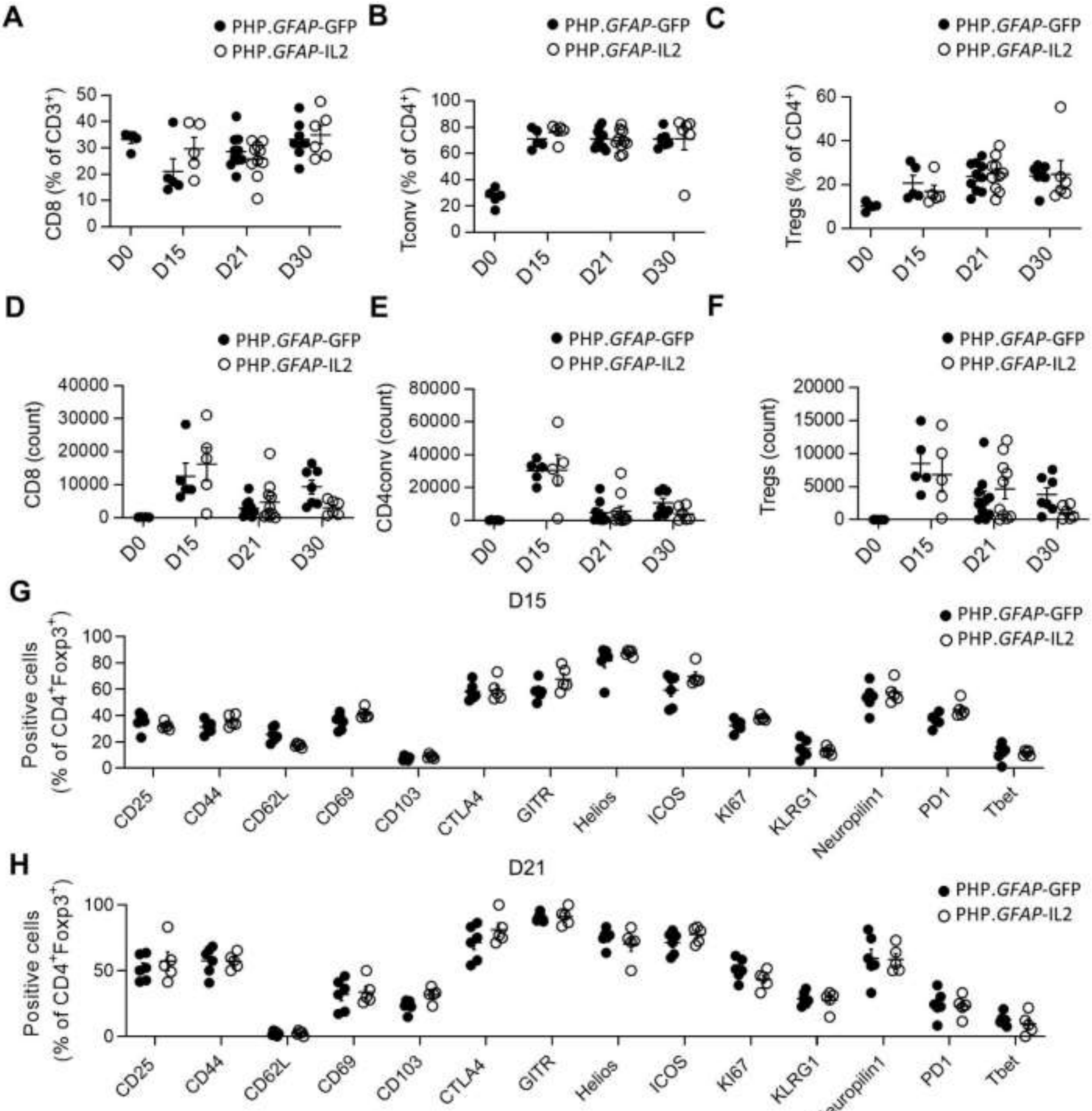

A
PHP.GFAP-GFP
PHP.GFAP-IL2
CD8 (% of CD3+)
D0
D15
D21
D30
B
Tconv (% of CD4+)
C
Tregs (% of CD4+)
D
CD8 (count)
E
CD4conv (count)
F
Tregs (count)
G
D15
Positive cells
(% of CD4+Foxp3+)
CD25
CD44
CD62L
CD69
CD103
CTLA4
GITR
Helios
ICOS
KI67
KLRG1
Neuropilin1
PD1
Tbet
H
D21

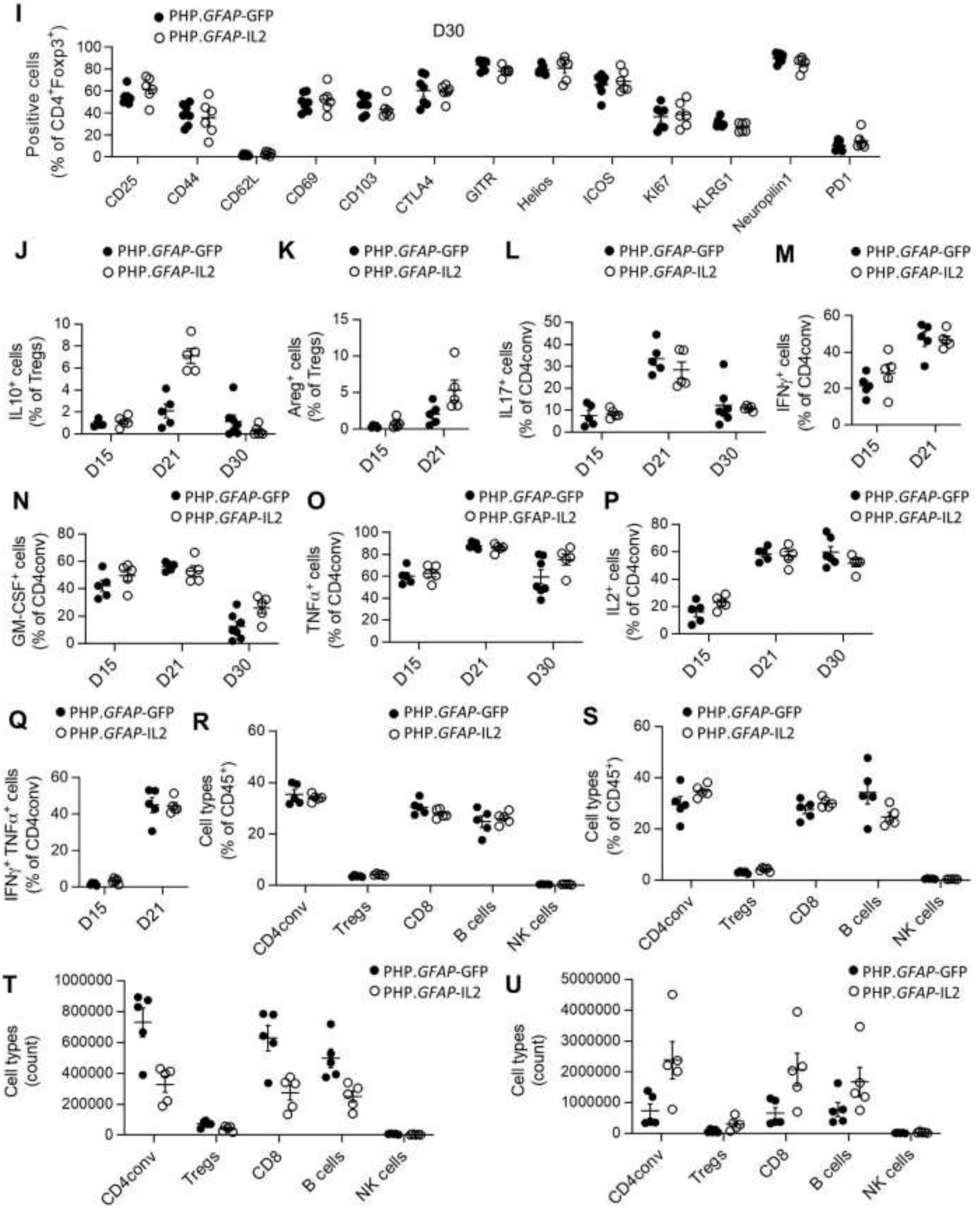
I
PHP.GFAP-GFP
PHP.GFAP-IL2
D30
Positive cells (% of CD4+Foxp3+)
CD25
CD44
CD62L
CD69
CD103
CTLA4
GITR
Helios
ICOS
KI67
KLRG1
Neuropilin1
PD1
J
IL10+ cells (% of Tregs)
K
Areg+ cells (% of Tregs)
L
IL17+ cells (% of CD4conv)
M
IFNγ+ cells (% of CD4conv)
D15
D21
D30
N
GM-CSF+ cells (% of CD4conv)
O
PHP.GFAP-IL2
TNFα+ cells (% of CD4conv)
P
IL2+ cells (% of CD4conv)
Q
IFNγ+ TNFα+ cells (% of CD4conv)
R
S
Cell types (% of CD45+)
CD4conv
Tregs
CD8
B cells
NK cells
T
U
Cell types (count)

**Supplementary Figure 25. Gating strategy for analyzing Tregs cells from the brain.** Representative gating strategy used to quantify Treg numbers in the brain using flow cytometry.

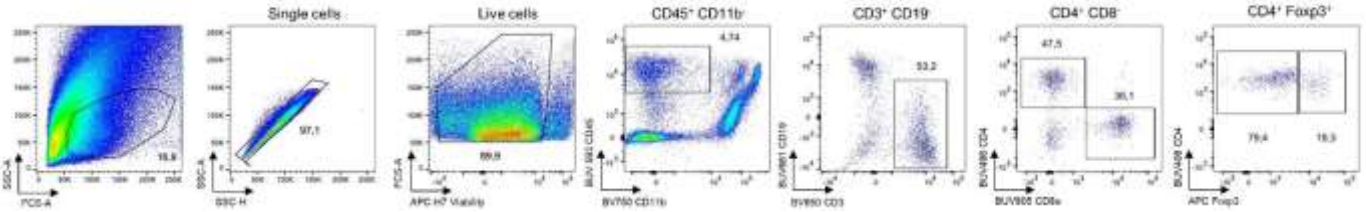
